# Horizontally acquired *IbACS* gene modulates rhizosphere microbiota and contributes to sweet potato growth

**DOI:** 10.64898/2025.12.18.694140

**Authors:** Aiko Tanaka, Motoyasu Otani, Osamu Nakayachi, Minenosuke Matsutani, Hiromasa Saitoh, Daigo Takemoto

## Abstract

*Ipomoea batatas* (sweet potato) exhibits tolerance to nutrient-poor environments and has historically been utilized as a staple crop during periods of food scarcity. Previously, genomic studies have identified *Agrobacterium*-derived sequences in the sweet potato genome. Among these, the *IbACS* (Agrocinopine Synthase) gene encodes an enzyme responsible for the biosynthesis of agrocinopine A, a sugar-phosphodiester opine that can be metabolized and utilized by specific microorganisms harboring the corresponding catabolic enzymes. Because *IbACS* has been conserved in cultivated sweet potato varieties, it is suggested that endogenous agrocinopine production by sweet potato contributes to interactions with environmental microbes, thereby supporting plant growth under nutrient-limited conditions. To test this hypothesis, we generated *IbACS*-disrupted lines using genome editing. Growth of the knockout plants was markedly reduced in non-sterile soil, while no difference was observed in sterile conditions. These results suggest that the role of *IbACS* is linked to interactions with the microbial community rather than to intrinsic plant development. Metagenomic analyses of rhizosphere communities revealed that *IbACS* disruption caused extensive remodeling of microbial composition, including strong reductions in Proteobacteria taxa such as Oxalobacteraceae and decreases in key genera like *Massilia* and *Ensifer*, along with lower community evenness and the expansion of opportunistic minor taxa. Together, these microbial shifts indicate that *IbACS* is required for the assembly of a functionally supportive rhizosphere microbiome that promotes sweet potato growth in soil environments.

## INTRODUCTION

Horizontal gene transfer (HGT), the acquisition of genetic material from unrelated organisms, is a well-documented mechanism of adaptation in microorganisms, enabling rapid shifts in metabolic capacity and ecological niche. In contrast, cases of HGT into plants are considered extremely rare, and only a few convincing examples have been reported (Suzuki et al. 2002; Chen et al. 2014; Matveeva and Otten 2019). One of the most notable cases is found in sweet potato (*Ipomoea batatas*), which carries stably integrated T-DNA (cellular T-DNA, cT-DNA) derived from ancestral *Agrobacterium* in its genome. Analysis of 291 cultivated varieties showed that all of them contain such cT-DNA sequences (Kyndt et al. 2015; Yan et al. 2024).

Because cT-DNAs are highly conserved among cultivars, it has been expected that they were not neutral remnants of infection events, but instead conferred traits that were positively selected during domestication and the evolutionary transition from wild relatives to modern cultivars (Yan et al. 2024). Detailed sequence analysis has revealed that one of these cT-DNAs, IbT-DNA1, carries at least four intact genes: *iaaM* (tryptophan monooxygenase) and *iaaH* (indole-acetamide hydrolase), both involved in auxin biosynthesis; *ACS* (agrocinopine synthase), responsible for the synthesis of the opine agrocinopine A; and an additional open reading frame encoding a small protein of unknown function (Kyndt et al. 2015; Tanaka et al. 2022; Yan et al. 2024). These findings indicate that sweet potato has inherited a set of *Agrobacterium* genes related to both plant hormone biosynthesis and opine production, although the biological significance of retaining these genes remains unclear.

Opines such as agrocinopine, octopines, and nopalines are well-studied in the context of *Agrobacterium* biology, where they function not only as nutrient sources but also as selective signals that recruit and enrich microbial populations capable of opine catabolism (Dessaux and Faure 2018; González-Mula et al. 2018). Through this mechanism, opine-producing tumors establish highly specialized microbial niches. These findings raise the possibility that opine production mediated by horizontally acquired genes may influence microbial community assembly beyond the *Agrobacterium*–host interaction system.

Agrocinopine is a unique sugar-phosphodiester compound composed of sucrose and arabinose (or glucose) linked via a phosphodiester bond. Its catabolic enzymes have been identified in *Agrobacterium* and a few other bacterial species (Kim and Farrand 1997; Tanaka et al. 2022), but there has been no evidence that plants themselves can produce this compound. However, we previously demonstrated that the sweet potato *IbACS* gene indeed synthesizes agrocinopine A from endogenous substrates. Moreover, expression of *IbACS* in transgenic tobacco altered the rhizosphere microbial community, including the enrichment of a *Leifsonia* species able to catabolize agrocinopine (Tanaka et al. 2022). These findings indicate that a cT-DNA-derived gene can modulate microbial assemblages in the rhizosphere, supporting the broader concept that HGT may provide plants with novel tools to influence microbe-dependent ecological interactions (Haimlich 2024).

Sweet potato is known for its remarkable ability to grow in nutrient-poor soils, and accumulating evidence suggests that plant performance under such conditions often depends on associations with beneficial microbial communities (Berendsen et al. 2012; Bulgarelli et al. 2013; Vandenkoornhuyse et al. 2015). These insights imply that horizontally acquired genes such as *IbACS* may contribute to the formation of microbially mediated traits that enhance plant fitness under ecologically demanding conditions.

Taken together, these lines of evidence suggest that sweet potato may use *IbACS* for agrocinopine (or its derivative) production as a mechanism to selectively interact with particular soil microorganisms, thereby creating a growth environment that is beneficial under nutrient-limited conditions. In this study, we investigated the role of *IbACS* in sweet potato by evaluating the performance of genome-edited knockout lines and assessing how its disruption affects plant growth and associations with the rhizosphere microbiota.

## RESULTS and DISCUSSION

### Generation of *IbACS*-disrupted sweet potato lines

To investigate the functional contribution of the *IbACS* gene in sweet potato, we generated loss-of-function lines using a CRISPR/Cas9-based approach. A binary vector harboring Cas9 together with four guide RNAs targeting distinct regions of the *IbACS* coding sequence (Fig. 1A) was introduced into *I. batatas* cv. Hanaranman via *Agrobacterium*-mediated transformation, yielding thirteen putative transgenic plants. PCR-based genotyping detected sequence alterations at the corresponding target sites, indicating that genome editing had operated effectively in this hexaploid background. Among the edited plants, three independent lines (KO3, KO8, and KO13) were selected for further analysis because each carried mutations at multiple *IbACS* loci (Fig. 1B), suggesting broad modification of the *IbACS* gene family. Given the hexaploid nature of sweet potato, multiple allelic copies are expected; thus, the observed mutation patterns imply that *IbACS* activity is likely substantially reduced in these lines. When maintained on MS medium under sterile conditions, the knockout lines exhibited no discernible defects in overall growth or morphology relative to empty-vector controls, indicating that disruption of *IbACS* does not markedly affect basal developmental processes during *in vitro* culture.

**Fig. 1.**
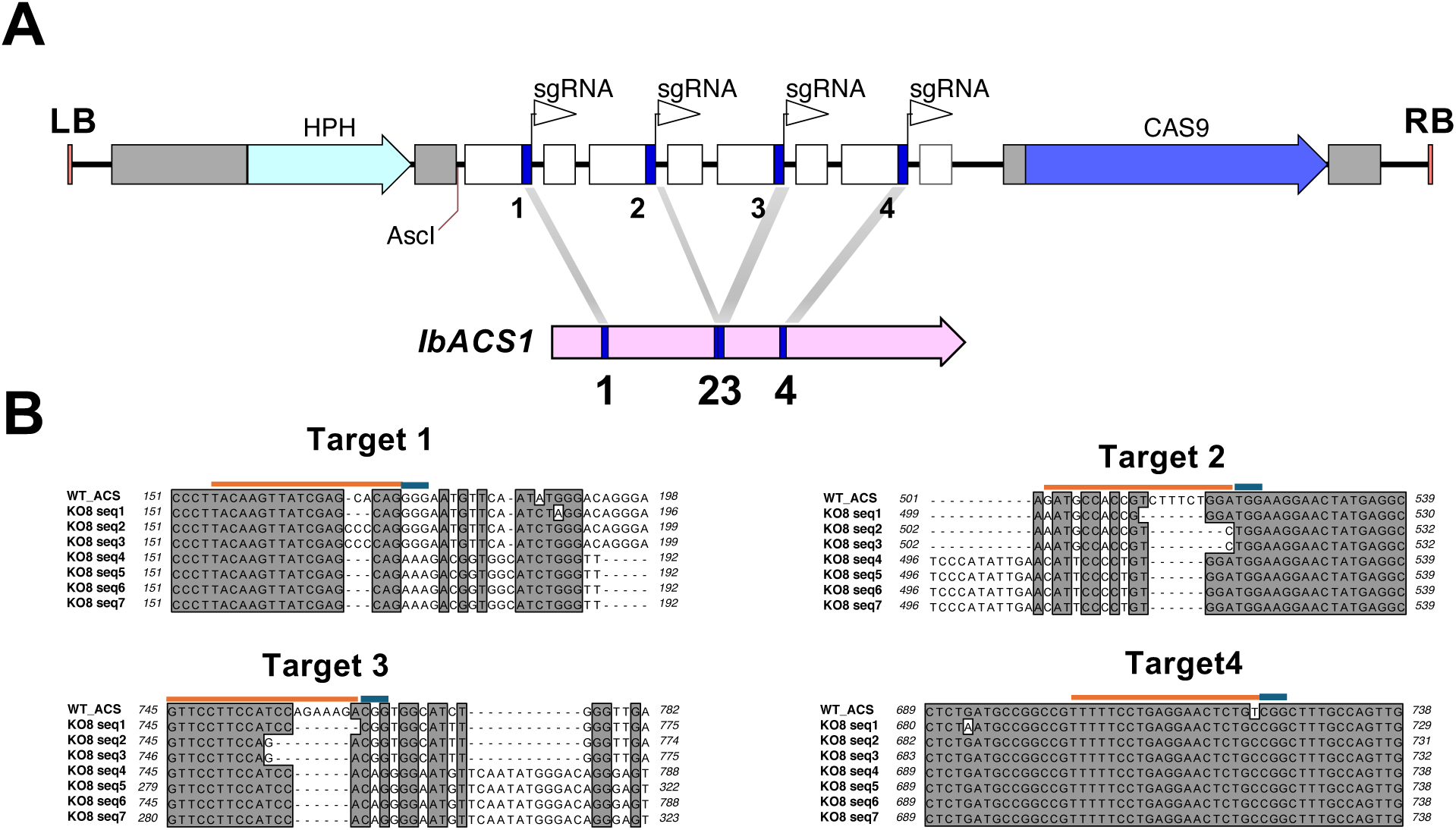
Targeted mutagenesis of *IbACS* in *Ipomoea batatas*. **(A)** Schematic representation of the CRISPR/Cas9 binary vector pZH2BCas9-IbACS used for genome editing of the *IbACS* gene. Four single-guide RNAs (sgRNAs, arrowheads), targeting distinct regions within the *IbACS* coding sequence, were individually inserted into the Arabidopsis U6 promoter-sgRNA scaffold cassette and tandemly cloned into the *Asc*I site of pZH2BCas9. **(B)** Representative sequence alignments showing mutations induced at sgRNA target sites in edited sweet potato line KO8. Insertions and deletions resulting in frameshift mutations are highlighted.

### Growth of *IbACS*-disrupted sweet potato in non-sterile soil

To obtain insight into how *IbACS* disruption influences plant performance, growth of *IbACS*-disrupted lines was evaluated following transfer from MS medium to a non-sterile soil environment. For this assay, a commercial soil supplemented with approximately 10% field soil collected from a sweet potato cultivation plot was used to reproduce conditions that more closely reflect natural rhizosphere environments. Upon transfer into this non-sterile soil, clear growth differences emerged between the *IbACS*-disrupted lines and the vector controls. After two weeks, all three *IbACS*-disrupted lines (KO3, KO8, and KO13) developed smaller shoots and weaker root systems relative to controls, as evidenced by significant reductions in shoot and/or root fresh weight in multiple lines (Fig. 2A and B), indicating that loss of *IbACS* function compromises early vegetative growth under conditions in which plants interact with the native soil microbiota.

**Fig. 2.**
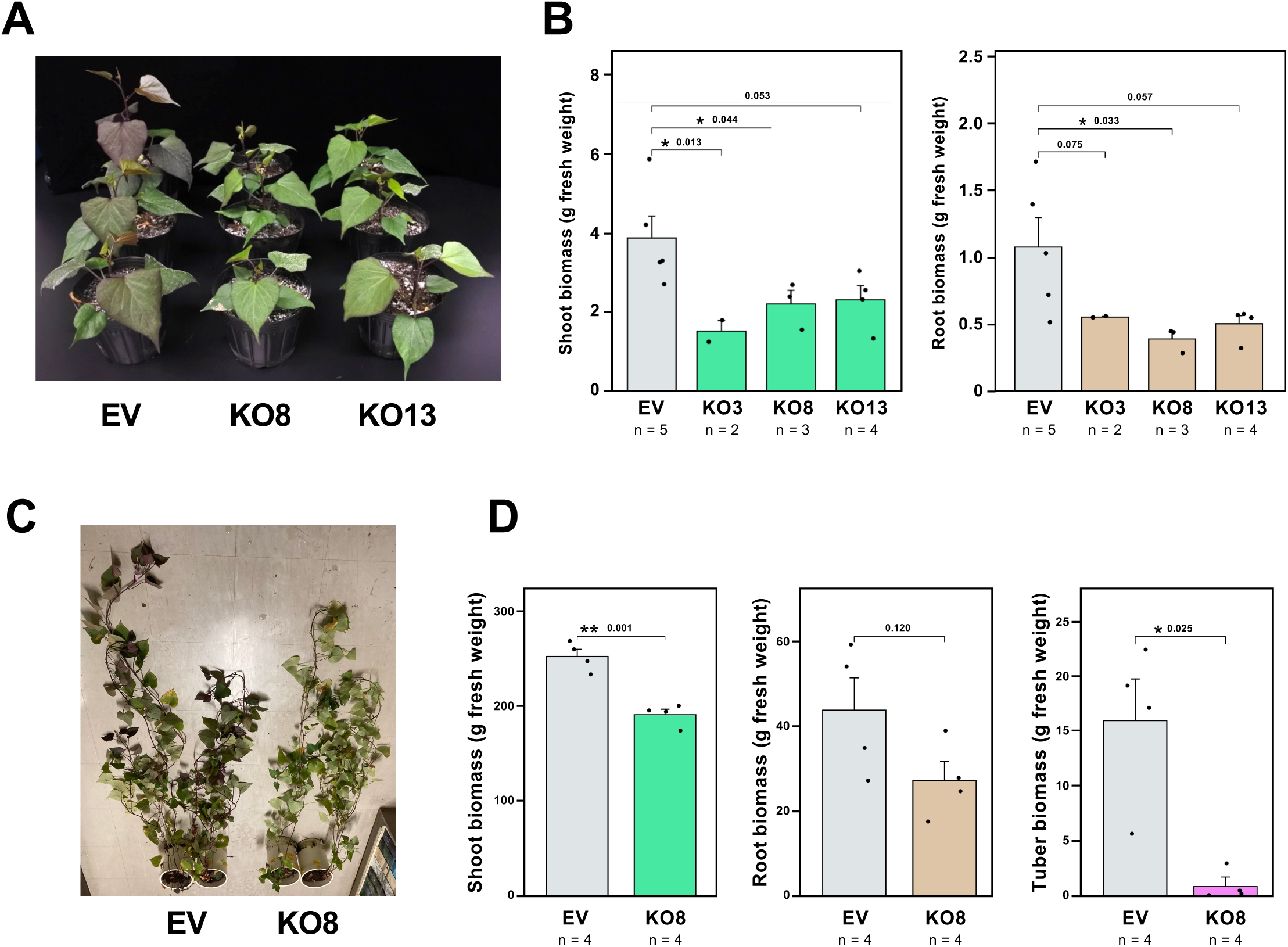
Growth defects of *IbACS*-disrupted sweet potato lines in non-sterile soil. **(A)** Growth phenotypes of control (empty vector, EV) and *IbACS*-disrupted sweet potato lines (KO8 and KO13) grown in non-sterile soil for two weeks after transfer from MS medium. Plants were cultivated in commercial soil supplemented with approximately 10% field soil collected from a sweet potato cultivation plot. **(B)** Quantification of shoot fresh weight (left) and root fresh weight (right) of EV and *IbACS*-disrupted lines KO3, KO8 and KO13 after two weeks of growth in non-sterile soil. Data marked with asterisks are significantly different from control as assessed by two-tailed Student’s *t* test: \**p* < 0.05. **(C)** Growth phenotypes of EV and *IbACS*-disrupted line KO8 grown in non-sterile soil for three months under greenhouse conditions. **(D)** Shoot (left), root (middle) and tuber (right) fresh weight of EV and KO8 plants after three months of cultivation in non-sterile soil. Data marked with asterisks are significantly different from control as assessed by two-tailed Student’s *t* test: \**p* < 0.05. \*\**p* < 0.01.

To further examine this phenotype, control and *IbACS*-disrupted KO8 plants were maintained in a greenhouse for two months, after which fresh weights of shoots and roots were measured separately. Biomass accumulation in both tissues was consistently reduced in KO8. Tuber (storage root) fresh weight was markedly decreased in the *IbACS*-disrupted line KO8 compared with control plants (Fig. 2C and D), indicating that *IbACS* disruption not only impairs early vegetative growth but also compromises the development of belowground sink organs during prolonged cultivation in non-sterile soil. These results collectively support a role for IbACS in sustaining both root system development and carbon allocation to storage tissues under soil-grown conditions.

### Growth of *IbACS*-disrupted sweet potato in sterile soil

To clarify whether the reduced growth of *IbACS*-disrupted plants in non-sterile soil reflects an intrinsic developmental defect or a response dependent on biotic factors in the soil environment, plant growth was next evaluated in sterile soil. Line KO8, which exhibited a representative reduction in growth under non-sterile conditions, was selected for further analysis. When *IbACS*-disrupted KO8 and control plants were grown in sterilized soil (prepared by autoclaving the same soil mixture used in the non-sterile experiments), the overall growth of control plants was markedly reduced relative to their performance in non-sterile soil (Figs. 2 and 3). Under these sterile conditions, KO8 showed no significant differences from the control, indicating that the growth impairment observed in non-sterile soil does not appear in the absence of a soil microbial community (Fig. 3). Consistent with these observations, no pronounced differences in growth performance were detected when plants were maintained on MS medium (data not shown). Taken together, these findings suggest that the impaired growth phenotype of *IbACS*-disrupted plants is not attributable to an intrinsic developmental abnormality, but instead emerges only under conditions that permit interactions with soil microorganisms, supporting a role for IbACS in microbe-dependent growth promotion in sweet potato.

**Fig. 3.**
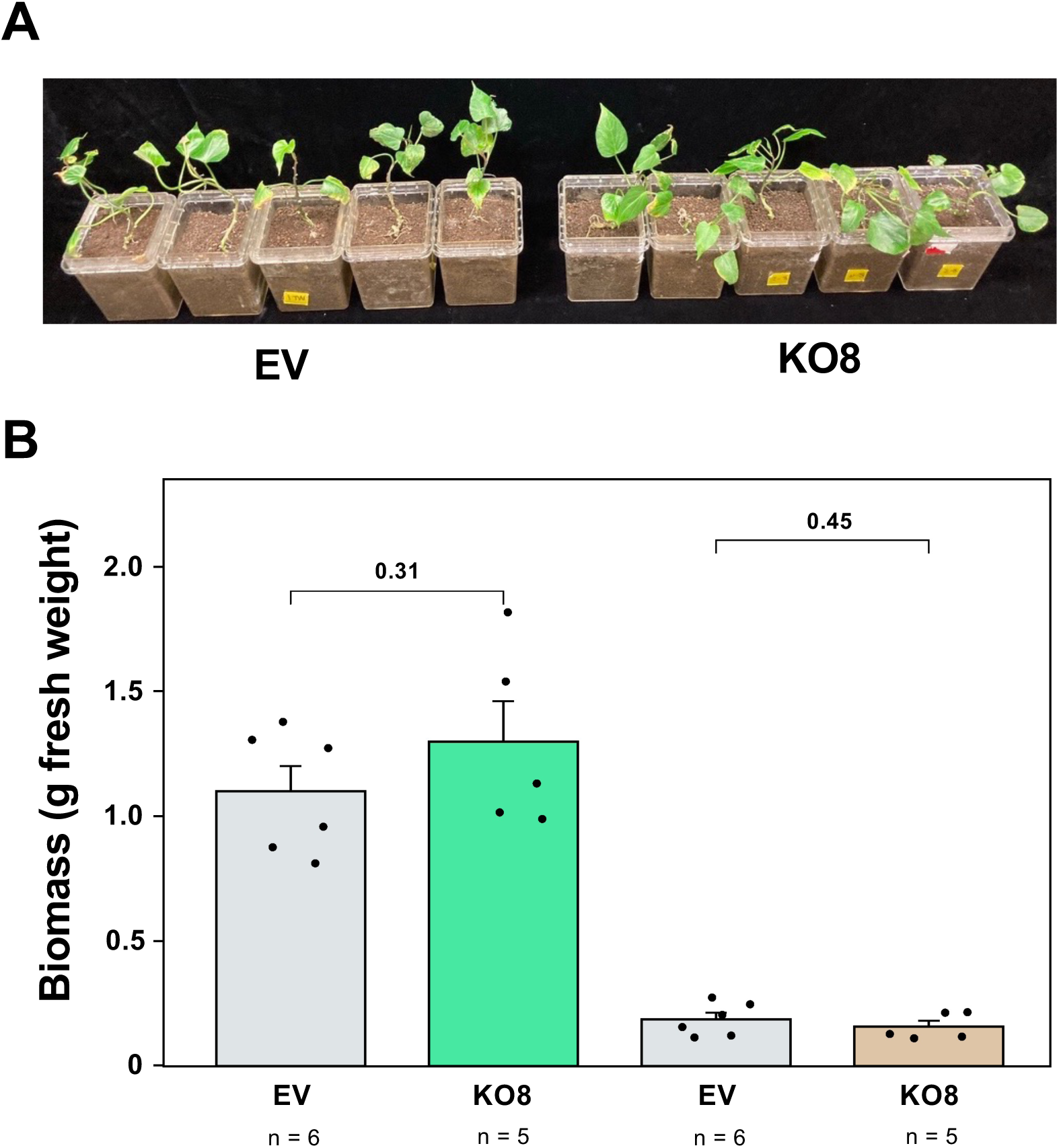
Growth of *IbACS*-disrupted sweet potato lines under sterile soil conditions. **(A)** Growth phenotypes of control (empty vector, EV) and *IbACS*-disrupted sweet potato line KO8 grown in sterilized soil for four weeks after transfer from MS medium. **(B)** Quantification of shoot and root fresh weight of EV and *IbACS*-disrupted KO8 plants after four weeks of growth in sterilized soil. Bars represent mean values ± SE (n = 5), and individual data points indicate biological replicates. Statistical significance was assessed by two-tailed Student’s *t*-test.

### Metagenomic analysis of rhizosphere community shifts (family-level) associated with *IbACS* disruption

Because growth differences between the control plants and the *IbACS* knockout lines became evident consistently when grown in soil containing field-derived sweet potato cultivation soil (Fig. S1), the potential influence of *IbACS* disruption on the rhizosphere microbial community was examined using a metagenomic analysis. Metagenomic sequencing of rhizosphere samples collected from the roots of control EV and *IbACS*-disrupted KO8 plants was performed to assess taxonomic shifts in the microbial community.

In the family-level analysis, a large fraction of reads (EV: approximately 60%, KO: approximately 70%) could not be assigned to any known family (Table S1), indicating that much of the rhizosphere microbiota remains taxonomically unresolved at this rank. PCA of the rhizosphere metagenomes showed a clear separation between EV and *IbACS*-disrupted KO8 plants (Fig. 4A). The three KO8 samples formed a tight cluster, whereas EV samples were positioned distantly along PC1, which accounted for 80.8% of the total variance. PC2 explained 16.5% of the variance and reflected within-group variability. This distinct separation indicates that loss of IbACS function leads to a reproducible, community-wide shift in rhizosphere microbiome composition.

**Fig. 4.**
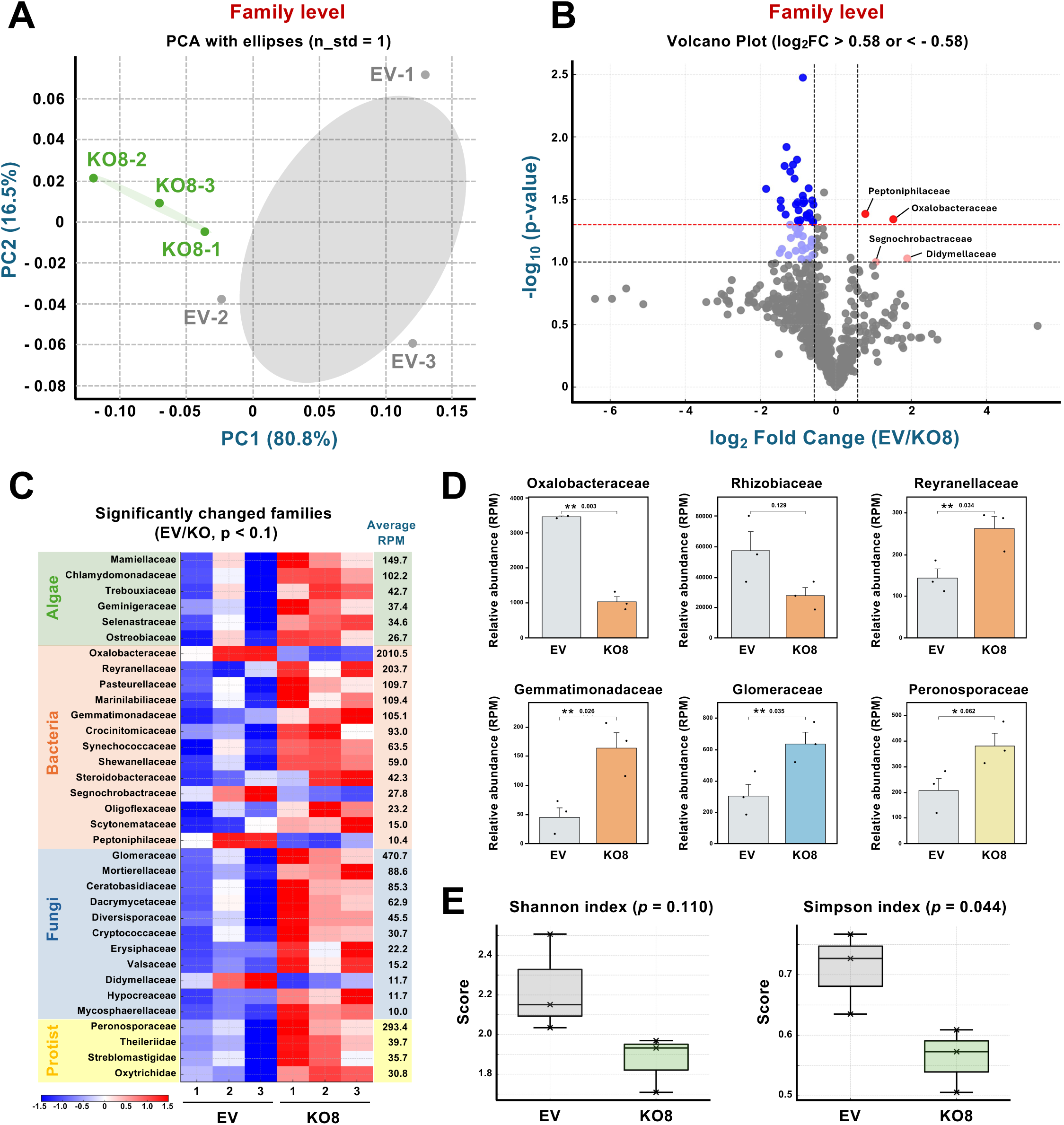
Family-level metagenomic analysis of rhizosphere communities in *IbACS*-disrupted sweet potato. (**A)** Principal component analysis (PCA) was performed using family-level relative abundance profiles obtained from rhizosphere metagenomes of sweet potato roots for the empty vector control (EV) and the *IbACS*-disrupted KO8 line (KO8). Each point represents an individual sample, and the first two principal components (PC1 and PC2) summarize the major axes of variation in microbial community composition. Standard-deviation covariance ellipses (n_std = 1.0) are drawn to indicate within-group variability. (**B)** Volcano plot of family-level differential abundance between EV and KO8 rhizosphere communities. Mean relative abundances of each microbial family were compared, and logLJ fold change (EV/KO) and Welch’s *t*-test *p* values were calculated. Dashed lines indicate the logLJ fold change threshold (±0.58, approx. 1.5-fold), and significance thresholds (*p* < 0.1 and *p* < 0.05). Red and blue dots represent families enriched in EV and KO8, respectively. **(C)** Heatmap showing the relative abundances of differentially abundant microbial families in the rhizosphere of EV and KO8 plants. Values were standardized by row (Z-score transformation) to emphasize relative changes across samples. Families are grouped by major taxonomic categories (Bacteria, Fungi, Protist, and Algae), and the average RPM across all samples is shown on the right. **(D)** Relative abundances expressed as reads per million (RPM) of selected representative families showing *IbACS*-dependent differences between EV and KO8 plants. Statistical significance was evaluated using Welch’s *t*-test: \*\**p* < 0.05, \**p* < 0.1. **(E)** α-diversity metrics of rhizosphere microbial communities at the family level. Shannon (left) and Simpson (right) diversity values were calculated for EV and KO8 samples. Boxes represent interquartile ranges, horizontal lines indicate medians, whiskers show the range. Statistical significance was assessed using Welch’s t-test.

Table 1 summarizes the differentially abundant families identified in this analysis, including their normalized read counts (RPM), fold-change values, and statistical significance. Taxa showing > 1.5-fold differences (logLJFC > 0.58, where logLJFC is defined as logLJ(EV/KO)) and p < 0.1 were considered differentially abundant, and the table lists families that increased or decreased in KO8 relative to EV. Differential abundance analysis revealed several microbial groups whose RPM values were relatively high and showed significant changes between control and *IbACS*-disrupted plants (Fig. 4B, C and D). Among the differentially abundant bacterial families, Oxalobacteraceae (Betaproteobacteria, Burkholderiales) represented the most prominent group, showing a marked reduction in *IbACS*-disrupted KO8 plants and a corresponding enrichment in EV plants (logLJFC = 1.53). This pattern indicates that Oxalobacteraceae is one of the dominant bacterial families positively associated with IbACS activity and likely dependent on IbACS-derived agrocinopine-related compounds.

**Table 1.** Differentially abundant rhizosphere microbial families in the IbACS-KO8 sweet potato line, selected from 781 detected families (logLJFC > 0.58 or < −0.58 and *p* < 0.1.)

| Family | Major_group | Higher_taxon | EV-1<br>(RPM) | EV-2<br>(RPM) | EV-3<br>(RPM) | KO8-1<br>(RPM) | KO8-2<br>(RPM) | KO8-3<br>(RPM) | log2FC<br>(EV/KO8) | p value |
| --- | --- | --- | --- | --- | --- | --- | --- | --- | --- | --- |
| Didymellaceae | Fungi | Ascomycota (Pleosporales) | 9.71 | 19.37 | 26.37 | 1.86 | 5.03 | 8.01 | 1.90 | 0.094 |
| Oxalobacteraceae | Bacteria | Pseudomonadota (Betaproteobacteria) | 2029.06 | 3438.05 | 3486.69 | 1306.97 | 818.45 | 983.76 | 1.53 | 0.046 |
| Segnochlorobacteraceae | Bacteria | Pseudomonadota (Alphaproteobacteria) | 25.69 | 36.87 | 50.07 | 21.56 | 17.36 | 14.98 | 1.06 | 0.100 |
| Peptoniphilaceae | Bacteria | Bacillota (Clostridia/Tissierellia) | 10.36 | 14.61 | 14.35 | 7.29 | 6.79 | 8.78 | 0.78 | 0.041 |
| Valsaceae | Fungi | Ascomycota (Diaporthales) | 11.66 | 12.91 | 11.68 | 19.85 | 15.60 | 19.38 | -0.60 | 0.035 |
| Selenastraceae | Algae | Chlorophyta (Sphaeropleales) | 26.12 | 34.32 | 21.86 | 39.40 | 42.02 | 43.66 | -0.60 | 0.048 |
| Marinilabiliaceae | Bacteria | Bacteroidota (Marinilabiliales) | 78.36 | 107.88 | 74.11 | 153.10 | 111.96 | 130.72 | -0.60 | 0.048 |
| Pseudeurotiaceae | Fungi | Ascomycota (Leotiomycetes) | 8.85 | 5.27 | 5.67 | 8.07 | 10.57 | 11.88 | -0.62 | 0.088 |
| Astrephomenaceae | Algae | Chlorophyta (Chlamydomonadales) | 3.45 | 5.44 | 4.34 | 6.20 | 7.30 | 6.98 | -0.63 | 0.032 |
| Synechococcaceae | Bacteria | Cyanobacteria | 39.50 | 65.24 | 44.23 | 77.87 | 80.26 | 74.14 | -0.64 | 0.066 |
| Pasteurellaceae | Bacteria | Pseudomonadota (Gammaproteobacteria) | 81.16 | 109.75 | 65.76 | 162.09 | 125.30 | 113.93 | -0.64 | 0.069 |
| Trebouxiaceae | Algae | Chlorophyta (Trebouxiphyceae) | 29.14 | 45.53 | 24.03 | 45.76 | 57.36 | 54.25 | -0.67 | 0.075 |
| Scytonemataceae | Bacteria | Cyanobacteria | 9.50 | 10.02 | 15.02 | 17.06 | 16.61 | 21.70 | -0.68 | 0.045 |
| Microascaceae | Fungi | Ascomycota (Microascales) | 1.73 | 1.70 | 1.00 | 2.64 | 2.26 | 2.33 | -0.71 | 0.041 |
| Sclerotiniaceae | Fungi | Ascomycota (Helotiales) | 4.32 | 2.72 | 3.17 | 5.12 | 5.28 | 6.46 | -0.72 | 0.026 |
| Steroidobacteraceae | Bacteria | Pseudomonadota (Gammaproteobacteria) | 25.04 | 29.73 | 39.56 | 37.23 | 59.38 | 62.78 | -0.76 | 0.095 |
| Chlamydomonadaceae | Algae | Chlorophyta (Chlamydomonadales) | 71.66 | 100.40 | 54.41 | 133.24 | 134.61 | 118.84 | -0.77 | 0.044 |
| Geminigeraceae | Algae | Cryptophyta (Cryptophyceae) | 30.22 | 34.49 | 18.19 | 55.06 | 47.05 | 39.53 | -0.77 | 0.042 |
| Serendipitaceae | Fungi | Basidiomycota (Sebacinales) | 4.32 | 2.89 | 2.34 | 4.65 | 6.29 | 5.43 | -0.78 | 0.042 |
| Ostreobiaceae | Algae | Chlorophyta (Bryopsidales) | 14.25 | 29.22 | 15.02 | 37.07 | 36.23 | 28.42 | -0.80 | 0.078 |
| Nephroselmidaceae | Algae | Chlorophyta (Nephroselmidales) | 1.30 | 2.38 | 2.17 | 3.57 | 4.03 | 2.84 | -0.84 | 0.033 |
| Chordaceae | Algae | Ochrophyta (Laminariales) | 4.75 | 5.78 | 3.51 | 9.93 | 6.79 | 8.78 | -0.86 | 0.032 |
| Reyranellaceae | Bacteria | Pseudomonadota (Alphaproteobacteria) | 135.34 | 110.60 | 186.27 | 288.66 | 205.56 | 295.80 | -0.87 | 0.034 |
| Mycosphaerellaceae | Fungi | Ascomycota (Mycosphaerellales) | 6.91 | 7.81 | 6.51 | 14.27 | 12.08 | 12.66 | -0.88 | 0.003 |
| Peronosporaceae | Protist | Oomycota (Peronosporales) | 231.62 | 270.97 | 117.50 | 476.50 | 349.22 | 314.40 | -0.88 | 0.062 |
| Shewanellaceae | Bacteria | Pseudomonadota (Gammaproteobacteria) | 36.91 | 54.87 | 32.88 | 78.02 | 78.25 | 73.11 | -0.88 | 0.030 |
| Mamiellaceae | Algae | Chlorophyta (Mamiellophyceae) | 88.93 | 163.26 | 62.76 | 235.61 | 188.20 | 159.40 | -0.89 | 0.081 |
| Streblomastigidae | Protist | Parabasalia | 28.28 | 33.98 | 12.02 | 55.84 | 49.06 | 34.88 | -0.91 | 0.073 |
| Caulerpacaeae | Algae | Chlorophyta (Bryopsidales) | 1.30 | 1.02 | 2.50 | 2.95 | 2.26 | 3.88 | -0.92 | 0.095 |
| Ascolobaceae | Fungi | Ascomycota (Pezizales) | 0.86 | 1.70 | 1.00 | 2.79 | 2.26 | 1.81 | -0.95 | 0.046 |
| Dacrymycetaceae | Fungi | Basidiomycota (Dacrymycetales) | 37.99 | 64.22 | 26.04 | 101.13 | 77.49 | 70.27 | -0.96 | 0.053 |
| Ceratobasidiaceae | Fungi | Basidiomycota (Cantharellales) | 52.67 | 84.43 | 35.05 | 140.07 | 101.14 | 98.17 | -0.98 | 0.048 |
| Prochlorococcaceae | Bacteria | Cyanobacteria | 4.75 | 4.93 | 6.84 | 7.76 | 12.33 | 12.66 | -0.99 | 0.059 |
| Crocinitomicaceae | Bacteria | Bacteroidota (Flavobacteriales) | 70.37 | 76.45 | 39.89 | 133.09 | 144.92 | 93.00 | -0.99 | 0.039 |
| Cryptococcaceae | Fungi | Basidiomycota (Tremellomycetes) | 16.41 | 30.07 | 14.52 | 51.65 | 36.23 | 35.13 | -1.01 | 0.047 |
| Oxytrichidae | Protist | Ciliophora (Spirotrichea) | 26.33 | 23.44 | 11.35 | 38.16 | 43.78 | 41.85 | -1.02 | 0.033 |
| Erysiphaceae | Fungi | Ascomycota (Erysiphales) | 12.74 | 15.63 | 15.52 | 32.88 | 21.64 | 34.88 | -1.03 | 0.060 |
| Theileriidae | Protist | Apicomplexa (Piroplasmida) | 29.14 | 32.79 | 16.36 | 60.65 | 54.09 | 45.47 | -1.03 | 0.015 |
| Glomeraceae | Fungi | Glomeromycota | 294.86 | 436.79 | 182.76 | 780.37 | 605.60 | 523.91 | -1.06 | 0.035 |
| Candidatus Ozmobacteraceae | Bacteria | Candidatus Ozmobacteria | 0.43 | 1.02 | 1.34 | 2.02 | 2.01 | 1.81 | -1.07 | 0.054 |
| Coniochaetaceae | Fungi | Ascomycota (Coniochaetales) | 6.04 | 3.06 | 1.50 | 7.91 | 5.79 | 8.53 | -1.07 | 0.081 |
| Diversisporaceae | Fungi | Glomeromycota | 29.57 | 40.60 | 16.69 | 72.75 | 56.86 | 56.58 | -1.10 | 0.022 |
| Mortierellaceae | Fungi | Mortierellomycota | 53.96 | 70.67 | 41.23 | 107.18 | 111.21 | 147.51 | -1.14 | 0.017 |
| Thermoproteaceae | Archaea | Thermoproteota (Thermoproteia) | 1.08 | 1.02 | 0.33 | 1.24 | 2.01 | 2.33 | -1.20 | 0.064 |
| Melampsoraceae | Fungi | Basidiomycota (Pucciniales) | 4.96 | 9.00 | 3.84 | 15.05 | 14.84 | 11.37 | -1.21 | 0.019 |
| Geoglossaceae | Fungi | Ascomycota (Geoglossales) | 1.08 | 1.36 | 1.17 | 2.02 | 3.27 | 3.10 | -1.22 | 0.050 |
| Rubritaleaceae | Bacteria | Verrucomicrobiota (Verrucomicrobiae) | 1.30 | 1.36 | 2.34 | 2.79 | 5.54 | 3.62 | -1.26 | 0.088 |
| Gelatosporiaceae | Fungi | Basidiomycota (Polyporales) | 3.67 | 6.80 | 3.17 | 11.48 | 12.33 | 10.08 | -1.31 | 0.012 |
| Hypocreaceae | Fungi | Ascomycota (Hypocreales) | 5.18 | 7.48 | 7.18 | 15.82 | 13.08 | 21.18 | -1.34 | 0.042 |
| Desulfonatronaceae | Bacteria | Desulfobacterota (Desulfovibrionia) | 1.73 | 1.19 | 2.00 | 3.72 | 3.77 | 5.17 | -1.36 | 0.017 |
| Stachybotryaceae | Fungi | Ascomycota (Hypocreales) | 2.81 | 4.42 | 1.50 | 9.77 | 4.53 | 9.30 | -1.44 | 0.079 |
| Oligoflexaceae | Bacteria | Bdellovibrionota (Oligoflexia) | 4.75 | 20.22 | 12.18 | 25.75 | 44.03 | 32.55 | -1.46 | 0.037 |
| Tuberaceae | Fungi | Ascomycota (Pezizales) | 0.65 | 0.68 | 0.67 | 1.71 | 2.26 | 1.55 | -1.47 | 0.032 |
| Thyridiaceae | Fungi | Ascomycota (Pleosporales) | 2.37 | 1.53 | 0.33 | 5.27 | 2.26 | 4.39 | -1.49 | 0.085 |
| Gemmatimonadaceae | Bacteria | Gemmatimonadota (Gemmatimonadetes) | 16.41 | 47.74 | 72.44 | 117.26 | 168.07 | 208.74 | -1.85 | 0.026 |

RPM, reads per million. See Table S1 for full list of all data for 781 families detected.

**Table 2.** Differentially abundant rhizosphere microbial genera in the IbACS-KO8 sweet potato line, selected from 986 detected genera (logLJFC > 0.58 or < −0.58 and *p* < 0.1).

| Genus | Major_group | Higher_taxon | EV-1<br>(RPM) | EV-2<br>(RPM) | EV-3<br>(RPM) | KO8-1<br>(RPM) | KO8-2<br>(RPM) | KO8-3<br>(RPM) | log<sub>2</sub>FC<br>(EV/KO8) | p value |
| --- | --- | --- | --- | --- | --- | --- | --- | --- | --- | --- |
| *Massilia* | Bacteria | Pseudomonadota (Betaproteobacteria) | 1190.74 | 2689.44 | 2173.65 | 570.66 | 358.54 | 388.21 | 2.20 | 0.066 |
| *Ensifer* | Bacteria | Pseudomonadota (Alphaproteobacteria) | 3896.35 | 7310.91 | 9818.89 | 2986.94 | 340.05 | 1674.86 | 2.07 | 0.072 |
| *Pullulanibacillus* | Bacteria | Bacillota (Bacilli) | 12.36 | 25.12 | 15.79 | 9.81 | 6.84 | 3.64 | 1.39 | 0.084 |
| *Segnochromobacterum* | Bacteria | Pseudomonadota (Alphaproteobacteria) | 25.80 | 37.08 | 50.39 | 21.65 | 17.47 | 15.09 | 1.06 | 0.100 |
| *Planctomonas* | Algae | Cryptophyta (Cryptophyceae) | 50.52 | 61.85 | 82.31 | 52.35 | 17.47 | 31.22 | 0.95 | 0.086 |
| *Saccharibacillus* | Bacteria | Bacillota (Bacilli) | 13.23 | 23.41 | 16.97 | 12.77 | 7.34 | 9.37 | 0.86 | 0.095 |
| *Pseudogymnoascus* | Fungi | Ascomycota (Pseudogymnoascales) | 8.89 | 5.30 | 5.71 | 8.10 | 10.63 | 11.97 | -0.62 | 0.088 |
| *Coccomyxa* | Algae | Chlorophyta (Trebouxiophyceae) | 50.95 | 60.31 | 44.85 | 69.95 | 81.03 | 90.03 | -0.63 | 0.020 |
| *Synechococcus* | Bacteria | Cyanobacteria | 39.68 | 65.27 | 44.01 | 77.12 | 79.76 | 74.15 | -0.63 | 0.069 |
| *Enterococcus* | Bacteria | Bacillota (Bacilli) | 125.32 | 203.49 | 105.66 | 278.40 | 218.51 | 194.62 | -0.67 | 0.095 |
| *Pseudonocardia* | Bacteria | Actinomycetota (Actinomycetia) | 2334.65 | 3072.84 | 2518.01 | 5108.01 | 3298.74 | 4259.08 | -0.68 | 0.079 |
| *Trebouxia* | Algae | Chlorophyta (Trebouxiophyceae) | 26.23 | 44.76 | 22.85 | 43.93 | 56.46 | 51.26 | -0.69 | 0.087 |
| *Raphidocelis* | Algae | Chlorophyta (Sphaeroleales) | 12.14 | 16.06 | 10.58 | 20.25 | 20.51 | 22.12 | -0.70 | 0.028 |
| *Muricauda* | Bacteria | Bacteroidota (Flavobacteriales) | 16.91 | 20.33 | 14.78 | 31.78 | 31.14 | 22.12 | -0.71 | 0.053 |
| *Guillardia* | Algae | Cryptophyta (Cryptophyceae) | 29.92 | 34.34 | 17.97 | 55.31 | 46.08 | 39.81 | -0.78 | 0.043 |
| *Chlamydomonas* | Algae | Chlorophyta (Chlamydomonadales) | 69.81 | 95.68 | 52.07 | 129.93 | 131.92 | 115.27 | -0.79 | 0.037 |
| *Ostreobium* | Algae | Chlorophyta (Bryopsidales) | 14.31 | 29.39 | 15.12 | 37.23 | 36.46 | 28.62 | -0.80 | 0.078 |
| *Mammaliococcus* | Bacteria | Bacillota (Bacilli) | 16.69 | 26.14 | 11.93 | 40.35 | 30.89 | 26.80 | -0.84 | 0.068 |
| *Scytonema* | Bacteria | Cyanobacteria | 7.59 | 4.44 | 6.55 | 8.57 | 10.89 | 14.05 | -0.85 | 0.067 |
| *Reyranella* | Bacteria | Pseudomonadota (Alphaproteobacteria) | 135.94 | 111.23 | 187.46 | 289.92 | 206.87 | 297.92 | -0.87 | 0.034 |
| *Micromonas* | Algae | Chlorophyta (Mamiellophyceae) | 89.11 | 163.51 | 62.82 | 235.55 | 188.38 | 159.50 | -0.89 | 0.082 |
| *Shewanella* | Bacteria | Pseudomonadota (Gammaproteobacteria) | 36.21 | 54.33 | 31.92 | 78.05 | 77.99 | 71.81 | -0.90 | 0.028 |
| *Shigella* | Bacteria | Pseudomonadota (Gammaproteobacteria) | 42.28 | 66.46 | 30.24 | 111.54 | 80.27 | 69.47 | -0.91 | 0.072 |
| *Streblomastix* | Protist | Metamonada | 28.40 | 34.17 | 12.09 | 56.08 | 49.37 | 35.13 | -0.91 | 0.073 |
| *Actinomyces* | Bacteria | Actinomycetota (Actinomycetia) | 23.42 | 35.88 | 35.11 | 47.67 | 58.24 | 77.02 | -0.95 | 0.056 |
| *Glomus* | Fungi | Glomeromycota | 17.13 | 33.49 | 12.93 | 50.63 | 40.01 | 33.82 | -0.97 | 0.067 |
| *Calocera* | Fungi | Basidiomycota (Dacrymycetales) | 36.86 | 63.73 | 25.53 | 100.17 | 77.23 | 70.51 | -0.97 | 0.052 |
| *Rhizoctonia* | Fungi | Basidiomycota (Cantharellales) | 49.43 | 81.67 | 34.10 | 138.03 | 98.24 | 93.41 | -1.00 | 0.052 |
| *Stylonychia* | Protist | Ciliophora | 25.80 | 22.55 | 10.92 | 37.08 | 42.79 | 40.59 | -1.02 | 0.034 |
| *Theileria* | Protist | Apicomplexa | 29.27 | 32.98 | 16.46 | 60.91 | 54.44 | 45.79 | -1.03 | 0.015 |
| *Cryptococcus* | Fungi | Basidiomycota (Tremellomycetes) | 15.18 | 29.05 | 13.61 | 50.79 | 35.20 | 32.52 | -1.03 | 0.057 |
| *Saitoella* | Fungi | Ascomycota (Taphrinomycetes) | 14.96 | 17.94 | 9.74 | 31.63 | 28.61 | 27.32 | -1.04 | 0.011 |
| *Rhodopila* | Bacteria | Pseudomonadota (Alphaproteobacteria) | 14.96 | 31.44 | 19.65 | 36.61 | 54.44 | 45.01 | -1.04 | 0.030 |
| *Putridiphycobacter* | Bacteria | Bacteroidota | 36.86 | 65.61 | 26.37 | 112.32 | 86.85 | 69.47 | -1.06 | 0.053 |
| *Golovinomyces* | Fungi | Ascomycota (Erysiphales) | 11.92 | 13.33 | 14.28 | 30.38 | 20.76 | 31.48 | -1.06 | 0.047 |
| *Pseudobrythopirellula* | Bacteria | Planctomycetota | 3.25 | 7.18 | 11.42 | 10.59 | 16.46 | 18.73 | -1.07 | 0.077 |
| *Plasmopara* | Protist | Oomycota | 156.97 | 241.42 | 91.72 | 442.29 | 319.29 | 279.97 | -1.09 | 0.050 |
| *Rhizophagus* | Fungi | Glomeromycota | 264.08 | 403.39 | 169.32 | 731.12 | 566.42 | 489.94 | -1.09 | 0.033 |
| *Diversispora* | Fungi | Glomeromycota | 29.70 | 40.83 | 16.80 | 73.07 | 57.22 | 56.98 | -1.10 | 0.022 |
| *Paraclostridium* | Bacteria | Bacillota (Clostridia) | 53.12 | 97.39 | 40.82 | 164.82 | 133.19 | 116.57 | -1.12 | 0.031 |
| *Pasteurella* | Bacteria | Pseudomonadota (Gammaproteobacteria) | 43.36 | 72.44 | 30.91 | 131.95 | 101.28 | 88.47 | -1.13 | 0.031 |
| *Linnemannia* | Fungi | Mortierellomycota | 25.15 | 43.06 | 19.65 | 78.36 | 68.11 | 48.92 | -1.15 | 0.035 |
| *Melampsora* | Fungi | Basidiomycota (Pucciniales) | 4.99 | 9.06 | 3.86 | 15.11 | 14.94 | 11.45 | -1.21 | 0.019 |
| *Trichoderma* | Fungi | Ascomycota (Hypocreales) | 4.77 | 7.18 | 6.72 | 13.40 | 11.65 | 20.03 | -1.27 | 0.065 |
| *Anatolimnocola* | Bacteria | Bacteroidota (Flavobacteriales) | 64.18 | 83.21 | 187.63 | 166.23 | 301.31 | 344.75 | -1.28 | 0.080 |
| *Botrimarina* | Fungi | Ascomycota (Pleosporales) | 4.99 | 9.06 | 21.33 | 19.47 | 27.09 | 39.29 | -1.28 | 0.092 |
| *Pirellulimonas* | Bacteria | Planctomycetota | 4.12 | 10.93 | 20.16 | 18.54 | 30.89 | 36.43 | -1.29 | 0.075 |
| *Acidisphaera* | Bacteria | Pseudomonadota (Alphaproteobacteria) | 42.93 | 57.41 | 44.51 | 74.78 | 164.84 | 133.48 | -1.36 | 0.098 |
| *Candidatus Protochlamydia* | Bacteria | Chlamydiota | 18.65 | 19.14 | 21.17 | 37.70 | 68.87 | 50.22 | -1.41 | 0.068 |
| *Oligoflexus* | Bacteria | Bdellovibrionota (Oligoflexia) | 4.77 | 20.33 | 12.26 | 25.86 | 44.31 | 32.78 | -1.46 | 0.037 |
| *Phragmitibacter* | Bacteria | Bacteroidota | 1.52 | 4.44 | 7.22 | 8.72 | 12.66 | 15.61 | -1.49 | 0.039 |
| *Gemmatimonas* | Bacteria | Gemmatimonadota | 6.72 | 21.87 | 31.24 | 49.23 | 70.90 | 86.90 | -1.79 | 0.025 |
| *Gemmatirosa* | Bacteria | Gemmatimonadota | 5.64 | 18.79 | 30.40 | 51.41 | 77.23 | 94.71 | -2.03 | 0.027 |
| *Youhaiella* | Bacteria | Pseudomonadota (Alphaproteobacteria) | 6.07 | 6.15 | 10.75 | 20.56 | 54.95 | 58.54 | -2.54 | 0.090 |
| *Aquihabitans* | Bacteria | Actinomycetota (Actinomycetia) | 45.96 | 18.45 | 23.69 | 154.39 | 243.84 | 320.56 | -3.03 | 0.044 |

RPM, reads per million. See Table S2 for full list of all data for 986 genera detected.

Rhizobiaceae also showed a relatively high overall abundance and tended to be more abundant in EV than in KO8, although this difference did not reach statistical significance at the family level (Fig. 4D). This trend is nonetheless noteworthy because Rhizobiaceae includes genera that exhibited significant *IbACS* -dependent enrichment at the genus level, as described in the following section.

In contrast to these EV-enriched bacterial families, several other taxa increased in relative abundance in KO8 plants. Among bacteria, Reyranellaceae and Gemmatimonadaceae were more abundant in KO8 than in EV, indicating that disruption of *IbACS* permits the expansion of bacterial lineages that are otherwise less prominent in the EV-associated rhizosphere community. *IbACS*-dependent shifts were also observed among eukaryotic components of the rhizosphere microbiome. The fungal family Glomeraceae, which comprises arbuscular mycorrhizal fungi involved in phosphorus uptake and root nutrient acquisition, was more abundant in KO8 than in control plants. Similarly, the protist family Peronosporaceae, a family of oomycetes that includes downy mildew–causing genera such as *Peronospora* and *Plasmopara*, showed increased abundance in KO8 plants. Although the absolute abundances of these eukaryotic taxa were lower than those of the dominant bacterial families, their enrichment in KO8 suggests that loss of *IbACS* function is accompanied by coordinated changes across bacterial, fungal, and protist lineages.

Consistent with these taxonomic shifts, α-diversity indices also differed between EV and KO8 plants (Fig. 4E). The Shannon index did not show a statistically significant difference, although KO8 tended to exhibit lower values. The Simpson index revealed a clearer reduction in KO8, indicating decreased community evenness. This suggests that *IbACS* disruption alters not only the abundance of specific microbial lineages but also the overall microbiome structure. The reduction in α-diversity is consistent with a simplified microbial community, in which a limited subset of taxa becomes disproportionately dominant or certain groups fail to establish in the KO8 rhizosphere. The observation that several families increased in abundance in KO8 (Fig. 4B and C) likely reflects this collapse in ecological complexity, permitting expansion of taxa that would otherwise remain minor components in a more balanced community associated with the roots of control plants.

### Metagenomic analysis of rhizosphere community shifts (genus-level) associated with *IbACS* disruption

To complement the family-level analysis, genus-level profiling was performed to obtain higher-resolution insights into microbial groups responding to *IbACS* disruption. Similar to the trends observed at the family rank, a substantial proportion of reads could not be assigned confidently at the genus level, reflecting the taxonomic complexity of the rhizosphere microbiota (Table S2). Approximately 70% of reads in EV samples and 80% in KO8 samples could not be assigned to any known genus, indicating that taxonomic resolution remains limited at genus rank in the sweet potato rhizosphere. Nonetheless, the classified fraction revealed clear compositional differences between EV and KO8 samples.

PCA based on genus-level abundances again showed a distinct separation between EV and KO8 plants, with KO8 replicates forming a compact cluster and EV samples dispersed along PC1 (Fig. 5A). This pattern is consistent with the broader restructuring of the microbiome observed at the family level.

**Fig. 5.**
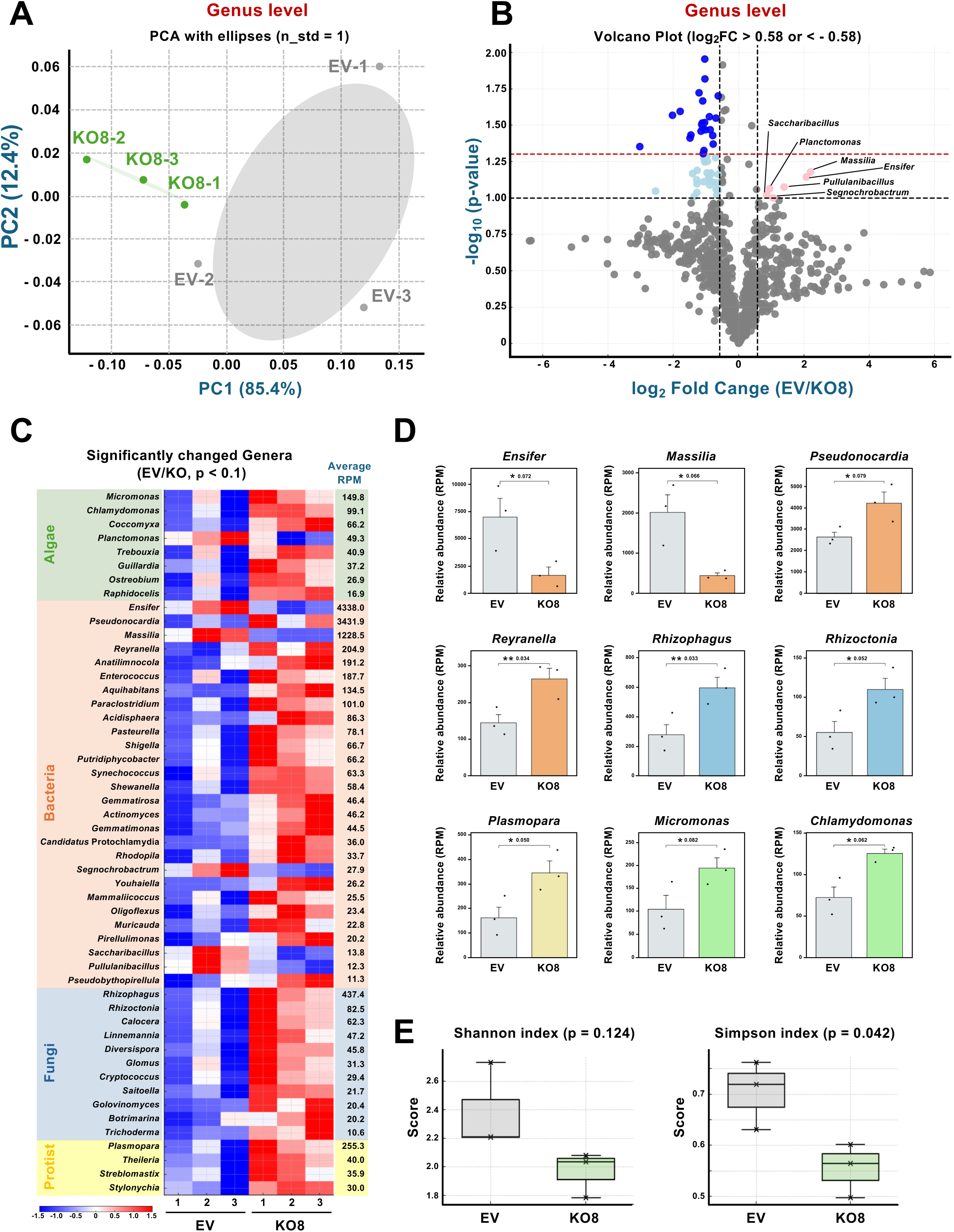
Genus-level metagenomic analysis of rhizosphere communities in *IbACS*-disrupted sweet potato. (**A)** Principal component analysis (PCA) was performed using genus-level relative abundance profiles obtained from rhizosphere metagenomes of sweet potato roots for the empty vector control (EV) and the *IbACS*-disrupted KO8 line (KO8). Each point represents an individual sample, and the first two principal components (PC1 and PC2) summarize the major axes of variation in microbial community composition. Standard-deviation covariance ellipses (n_std = 1.0) are drawn to indicate within-group variability. (**B)** Volcano plot of genus-level differential abundance between EV and KO8 rhizosphere communities. Mean relative abundances of each microbial genus were compared, and logLJ fold change (EV/KO) and Welch’s *t*-test *p* values were calculated. Dashed lines indicate the logLJ fold change threshold (±0.58, approx. 1.5-fold), and significance thresholds (*p* < 0.1 and *p* < 0.05). Red and blue dots represent genera enriched in EV and KO8, respectively. **(C)** Heatmap showing the relative abundances of differentially abundant microbial genera in the rhizosphere of EV and KO8 plants. Values were standardized by row (Z-score transformation) to emphasize relative changes across samples. Genera are grouped by major taxonomic categories (Bacteria, Fungi, Protist, and Algae), and the average RPM across all samples is shown on the right. **(D)** Relative abundances expressed as reads per million (RPM) of selected representative genera showing *IbACS*-dependent differences between EV and KO8 plants. Statistical significance was evaluated using Welch’s *t*-test: \*\**p* < 0.05, \**p* < 0.1. **(E)** α-diversity metrics of rhizosphere microbial communities at the genus level. Shannon (left) and Simpson (right) diversity values were calculated for EV and KO8 samples. Boxes represent interquartile ranges, horizontal lines indicate medians, whiskers show the range. Statistical significance was assessed using Welch’s *t-*test.

Genus-level α-diversity patterns were also consistent with the family-level results. The Shannon index did not differ significantly between EV and KO8 plants, whereas the Simpson index showed reduced evenness in KO8 (Fig. 5E). These trends indicate a simplified community dominated by a smaller number of genera, likely reflecting the same collapse in ecological complexity inferred from family-level analyses.

Table S2 summarizes the differentially abundant genera identified in this analysis, including their normalized read counts (RPM), fold-change values, and statistical significance. Differential abundance analysis identified several genera with relatively high RPM values and significant *IbACS*-dependent changes (Fig. 5B, C and D).

Bacterial genera accounted for the majority of taxa exhibiting significant *IbACS*-dependent changes at the genus level, consistent with the dominant contribution of bacteria to the rhizosphere community. Bacterial genera, *Ensifer* (Rhizobiaceae) and *Massilia* (Oxalobacteraceae) showed the most pronounced enrichment in EV plants (Fig. 5B, C and D). These genus-level patterns closely mirrored the family-level results, in which Oxalobacteraceae and Rhizobiaceae were among the most abundant bacterial families and showed EV-biased trends (Fig. 4).

The genus *Ensifer*, a member of the Alphaproteobacteria within the family Rhizobiaceae, includes well-characterized rhizobial species capable of symbiotic nitrogen fixation, although this trait is not universally conserved across the genus. In addition to nitrogen fixation, *Ensifer* species are known for their metabolic versatility and effective association with plant roots, contributing to nutrient acquisition and rhizosphere competence (Remigi et al. 2016; Poole et al. 2018). *Massilia*, a genus within the Betaproteobacteria (Oxalobacteraceae), is widely distributed in soil and rhizosphere environments and is frequently reported as a root-associated bacterium with strong rhizosphere competence. Members of this genus are characterized by broad metabolic flexibility, including the ability to utilize diverse carbon sources and to respond to plant-derived metabolites (Ofek et al. 2012; Amirhosseini et al. 2025). In contrast, *Pseudonocardia* showed increased abundance in KO8 and displayed relatively high RPM values among KO8-enriched taxa. *Pseudonocardia* is a genus of Actinobacteria commonly found in soil and is known for its ability to utilize complex organic substrates and to produce diverse secondary metabolites, including antimicrobial compounds (Barka et al. 2015). Some bacterial genera were also enriched in KO8 plants, including *Reyranella*, *Anatilimnocola* and others (Fig. 5C and D), despite their relatively lower RPM values. This pattern is consistent with the reduced community evenness observed in KO8 and may reflect the opportunistic expansion of taxa that are less competitive in the rhizosphere of EV plants.

In addition to bacterial taxa, several fungal, protist, and algal genera showed increased abundance in KO8 plants at the genus level. Among fungi, *Rhizophagus* (Glomeraceae), a representative arbuscular mycorrhizal (AM) fungal genus, was enriched in KO8 plants (Fig. 5D). Members of this genus form symbiotic associations with plant roots and are widely recognized for their role in phosphorus acquisition and nutrient exchange (Genre et al. 2020). In contrast, *Rhizoctonia* (Ceratobasidiaceae), a genus that includes soilborne fungi with saprotrophic or pathogenic lifestyles (Nizamani et al. 2025), was also more abundant in KO8 plants (Fig. 5D).

Protist *Plasmopara* (Peronosporaceae), an oomycete genus that includes obligate plant-pathogenic species (Gouveia et al. 2024), also showed increased abundance in KO8 roots. Similarly, several algal genera increased in KO8 plants. *Micromonas* (Mamiellaceae) and *Chlamydomonas* (Chlamydomonadaceae), both unicellular green algae often detected in aquatic environments and soil habitats (Guo et al., 2024; Joseph and Ray 2024), showed higher relative abundances in KO8. These phototrophic or mixotrophic microorganisms are often regarded as opportunistic taxa in soil ecosystems. Together, the increase of minor taxa suggests that disruption of IbACS may alter rhizosphere conditions in a manner that permits the expansion of organisms typically suppressed in a more balanced microbial community.

## Conclusion remarks

Our findings demonstrate that the horizontally acquired *IbACS* gene plays a key role in enabling sweet potato to establish and maintain a functionally robust rhizosphere microbiome that supports plant growth in soil environments. Disruption of *IbACS* resulted in a pronounced reduction in plant biomass specifically under non-sterile conditions, indicating that the observed growth defects arise not from intrinsic developmental abnormalities but from altered interactions with soil microorganisms.

Metagenomic profiling revealed that *IbACS* disruption triggers coordinated shifts in rhizosphere community composition across multiple taxonomic levels. These changes included a decline in several Proteobacteria-related bacterial groups enriched in control plants, such as Oxalobacteraceae-associated genera (e.g., *Massilia*) and Rhizobiaceae-related taxa (e.g., *Ensifer*), alongside reduced community evenness and the expansion of otherwise minor bacterial, fungal, protist, and algal taxa in *IbACS*-disrupted plants. Collectively, these patterns suggest that *IbACS* activity contributes to maintaining a balanced microbial community structure, rather than selectively promoting a single functional guild.

The close correspondence between impaired plant growth and rhizosphere community restructuring supports the idea that *IbACS*-derived metabolites—potentially including agrocinopine-related compounds—act as determinants of microbial recruitment and persistence in the sweet potato rhizosphere. Loss of these signals is likely to diminish the establishment of microbial partners that contribute to nutrient acquisition or stress mitigation, thereby reinforcing a negative feedback loop between reduced root development and simplified microbial community structure.

In addition, the metagenomic data presented here warrant further in-depth interrogation beyond taxonomic profiling. Genome-resolved analyses focusing on enriched metagenome-assembled genomes (MAGs) will be essential to clarify the functional capacities of microbial taxa responding to *IbACS* disruption. In particular, comparative analyses targeting genes related to nitrogen fixation, plant hormone biosynthesis or modulation, and candidate pathways involved in agrocinopine uptake and catabolism may provide critical insights into how *IbACS*-derived metabolites shape rhizosphere function. Such functional genomic approaches will help to move beyond correlative community shifts and more directly link IbACS activity to specific microbial processes that contribute to the growth of sweet potato and rhizosphere stability.

Finally, it should be noted that the rhizosphere community shifts described in this study were characterized using soil supplemented with field soil from a single sweet potato cultivation site. To assess the generality and reproducibility of *IbACS*-dependent microbiome modulation, future studies will need to examine plant performance and microbial community dynamics across soils of distinct origins, physicochemical properties, and microbial backgrounds. Such analyses will be essential to determine whether IbACS-mediated rhizosphere interactions represent a broadly conserved mechanism in sweet potato.

## MATERIALS AND METHODS

### Plant materials and growth conditions

*Ipomoea batatas* cv. Hanaranman was provided by the Kyushu Okinawa Agricultural Research Center (Japan). *In vitro*-grown plants were maintained on Murashige and Skoog (MS) agar medium in a growth chamber at 23 °C under a 16/8 h light/dark photoperiod. Growth experiments under non-sterile conditions were conducted using a commercial soil (Sakata Soil-Mix A, Sakata Seed Corporation, Japan) supplemented with approximately 10% (v/v) field soil collected from a sweet potato cultivation plot at the Togo Farm of Nagoya University (Togo, Japan).

### Vector construction and plant transformation

For CRISPR–Cas9–mediated gene inactivation, pZH2BCas9 was constructed by cloning the 35S–Cas9–rbcsT cassette from pHSE401 (Addgene; Xing et al. 2014) into pZH2B (Kuroda et al. 2010), a binary vector derived from pPZP202 (Hajdukiewicz et al. 1994). Target sequences for genome editing were designed, and the risk of off-target effects was evaluated using appropriate software.

The 20-bp target sequence of *IbACS* was inserted as an annealed oligonucleotide into the *Bsa*I-digested pUC198AMSca vector between the Arabidopsis U6 promoter and the sgRNA scaffold using the In-Fusion HD Cloning Kit (Takara Bio, Japan). Annealed oligonucleotides were prepared by mixing the target oligonucleotides (100 μM each; Table S3), boiling for 1 min, and gradually cooling to 30°C. The resulting AtU6–target sequence–sgRNA scaffold–terminator cassette was introduced into pZH2BCas9 to generate pZH2BCas9-IbACS, which carries four independent sgRNA cassettes. *I. batatas* cv. Hanaranman was transformed as previously described (Otani et al. 1998) using *Agrobacterium tumefaciens* strain EHA101 harboring pZH2BCas9-IbACS.

### DNA isolation and sequencing for metagenomic analysis

Plantlets of *I. batatas* cv. Hanaranman with four to five leaves grown on MS medium were transplanted to sterile or non-sterile soil and cultivated for 7 weeks. Microbial cells were isolated from *I. batatas* roots using a modified protocol based on Ikeda et al. (2009, 2014). Approximately 2-5 g of roots were homogenized in bacterial cell extraction (BCE) buffer (10 ml per g) using a homogenizer (HG30, Hitachi, Japan) for five cycles of 60 s, with cooling on ice for 1 min between each cycle. The homogenates were filtered through Miracloth (Calbiochem, USA) and centrifuged at 5,500 × g for 10 min at 4°C. The resulting pellet was resuspended in 1 mL of BCE buffer, layered onto 1 mL of 80% (w/v) Nycodenz solution prepared in 50 mM Tris–HCl (pH 7.5), and centrifuged at 11,000 × g for 40 min at 4°C. Microbial cells forming a white layer at the interface between the BCE buffer and Nycodenz solution were collected and diluted with an equal volume of sterile water. Cells were pelleted by centrifugation at 11,000 × g for 10 min at 4°C and stored at −80°C until further use. Total DNA was extracted from the microbial cell pellets using the DNeasy PowerSoil Pro Kit (Qiagen, Germany) according to the manufacturer’s instructions. Metagenomic libraries were prepared using the NEBNext Ultra II FS Library Prep Kit for Illumina and NEBNext Multiplex Oligos for Illumina (Unique Dual Index Primer Pairs) (New England Biolabs, USA). Indexed libraries were sequenced in paired-end mode on an Illumina NextSeq 1000 platform (Illumina) at Tokyo University of Agriculture (Japan).

### Metagenomic and bioinformatic analyses

Raw sequencing reads were processed and analyzed using the DOE Systems Biology Knowledgebase (KBase), an integrated platform for reproducible data analysis and sharing in systems biology research (Arkin et al. 2018). Quality-filtered reads were taxonomically classified using Kaiju (v1.9.0), a protein-level metagenomic classifier that assigns sequencing reads based on translated amino acid sequences, enabling sensitive detection of phylogenetically diverse and distantly related microorganisms (Menzel et al. 2016). Taxonomic profiles generated by Kaiju were subsequently normalized to reads per million (RPM) and used for downstream comparative analyses of microbial community composition among samples (Tables S1 and S2).

PCA was conducted on family- or genus-level relative abundance profiles using scikit-learn. Relative abundances were calculated by column-wise normalization, and PCA was performed on the transposed abundance matrix. Group variation was visualized using 1-SD covariance ellipses computed from the sample covariance matrices of EV and KO8 groups. Statistical significance was assessed using a two-tailed Welch’s *t*-test. The resulting *p*-values and logLJ fold changes were visualized in a volcano plot, where taxa with logLJFC ≥ 0.58 (≈1.5-fold) and p < 0.1 were highlighted as differentially abundant, with stricter significance (p < 0.05) indicated by stronger color intensity. Alpha-diversity was calculated at the family level based on the read-count table. For each sample, Shannon’s diversity index and Simpson’s diversity index (Shannon 1948; Simpson 1949) were computed using standard abundance-based formulas after excluding zero-count taxa. Diversity values for EV and KO groups were compared using Welch’s two-sample t-test, and the results were visualized as box-and-whisker plots with individual sample points overlaid.

## ACKNOWLEDGEMENTS

We thank Kyushu Okinawa Agricultural Research Center (NARO, Japan) for providing sweet potato cultivars Hanaranman and Y. Tahara (Nagoya University, Japan) for providing agricultural field soils. This work was supported partially by a Grant-in-Aid for Scientific Research (C) (22K06078), and Grant-in-Aid for Challenging Research (Exploratory, 18K19209) from the Japan Society for the Promotion of Science (JSPS), and Cooperative Research Grant of the Genome Research for BioResource, NODAI Genome Research Center, Tokyo University of Agriculture. The authors used ChatGPT (OpenAI, GPT-5, 2025) only for English editing, and take full responsibility for the final text.

## REFERENCES

Amirhosseini K, Alizadeh M, Azarbad H. (2025) Harnessing the ecological and genomic adaptability of the bacterial genus *Massilia* for environmental and industrial applications. Microb. Biotechnol. 18: e70156. 10.1111/1751-7915.70156

Arkin AP, Cottingham RW, Henry CS, Harris NL, Stevens RL, Maslov S, Dehal P, Ware D, Perez F, Canon S, Sneddon MW, Henderson ML, Riehl WJ, Murphy-Olson D, Chan SY, Kamimura RT, Kumari S, Drake MM, Brettin TS, Glass EM, Chivian D, Gunter D, Weston DJ, Allen BH, Baumohl J, Best AA, Bowen B, Brenner SE, Bun CC, Chandonia JM, Chia JM, Colasanti R, Conrad N, Davis JJ, Davison BH, DeJongh M, Devoid S, Dietrich E, Dubchak I, Edirisinghe JN, Fang G, Faria JP, Frybarger PM, Gerlach W, Gerstein M, Greiner A, Gurtowski J, Haun HL, He F, Jain R, Joachimiak MP, Keegan KP, Kondo S, Kumar V, Land ML, Meyer F, Mills M, Novichkov PS, Oh T, Olsen GJ, Olson R, Parrello B, Pasternak S, Pearson E, Poon SS, Price GA, Ramakrishnan S, Ranjan P, Ronald PC, Schatz MC, Seaver SMD, Shukla M, Sutormin RA, Syed MH, Thomason J, Tintle NL, Wang D, Xia F, Yoo H, Yoo S, Yu D. (2018) KBase: The United States department of energy systems biology knowledgebase. Nat. Biotechnol. 36: 566–569. 10.1038/nbt.4163

Barka EA, Vatsa P, Sanchez L, Gaveau-Vaillant N, Jacquard C, Meier-Kolthoff JP, Klenk HP, Clément C, Ouhdouch Y, van Wezel GP. (2015) Taxonomy, physiology, and natural products of actinobacteria. Microbiol. Mol. Biol. Rev. 80: 1–43. 10.1128/MMBR.00019-15

Berendsen RL, Pieterse CM, Bakker PA. (2012) The rhizosphere microbiome and plant health. Trends Plant Sci. 17: 478–86. 10.1016/j.tplants.2012.04.001

Bulgarelli D, Schlaeppi K, Spaepen S, Ver Loren van Themaat E, Schulze-Lefert P. (2013) Structure and functions of the bacterial microbiota of plants. Annu. Rev. Plant Biol. 64: 807–838. 10.1146/annurev-arplant-050312-120106

Chen K, Dorlhac de Borne F, Szegedi E, Otten L. (2014) Deep sequencing of the ancestral tobacco species *Nicotiana tomentosiformis* reveals multiple T-DNA inserts and a complex evolutionary history of natural transformation in the genus *Nicotiana*. Plant J. 80: 669–682. 10.1111/tpj.12661

Dessaux Y, Faure D. (2018) Niche construction and exploitation by *Agrobacterium*: How to survive and face competition in soil and plant habitats. Curr. Top. Microbiol. Immunol. 418: 55–86. 10.1007/82_2018_83

Genre A, Lanfranco L, Perotto S, Bonfante P. (2020) Unique and common traits in mycorrhizal symbioses. Nat. Rev. Microbiol. 18: 649–660. 10.1038/s41579-020-0402-3

González-Mula A, Lang J, Grandclément C, Naquin D, Ahmar M, Soulère L, Queneau Y, Dessaux Y, Faure D. (2018) Lifestyle of the biotroph *Agrobacterium tumefaciens* in the ecological niche constructed on its host plant. New Phytol. 219: 350–362. 10.1111/nph.15164

Gouveia C, Santos RB, Paiva-Silva C, Buchholz G, Malhó R, Figueiredo A. (2024) The pathogenicity of *Plasmopara viticola*: a review of evolutionary dynamics, infection strategies and effector molecules. BMC Plant Biol. 24: 327. 10.1186/s12870-024-05037-0

Guo X, Pang M, Zheng X, Huang L. (2024) Micromonas, a small pigmented flagellate, predominates the nanoflagellate and photosynthetic picoeukaryote communities in the northern South China Sea. Environ. Microbiol. Rep. 16: e13244. 10.1111/1758-2229.13244

Haimlich S, Fridman Y, Khandal H, Savaldi-Goldstein S, Levy A. (2024) Widespread horizontal gene transfer between plants and bacteria. ISME Commun. 4: ycae073. 10.1093/ismeco/ycae073

Hajdukiewicz P, Svab Z, Maliga P. (1994) The small, versatile pPZP family of *Agrobacterium* binary vectors for plant transformation. Plant Mol Biol. 25: 989–994. 10.1007/BF00014672

Ikeda S, Kaneko T, Okubo T, Rallos LE, Eda S, Mitsui H, Sato S, Nakamura Y, Tabata S, Minamisawa K. (2009) Development of a bacterial cell enrichment method and its application to the community analysis in soybean stems. Microb. Ecol. 58: 703–714. 10.1007/s00248-009-9566-0

Ikeda S, Sasaki K, Okubo T, Yamashita A, Terasawa K, Bao Z, Liu D, Watanabe T, Murase J, Asakawa S, Eda S, Mitsui H, Sato T, Minamisawa K. (2014) Low nitrogen fertilization adapts rice root microbiome to low nutrient environment by changing biogeochemical functions. Microbes Environ. 29: 50–59. 10.1264/jsme2.me13110

Joseph J, Ray JG. (2024) A critical review of soil algae as a crucial soil biological component of high ecological and economic significance. J. Phycol. 60: 229–253. 10.1111/jpy.13444

Kim H, Farrand SK. (1997) Characterization of the acc operon from the nopaline-type Ti plasmid pTiC58, which encodes utilization of agrocinopines A and B and susceptibility to agrocin 84. J. Bacteriol. 179: 7559–7572. 10.1128/jb.179.23.7559-7572.1997

Kuroda M, Kimizu M, Mikami C. (2010) A simple set of plasmids for the production of transgenic plants. Biosci. Biotechnol. Biochem. 74: 2348–2351. 10.1271/bbb.100465

Kyndt T, Quispe D, Zhai H, Jarret R, Ghislain M, Liu Q, Gheysen G, Kreuze JF. (2015) The genome of cultivated sweet potato contains *Agrobacterium* T-DNAs with expressed genes: An example of a naturally transgenic food crop. Proc. Natl. Acad. Sci. USA 112: 5844–5849. 10.1073/pnas.1419685112

Matveeva TV, Otten L. (2019) Widespread occurrence of natural genetic transformation of plants by *Agrobacterium*. Plant Mol. Biol. 101: 415–437. 10.1007/s11103-019-00913-y

Menzel P, Ng KL, Krogh A. (2016) Fast and sensitive taxonomic classification for metagenomics with Kaiju. Nat. Commun. 7: 11257. 10.1038/ncomms11257

Nizamani MM, Zhang Q, Asif M, Khaskheli MA, Wang Y, Li C (2025) Decoding *Rhizoctonia* spp. in-depth genomic analysis, pathogenic mechanisms, and host interactions. Phytopathol. Res. 7: 12. 10.1186/s42483-024-00297-y

Ofek M, Hadar Y, Minz D. (2012) Ecology of root colonizing *Massilia* (Oxalobacteraceae). PLoS One 7: e40117. 10.1371/journal.pone.0040117

Otani M, Shimada T, Kimura T, Saito A. (1998) Transgenic plant production from embryogenic callus of sweet potato (*Ipomoea batatas* (L.) Lam.) using *Agrobacterium tumefaciens*. Plant Biotechnol. 15: 11–16. 10.5511/plantbiotechnology.15.11

Poole P, Ramachandran V, Terpolilli J. (2018) Rhizobia: from saprophytes to endosymbionts. Nat. Rev. Microbiol. 16: 291–303. 10.1038/nrmicro.2017.171

Remigi P, Zhu J, Young JPW, Masson-Boivin C. (2016) Symbiosis within symbiosis: evolving nitrogen-fixing legume symbionts. Trends Microbiol. 24: 63–75. 10.1016/j.tim.2015.10.007

Shannon CE (1948) A mathematical theory of communication. Bell Syst. Tech. J. 27: 379–423. 10.1002/j.1538-7305.1948.tb01338.x

Simpson EH (1949) Measurement of diversity. Nature 163: 688. 10.1038/163688a0

Suzuki K, Yamashita I, Tanaka N. (2002) Tobacco plants were transformed by *Agrobacterium rhizogenes* infection during their evolution. Plant J. 32: 775–787. 10.1046/j.1365-313x.2002.01468.x

Tanaka A, Ryder MH, Suzuki T, Uesaka K, Yamaguchi N, Amimoto T, Otani M, Nakayachi O, Arakawa K, Tanaka N, Takemoto D. (2022) Production of agrocinopine A by *Ipomoea batatas* agrocinopine synthase in transgenic tobacco and its effect on the rhizosphere microbial community. Mol. Plant-Microbe Interact. 35: 73–84. 10.1094/MPMI-05-21-0114-R

Vandenkoornhuyse P, Quaiser A, Duhamel M, Le Van A, Dufresne A. (2015) The importance of the microbiome of the plant holobiont. New Phytol. 206: 1196–1206. 10.1111/nph.13312

Xing HL, Dong L, Wang ZP, Zhang HY, Han CY, Liu B, Wang XC, Chen QJ. (2014) A CRISPR/Cas9 toolkit for multiplex genome editing in plants. BMC Plant Biol. 14: 327. 10.1186/s12870-014-0327-y

Yan M, Li M, Wang Y, Wang X, Moeinzadeh MH, Quispe-Huamanquispe DG, Fan W, Fang Y, Wang Y, Nie H, Wang Z, Tanaka A, Heider B, Kreuze JF, Gheysen G, Wang H, Vingron M, Bock R, Yang J. (2024) Haplotype-based phylogenetic analysis and population genomics uncover the origin and domestication of sweetpotato. Mol. Plant 17: 277–296. 10.1016/j.molp.2023.12.019

