## Supplementary material for "Horizontally acquired *IbACS* gene modulates rhizosphere microbiota and contributes to sweet potato growth": Figure S1

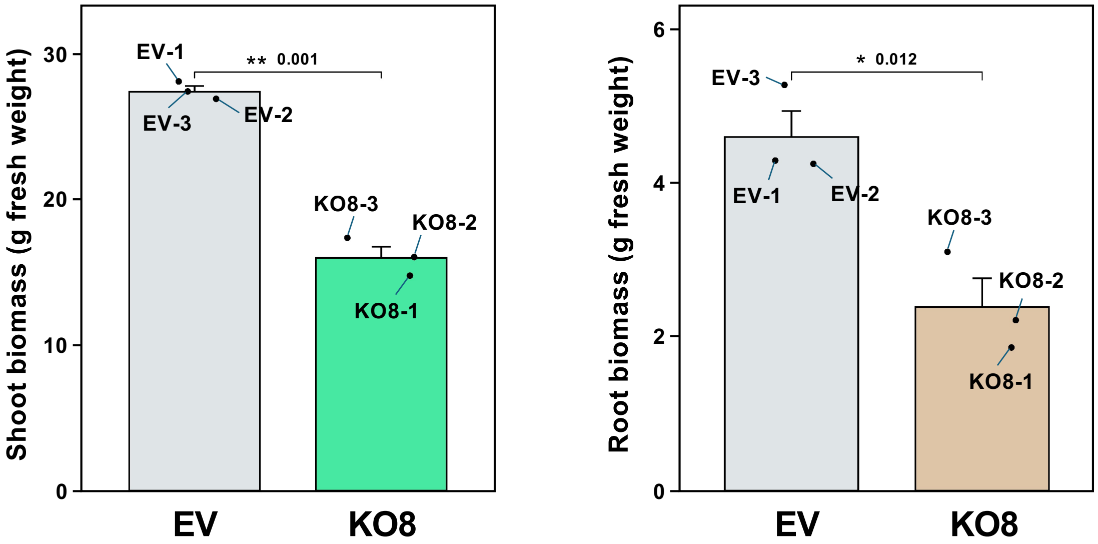


**Fig. S1** Growth phenotypes of control (empty vector, EV) and *IbACS*-disrupted sweet potato line KO8 grown in non-sterile soil for seven weeks after transfer from MS medium. Plants were cultivated in commercial soil supplemented with approximately 10% field soil collected from a sweet potato cultivation plot. The same plant materials were used for subsequent metagenomic analyses. Quantification of shoot fresh weight (left) and root fresh weight (right) of EV and *IbACS*-disrupted line KO8 after 7 weeks of growth in non-sterile soil. Each dot represents an individual biological sample, which corresponds to a rhizosphere sample used for metagenomic analysis. Data marked with asterisks are significantly different from control as assessed by two-tailed Student's *t* test: *$p < 0.05$. **$p < 0.01$.
