## Supplementary material for "Horizontally acquired *IbACS* gene modulates rhizosphere microbiota and contributes to sweet potato growth": Table S1

**Table S1.** Complete list of rhizosphere microbial families detected in the IbACS-KO8 and control sweet potato lines, with normalized read counts, fold-change values, and statistical significance.

| Family | EV-1 (RPM) | EV-2 (RPM) | EV-3 (RPM) | KO8-1 (RPM) | KO8-2 (RPM) | KO8-3 (RPM) | log2FC (EV/KO8) | *p* value |
|---|---|---|---|---|---|---|---|---|
| unclassified | 487243.46 | 594660.31 | 461808.77 | 617637.62 | 698441.62 | 647163.10 | -0.35 | 0.054 |
| cannot be assigned to a (non-viral) family | 129259.68 | 47716.27 | 76491.44 | 62570.81 | 57709.90 | 65121.10 | 0.45 | 0.442 |
| Streptomycetaceae | 28591.86 | 49650.80 | 27376.97 | 32984.96 | 36111.17 | 35196.53 | 0.02 | 0.957 |
| Rhizobiaceae | 37194.65 | 55287.90 | 80752.42 | 37284.19 | 18834.16 | 27951.63 | 1.04 | 0.129 |
| Phyllobacteriaceae | 9802.96 | 4518.54 | 8911.18 | 5657.08 | 12902.72 | 20267.82 | -0.74 | 0.346 |
| Pseudonocardiaceae | 14241.64 | 15341.68 | 12910.61 | 16995.74 | 12117.74 | 17392.24 | -0.13 | 0.525 |
| Bradyrhizobiaceae | 5585.32 | 10022.78 | 15900.42 | 8483.21 | 13750.61 | 16280.86 | -0.29 | 0.571 |
| Sphingomonadaceae | 7200.80 | 20051.34 | 30223.91 | 16037.62 | 16643.24 | 13617.11 | 0.31 | 0.633 |
| Brucellaceae | 6949.11 | 451.23 | 1325.24 | 13546.52 | 5475.79 | 8326.05 | -1.65 | 0.119 |
| Mycobacteriaceae | 2057.56 | 2883.36 | 15681.11 | 2099.44 | 5341.94 | 8030.76 | 0.41 | 0.744 |
| Xanthomonadaceae | 120749.49 | 1285.38 | 24203.73 | 16832.72 | 6009.93 | 7552.06 | 2.27 | 0.402 |
| Microbacteriaceae | 20468.70 | 18078.42 | 52607.51 | 31043.74 | 4886.80 | 7501.17 | 1.07 | 0.321 |
| Paenibacillaceae | 9621.64 | 32085.45 | 12844.35 | 15671.09 | 4541.10 | 7244.89 | 0.99 | 0.333 |
| Nocardioidaceae | 8669.28 | 2640.42 | 8905.84 | 2862.90 | 4318.94 | 6976.99 | 0.51 | 0.454 |
| Gemmataceae | 1566.48 | 793.38 | 2385.27 | 889.10 | 6488.98 | 6970.02 | -1.60 | 0.239 |
| Burkholderiaceae | 5668.42 | 14970.13 | 10275.81 | 11172.85 | 7526.82 | 4992.17 | 0.38 | 0.503 |
| Nocardiaceae | 1323.86 | 7115.81 | 8195.15 | 1133.09 | 3497.47 | 4519.14 | 0.86 | 0.371 |
| Boseaceae | 2053.45 | 4937.49 | 10254.45 | 6128.15 | 2112.42 | 4456.11 | 0.44 | 0.611 |
| Devosiaceae | 3373.42 | 8415.63 | 9534.75 | 2160.71 | 4057.03 | 4176.84 | 1.04 | 0.186 |
| Hyphomicrobiaceae | 1004.60 | 512.05 | 1123.29 | 556.08 | 2942.19 | 3584.47 | -1.42 | 0.247 |
| Xanthobacteraceae | 7362.69 | 4289.70 | 9539.59 | 5411.54 | 3558.11 | 3435.15 | 0.77 | 0.185 |
| Planctomycetaceae | 1930.42 | 5489.80 | 9810.98 | 11363.17 | 2643.29 | 3199.28 | 0.00 | 0.998 |
| Enterobacteriaceae | 2083.24 | 3335.26 | 1507.84 | 3766.58 | 3077.55 | 2988.74 | -0.51 | 0.207 |
| Comamonadaceae | 23507.12 | 2018.79 | 5434.00 | 4098.83 | 2946.22 | 2961.35 | 1.63 | 0.405 |
| Nitrososphaeraceae | 46.84 | 8.66 | 17.53 | 5.74 | 1903.59 | 2613.63 | -5.95 | 0.197 |
| Aspergillaceae | 734.13 | 1986.51 | 3981.74 | 1152.79 | 1425.56 | 2369.75 | 0.44 | 0.611 |
| Micromonosporaceae | 1001.36 | 2142.98 | 4544.89 | 3083.15 | 1353.35 | 2350.90 | 0.18 | 0.813 |
| Caulobacteraceae | 2155.12 | 3655.85 | 2924.05 | 1490.31 | 3771.96 | 2142.16 | 0.24 | 0.616 |
| Clostridiaceae | 1447.54 | 3184.91 | 1369.64 | 3240.28 | 2470.19 | 2126.91 | -0.38 | 0.431 |
| Pseudomonadaceae | 1196.28 | 1520.51 | 1887.55 | 1455.88 | 1190.56 | 2057.16 | -0.03 | 0.924 |
| Isosphaeraceae | 548.93 | 1953.90 | 3600.86 | 1271.14 | 1362.91 | 1899.58 | 0.43 | 0.616 |
| Thermomonosporaceae | 1262.12 | 3858.69 | 1938.13 | 2107.04 | 1404.17 | 1782.81 | 0.42 | 0.532 |
| Pirellulaceae | 383.36 | 869.66 | 2186.99 | 1643.25 | 1641.18 | 1763.43 | -0.55 | 0.425 |
| Streptosporangiaceae | 1114.04 | 2809.12 | 1626.85 | 1739.58 | 1319.13 | 1665.52 | 0.23 | 0.643 |
| Rhodanobacteraceae | 3733.26 | 21528.53 | 16718.77 | 15080.27 | 3882.16 | 1665.00 | 1.03 | 0.354 |
| Cellulomonadaceae | 175.49 | 162.41 | 414.43 | 173.10 | 1017.21 | 1522.40 | -1.85 | 0.236 |
| Bacillaceae | 1262.98 | 10752.63 | 11494.07 | 3924.48 | 1198.11 | 1454.97 | 1.84 | 0.224 |
| Moraxellaceae | 761.55 | 1302.88 | 579.34 | 1648.84 | 1305.54 | 1126.62 | -0.63 | 0.154 |
| Spirosomaceae | 671.53 | 282.87 | 1660.56 | 745.47 | 968.15 | 1034.14 | -0.07 | 0.925 |
| Didymellaceae | 9.71 | 19.37 | 26.37 | 1.86 | 5.03 | 8.01 | 1.90 | 0.094 |
| Acetobacteraceae | 427.18 | 548.06 | 621.56 | 593.15 | 911.29 | 879.65 | -0.58 | 0.105 |
| Alcaligenaceae | 8360.38 | 2233.87 | 3603.86 | 3271.31 | 718.82 | 878.36 | 1.54 | 0.231 |
| Iamiaceae | 166.86 | 150.69 | 102.31 | 208.47 | 632.52 | 859.24 | -2.02 | 0.153 |
| Rhodobacteraceae | 610.01 | 824.30 | 1201.23 | 675.98 | 706.49 | 800.85 | 0.27 | 0.477 |
| Erythrobacteraceae | 458.91 | 990.11 | 1289.03 | 1205.22 | 889.15 | 766.24 | -0.06 | 0.891 |
| Micrococcaceae | 727.87 | 668.51 | 990.09 | 933.93 | 689.88 | 757.45 | 0.00 | 0.989 |
| Conexibacteraceae | 148.51 | 528.02 | 218.98 | 113.70 | 654.41 | 738.59 | -0.75 | 0.432 |
| Beijerinckiaceae | 195.14 | 228.33 | 305.27 | 207.69 | 684.35 | 730.07 | -1.15 | 0.213 |
| Frankiaceae | 385.74 | 457.34 | 435.29 | 513.42 | 586.98 | 709.40 | -0.50 | 0.076 |
| Methylobacteriaceae | 562.74 | 701.98 | 902.80 | 477.12 | 646.86 | 708.11 | 0.24 | 0.412 |
| Propionibacteriaceae | 392.21 | 797.46 | 932.34 | 727.47 | 369.85 | 671.17 | 0.26 | 0.585 |
| Nocardiopsaceae | 432.79 | 1367.44 | 873.26 | 801.15 | 499.67 | 659.03 | 0.45 | 0.477 |
| Dermacoccaceae | 139.23 | 132.51 | 197.62 | 253.14 | 274.75 | 642.49 | -1.32 | 0.203 |
| Intrasporangiaceae | 679.52 | 337.91 | 488.20 | 248.33 | 485.59 | 579.20 | 0.20 | 0.669 |
| Sphingobacteriaceae | 795.65 | 634.03 | 753.75 | 824.11 | 725.36 | 575.58 | 0.04 | 0.835 |
| Thermoactinomycetaceae | 275.43 | 1994.67 | 2486.08 | 788.43 | 270.97 | 570.42 | 1.55 | 0.257 |
| Verrucomicrobiaceae | 110.73 | 101.42 | 213.64 | 194.98 | 368.34 | 534.25 | -1.37 | 0.138 |
| Oxalobacteraceae | 2029.06 | 3438.05 | 3486.69 | 1306.97 | 818.45 | 983.76 | 1.53 | 0.046 |
| Polyangiaceae | 130.81 | 39.92 | 72.94 | 48.55 | 532.89 | 523.91 | -2.18 | 0.212 |
| Geodermatophilaceae | 327.46 | 468.89 | 377.21 | 397.09 | 425.96 | 522.11 | -0.20 | 0.366 |
| Chlorellaceae | 98.22 | 177.36 | 250.69 | 183.50 | 265.69 | 495.24 | -0.84 | 0.274 |
| Rhodospirillaceae | 246.73 | 234.11 | 364.19 | 228.94 | 463.70 | 487.75 | -0.48 | 0.314 |
| Chitinophagaceae | 472.51 | 230.88 | 285.24 | 220.41 | 464.20 | 480.51 | -0.24 | 0.626 |
| Kaistiaceae | 333.50 | 456.49 | 577.83 | 304.48 | 302.17 | 478.70 | 0.33 | 0.364 |
| Solirubrobacteraceae | 280.40 | 179.74 | 254.37 | 156.82 | 395.01 | 460.36 | -0.50 | 0.398 |
| Gordoniaceae | 111.17 | 165.98 | 157.89 | 137.74 | 269.97 | 436.08 | -0.96 | 0.253 |
| Lacipirellulaceae | 71.45 | 185.18 | 309.28 | 209.09 | 331.36 | 432.72 | -0.78 | 0.224 |
| Legionellaceae | 795.22 | 103.46 | 173.92 | 58.94 | 309.47 | 427.81 | 0.43 | 0.733 |
| Viruses | 208.09 | 390.58 | 177.76 | 412.60 | 431.49 | 413.09 | -0.70 | 0.136 |
| Roseobacteraceae | 294.21 | 409.77 | 533.77 | 338.92 | 363.31 | 409.47 | 0.16 | 0.611 |
| Kribbellaceae | 284.07 | 1228.98 | 1627.51 | 585.24 | 271.47 | 404.30 | 1.32 | 0.253 |
| Bdellovibrionaceae | 43.60 | 61.84 | 51.91 | 53.98 | 216.63 | 400.43 | -2.09 | 0.229 |
| Opitutaceae | 126.28 | 38.39 | 74.11 | 59.25 | 306.45 | 370.72 | -1.62 | 0.218 |
| Corynebacteriaceae | 214.99 | 427.10 | 229.50 | 403.76 | 379.66 | 351.60 | -0.38 | 0.327 |
| Methylocystaceae | 127.57 | 113.49 | 186.27 | 102.68 | 319.78 | 350.83 | -0.86 | 0.275 |
| Myxococcaceae | 161.25 | 94.63 | 118.84 | 118.97 | 334.12 | 339.20 | -1.08 | 0.189 |
| Capsulimonadaceae | 79.22 | 6.29 | 16.19 | 9.46 | 391.24 | 326.80 | -2.84 | 0.216 |
| Aurantimonadaceae | 325.95 | 482.32 | 664.96 | 429.66 | 273.99 | 320.34 | 0.52 | 0.265 |
| Labilitrichaceae | 41.88 | 10.87 | 14.69 | 8.84 | 271.73 | 317.50 | -3.15 | 0.206 |
| Segnochrobactraceae | 25.69 | 36.87 | 50.07 | 21.56 | 17.36 | 14.98 | 1.06 | 0.100 |
| Vibrionaceae | 221.69 | 291.19 | 200.79 | 376.46 | 316.01 | 308.46 | -0.49 | 0.054 |
| Peptoniphilaceae | 10.36 | 14.61 | 14.35 | 7.29 | 6.79 | 8.78 | 0.78 | 0.041 |
| Acidobacteriaceae | 68.64 | 72.71 | 117.67 | 78.18 | 316.76 | 279.01 | -1.38 | 0.199 |
| Kofleriaceae | 19.21 | 11.38 | 45.90 | 11.94 | 277.01 | 266.09 | -2.86 | 0.205 |
| Azospirillaceae | 189.31 | 389.22 | 250.36 | 174.66 | 268.46 | 259.63 | 0.24 | 0.571 |
| Archangiaceae | 93.25 | 57.08 | 67.43 | 62.20 | 229.46 | 252.66 | -1.32 | 0.208 |
| Flavobacteriaceae | 241.11 | 165.98 | 260.21 | 164.11 | 203.79 | 252.40 | 0.11 | 0.705 |
| Rhizopodaceae | 5692.60 | 67.62 | 1154.66 | 1071.20 | 184.92 | 234.57 | 2.21 | 0.404 |
| Sporichthyaceae | 35.40 | 29.90 | 26.87 | 23.27 | 204.30 | 232.25 | -2.32 | 0.202 |
| Aestuariivirgaceae | 329.40 | 1248.01 | 2332.86 | 997.52 | 128.82 | 226.31 | 1.53 | 0.280 |
| Peptococcaceae | 61.74 | 113.32 | 203.79 | 76.16 | 205.30 | 221.66 | -0.41 | 0.541 |
| Valsaceae | 11.66 | 12.91 | 11.68 | 19.85 | 15.60 | 19.38 | -0.60 | 0.035 |
| Cytophagaceae | 363.50 | 438.48 | 2592.40 | 493.26 | 525.09 | 202.80 | 1.47 | 0.426 |
| Enterococcaceae | 126.49 | 216.95 | 109.66 | 278.89 | 219.90 | 198.41 | -0.62 | 0.126 |
| Chthoniobacteraceae | 15.33 | 11.38 | 27.87 | 17.22 | 214.36 | 196.86 | -2.97 | 0.186 |
| Parachlamydiaceae | 72.53 | 63.20 | 94.47 | 94.93 | 239.52 | 194.79 | -1.20 | 0.139 |
| Nakamurellaceae | 87.21 | 112.13 | 138.37 | 88.10 | 153.48 | 194.53 | -0.37 | 0.413 |

| | | | | | | | | |
|---|---|---|---|---|---|---|---|---|
| Stappiaceae | 148.94 | 165.47 | 263.88 | 174.04 | 177.63 | 188.85 | 0.10 | 0.760 |
| Actinopolysporaceae | 184.13 | 157.66 | 138.70 | 145.65 | 124.54 | 184.71 | 0.08 | 0.719 |
| Staphylococcaceae | 119.59 | 216.10 | 202.63 | 311.15 | 184.67 | 182.90 | -0.33 | 0.425 |
| Sinobacteraceae | 170.31 | 63.37 | 153.89 | 103.30 | 142.91 | 175.41 | -0.12 | 0.789 |
| Promicromonosporaceae | 110.95 | 123.00 | 157.56 | 128.12 | 141.40 | 168.44 | -0.16 | 0.447 |
| Jiangellaceae | 96.27 | 167.00 | 149.38 | 118.51 | 128.82 | 167.15 | -0.01 | 0.983 |
| Sandaracinaceae | 23.74 | 6.80 | 11.68 | 8.38 | 142.66 | 166.37 | -2.91 | 0.202 |
| Kineosporiaceae | 90.88 | 119.77 | 118.34 | 92.29 | 124.04 | 161.98 | -0.20 | 0.516 |
| Chelatococcaceae | 93.90 | 115.69 | 167.24 | 92.45 | 138.13 | 160.17 | -0.05 | 0.883 |
| Selenastraceae | 26.12 | 34.32 | 21.86 | 39.40 | 42.02 | 43.66 | -0.60 | 0.048 |
| Acidothermaceae | 17.70 | 23.61 | 24.37 | 12.87 | 157.50 | 159.40 | -2.33 | 0.211 |
| Ilumatobacteraceae | 61.52 | 25.14 | 29.88 | 22.18 | 125.80 | 155.78 | -1.38 | 0.260 |
| Marinilabiliaceae | 78.36 | 107.88 | 74.11 | 153.10 | 111.96 | 130.72 | -0.60 | 0.048 |
| Sphingosinicellaceae | 103.61 | 77.81 | 99.31 | 64.84 | 151.71 | 145.45 | -0.37 | 0.438 |
| Actinospicaceae | 47.92 | 65.41 | 47.07 | 40.17 | 87.05 | 141.57 | -0.74 | 0.342 |
| Treboniaceae | 45.33 | 135.06 | 68.60 | 76.94 | 106.43 | 141.57 | -0.38 | 0.488 |
| Alicyclobacillaceae | 108.36 | 172.44 | 172.92 | 111.22 | 123.28 | 141.31 | 0.27 | 0.353 |
| Jatrophihabitantaceae | 23.31 | 36.19 | 38.72 | 22.03 | 127.81 | 139.50 | -1.56 | 0.229 |
| Actinomycetaceae | 58.93 | 99.72 | 87.79 | 86.09 | 105.92 | 135.89 | -0.41 | 0.226 |
| Pseudeurotiaceae | 8.85 | 5.27 | 5.67 | 8.07 | 10.57 | 11.88 | -0.62 | 0.088 |
| Acidimicrobiaceae | 35.83 | 18.01 | 19.53 | 14.43 | 109.19 | 130.20 | -1.79 | 0.231 |
| Patulibacteraceae | 27.85 | 24.29 | 34.88 | 17.53 | 103.66 | 128.40 | -1.52 | 0.247 |
| Coxiellaceae | 96.70 | 291.87 | 429.79 | 23.73 | 98.12 | 127.10 | 1.72 | 0.180 |
| Roseiarcaceae | 16.19 | 21.58 | 27.71 | 14.12 | 112.46 | 125.04 | -1.94 | 0.218 |
| Astrephomenaceae | 3.45 | 5.44 | 4.34 | 6.20 | 7.30 | 6.98 | -0.63 | 0.032 |
| Hymenobacteraceae | 67.13 | 111.28 | 134.86 | 84.07 | 135.61 | 117.03 | -0.10 | 0.771 |
| Pleomorphomonadaceae | 97.35 | 134.55 | 199.62 | 102.68 | 106.68 | 115.99 | 0.41 | 0.358 |
| Nitrosomonadaceae | 99.08 | 33.64 | 39.89 | 37.38 | 92.09 | 114.70 | -0.50 | 0.485 |
| Synechococcaceae | 39.50 | 65.24 | 44.23 | 77.87 | 80.26 | 74.14 | -0.64 | 0.066 |
| Nostocaceae | 51.37 | 93.78 | 95.47 | 91.36 | 96.11 | 113.15 | -0.32 | 0.302 |
| Methylophilaceae | 319.90 | 127.59 | 327.47 | 214.67 | 102.15 | 110.05 | 0.86 | 0.215 |
| Thermoanaerobacteraceae | 10.58 | 35.00 | 63.93 | 18.15 | 101.39 | 109.54 | -1.06 | 0.313 |
| Baekduiaceae | 42.31 | 85.79 | 71.10 | 31.95 | 80.51 | 109.28 | -0.15 | 0.790 |
| Halomonadaceae | 79.65 | 80.87 | 103.98 | 79.73 | 103.91 | 106.95 | -0.14 | 0.499 |
| Methylococcaceae | 91.31 | 51.14 | 88.63 | 54.29 | 99.13 | 105.14 | -0.16 | 0.681 |
| Nitrospiraceae | 87.64 | 35.17 | 60.25 | 38.47 | 92.34 | 104.11 | -0.36 | 0.534 |
| Demequinaceae | 28.92 | 24.12 | 49.57 | 31.02 | 108.44 | 103.08 | -1.24 | 0.196 |
| Verrucomicrobia subdivision 3 | 55.04 | 33.64 | 94.97 | 58.48 | 105.42 | 101.53 | -0.53 | 0.311 |
| Listeriaceae | 51.59 | 124.02 | 56.41 | 147.67 | 110.20 | 98.94 | -0.62 | 0.220 |
| Pasteurellaceae | 81.16 | 109.75 | 65.76 | 162.09 | 125.30 | 113.93 | -0.64 | 0.069 |
| Chromatiaceae | 80.73 | 34.66 | 70.94 | 49.95 | 84.03 | 96.88 | -0.31 | 0.495 |
| Trebouxiaceae | 29.14 | 45.53 | 24.03 | 45.76 | 57.36 | 54.25 | -0.67 | 0.075 |
| Cellvibrionaceae | 63.89 | 23.27 | 28.87 | 17.22 | 51.33 | 92.23 | -0.47 | 0.592 |
| Anaeromyxobacteraceae | 19.64 | 15.80 | 20.20 | 15.05 | 88.06 | 89.64 | -1.79 | 0.204 |
| Actinopolymorphaceae | 80.73 | 121.81 | 81.78 | 84.69 | 78.00 | 89.13 | 0.18 | 0.510 |
| Ectothiorhodospiraceae | 37.56 | 41.11 | 58.75 | 35.37 | 79.51 | 86.80 | -0.55 | 0.315 |
| Solibacteraceae | 9.71 | 9.34 | 16.19 | 11.48 | 73.97 | 86.29 | -2.28 | 0.187 |
| Geobacteraceae | 26.33 | 34.66 | 49.91 | 32.57 | 78.25 | 81.89 | -0.80 | 0.222 |
| Desulfobacteraceae | 38.85 | 30.41 | 48.24 | 31.95 | 81.27 | 81.89 | -0.73 | 0.254 |
| Morganellaceae | 71.45 | 97.01 | 61.09 | 113.70 | 87.30 | 81.89 | -0.30 | 0.288 |
| Glycomycetaceae | 54.18 | 82.74 | 67.93 | 87.17 | 61.39 | 79.31 | -0.15 | 0.532 |
| Sphaerobacteraceae | 19.43 | 10.02 | 16.86 | 5.27 | 49.56 | 78.28 | -1.52 | 0.305 |
| Desulfovibrionaceae | 33.03 | 34.15 | 48.57 | 38.78 | 65.42 | 76.73 | -0.64 | 0.184 |
| Leptolyngbyaceae | 30.44 | 36.70 | 50.74 | 45.60 | 74.98 | 75.69 | -0.74 | 0.102 |
| Scytonemataceae | 9.50 | 10.02 | 15.02 | 17.06 | 16.61 | 21.70 | -0.68 | 0.045 |
| Nectriaceae | 37.34 | 42.81 | 33.21 | 59.10 | 38.75 | 74.14 | -0.60 | 0.191 |
| Microascaceae | 1.73 | 1.70 | 1.00 | 2.64 | 2.26 | 2.33 | -0.71 | 0.041 |
| Oscillospiraceae | 33.46 | 76.79 | 131.02 | 49.79 | 54.85 | 73.11 | 0.44 | 0.535 |
| Nannocystaceae | 18.35 | 18.18 | 38.89 | 13.80 | 60.64 | 72.85 | -0.97 | 0.315 |
| Thalassospiraceae | 26.77 | 29.90 | 49.40 | 25.75 | 29.44 | 70.79 | -0.25 | 0.709 |
| Sclerotiniaceae | 4.32 | 2.72 | 3.17 | 5.12 | 5.28 | 6.46 | -0.72 | 0.026 |
| Miltoncostaeaceae | 7.56 | 3.91 | 7.51 | 5.43 | 34.72 | 69.24 | -2.53 | 0.243 |
| Micropepsaceae | 117.86 | 88.34 | 55.91 | 62.35 | 114.23 | 69.24 | 0.09 | 0.833 |
| Leptospiraceae | 40.37 | 22.26 | 38.05 | 25.13 | 65.67 | 68.72 | -0.66 | 0.297 |
| Catenulisporaceae | 29.14 | 50.46 | 36.39 | 38.16 | 62.40 | 68.20 | -0.54 | 0.199 |
| Deinococcaceae | 28.49 | 33.13 | 50.91 | 33.50 | 62.65 | 67.69 | -0.54 | 0.259 |
| Chromobacteriaceae | 113.76 | 37.38 | 56.58 | 39.55 | 62.40 | 66.65 | 0.30 | 0.637 |
| Cryptosporangiaceae | 34.54 | 61.67 | 54.08 | 43.12 | 51.07 | 65.36 | -0.09 | 0.781 |
| Lachnospiraceae | 30.87 | 77.81 | 118.50 | 44.21 | 56.36 | 64.84 | 0.46 | 0.504 |
| Rubrobacteraceae | 28.28 | 37.04 | 32.05 | 29.78 | 56.86 | 64.59 | -0.64 | 0.227 |
| Brevibacteriaceae | 41.23 | 69.48 | 78.78 | 59.41 | 50.07 | 64.59 | 0.12 | 0.703 |
| Parvibaculaceae | 55.26 | 37.89 | 68.60 | 37.23 | 66.17 | 64.33 | -0.05 | 0.885 |
| Rhodocyclaceae | 84.83 | 28.71 | 42.39 | 28.54 | 57.36 | 63.04 | 0.07 | 0.914 |
| Steroidobacteraceae | 25.04 | 29.73 | 39.56 | 37.23 | 59.38 | 62.78 | -0.76 | 0.095 |
| Oceanospirillaceae | 44.03 | 37.38 | 54.08 | 29.32 | 54.35 | 61.74 | -0.10 | 0.783 |
| Motilibacteraceae | 25.26 | 58.78 | 37.22 | 38.93 | 47.30 | 58.90 | -0.26 | 0.532 |
| Trichocomaceae | 15.11 | 24.97 | 37.55 | 13.03 | 33.71 | 58.13 | -0.43 | 0.578 |
| Microthrixaceae | 10.36 | 5.95 | 5.34 | 8.38 | 41.51 | 57.35 | -2.31 | 0.185 |
| Chlamydomonadaceae | 71.66 | 100.40 | 54.41 | 133.24 | 134.61 | 118.84 | -0.77 | 0.044 |
| Yersiniaceae | 1504.31 | 57.59 | 416.60 | 772.92 | 110.20 | 56.32 | 1.07 | 0.532 |
| Bathycoccaceae | 36.48 | 63.37 | 29.71 | 64.22 | 63.40 | 55.54 | -0.50 | 0.219 |
| Ophiocordycipitaceae | 7.77 | 14.61 | 8.18 | 23.58 | 17.61 | 55.03 | -1.65 | 0.197 |
| Geminigeraceae | 30.22 | 34.49 | 18.19 | 55.06 | 47.05 | 39.53 | -0.77 | 0.042 |
| Gaiellaceae | 35.83 | 4.93 | 10.85 | 4.50 | 56.36 | 53.99 | -1.15 | 0.353 |
| Bryobacteraceae | 8.42 | 5.44 | 13.85 | 11.01 | 54.85 | 53.48 | -2.11 | 0.164 |
| Alteromonadaceae | 30.22 | 29.90 | 35.22 | 27.76 | 54.09 | 51.41 | -0.48 | 0.268 |
| Pectobacteriaceae | 42.09 | 58.10 | 35.55 | 60.34 | 46.29 | 50.89 | -0.21 | 0.418 |
| Cyclobacteriaceae | 45.33 | 49.61 | 62.42 | 39.09 | 52.84 | 50.89 | 0.14 | 0.510 |
| Marinobacteraceae | 44.90 | 33.64 | 36.55 | 26.68 | 39.25 | 50.89 | -0.02 | 0.946 |
| Abitibacteriaceae | 2.37 | 3.57 | 67.60 | 35.83 | 78.75 | 50.38 | -1.17 | 0.304 |
| Zoogloeaceae | 42.09 | 28.71 | 44.40 | 29.47 | 51.83 | 50.38 | -0.19 | 0.568 |
| Lactobacillaceae | 44.25 | 69.82 | 54.58 | 48.55 | 52.84 | 49.86 | 0.16 | 0.518 |
| Ornithinimicrobiaceae | 28.28 | 42.81 | 49.74 | 31.49 | 41.77 | 49.60 | -0.02 | 0.939 |
| Tissierellaceae | 26.55 | 245.66 | 129.69 | 59.56 | 45.79 | 49.34 | 1.38 | 0.323 |
| Carbonactinosporaceae | 23.74 | 66.94 | 35.72 | 40.33 | 38.49 | 48.31 | -0.01 | 0.987 |
| Planococcaceae | 39.07 | 409.43 | 144.37 | 71.51 | 41.51 | 48.05 | 1.88 | 0.321 |
| Bogoriellaceae | 32.81 | 42.98 | 43.90 | 36.76 | 41.01 | 48.05 | -0.07 | 0.695 |
| Salinarimonadaceae | 29.79 | 25.14 | 46.40 | 21.87 | 40.76 | 47.28 | -0.12 | 0.789 |
| Phreatobacteraceae | 31.95 | 27.86 | 55.08 | 28.85 | 43.53 | 47.28 | -0.06 | 0.885 |
| Dermabacteraceae | 33.03 | 36.70 | 42.73 | 47.15 | 45.04 | 47.02 | -0.31 | 0.080 |

| | | | | | | | | |
|---|---|---|---|---|---|---|---|---|
| Caldilineaceae | 14.46 | 9.00 | 20.53 | 6.51 | 29.44 | 46.24 | -0.90 | 0.386 |
| Clavicipitaceae | 13.60 | 13.08 | 11.02 | 28.70 | 17.36 | 45.73 | -1.28 | 0.159 |
| Alsobacteraceae | 28.71 | 24.46 | 33.55 | 26.21 | 41.26 | 45.73 | -0.38 | 0.272 |
| Serendipitaceae | 4.32 | 2.89 | 2.34 | 4.65 | 6.29 | 5.43 | -0.78 | 0.042 |
| Fulvivirgaceae | 330.48 | 43.49 | 37.22 | 18.15 | 39.50 | 45.47 | 2.00 | 0.400 |
| Rhodovibrionaceae | 31.08 | 27.18 | 39.06 | 19.70 | 46.80 | 45.47 | -0.20 | 0.647 |
| Ostreobiaceae | 14.25 | 29.22 | 15.02 | 37.07 | 36.23 | 28.42 | -0.80 | 0.078 |
| Geminicoccaceae | 28.71 | 35.17 | 42.23 | 38.00 | 39.75 | 42.37 | -0.18 | 0.355 |
| Nephroselmidaceae | 1.30 | 2.38 | 2.17 | 3.57 | 4.03 | 2.84 | -0.84 | 0.033 |
| Alcanivoracaceae | 15.33 | 19.37 | 20.70 | 15.98 | 29.94 | 41.59 | -0.66 | 0.283 |
| Anaerolineaceae | 9.93 | 14.27 | 22.53 | 11.17 | 29.69 | 41.33 | -0.81 | 0.312 |
| Thalassobaculaceae | 20.72 | 19.37 | 30.71 | 18.92 | 38.75 | 40.30 | -0.47 | 0.327 |
| Amorphaceae | 20.29 | 26.16 | 34.55 | 19.85 | 32.20 | 40.30 | -0.19 | 0.632 |
| Debaryomycetaceae | 9.50 | 10.02 | 10.85 | 10.70 | 45.04 | 39.78 | -1.65 | 0.179 |
| Thermoanaerobacterales Family III. Incertae Sedis | 4.53 | 6.80 | 11.18 | 5.27 | 32.96 | 39.78 | -1.79 | 0.219 |
| Hyphomonadaceae | 28.49 | 27.69 | 37.05 | 24.51 | 37.49 | 39.78 | -0.13 | 0.644 |
| Chordaceae | 4.75 | 5.78 | 3.51 | 9.93 | 6.79 | 8.78 | -0.86 | 0.032 |
| Criblamydiaceae | 20.51 | 8.83 | 16.86 | 6.67 | 28.68 | 39.01 | -0.69 | 0.435 |
| Dermatophilaceae | 19.00 | 29.56 | 29.88 | 22.80 | 27.17 | 39.01 | -0.18 | 0.593 |
| Scenedesmaceae | 30.22 | 37.04 | 22.20 | 37.54 | 43.27 | 38.75 | -0.42 | 0.131 |
| Lichenibacteriaceae | 19.00 | 21.58 | 28.04 | 20.01 | 32.96 | 38.75 | -0.42 | 0.303 |
| Brachybasidiaceae | 0.65 | 0.51 | 0.50 | 2.33 | 16.86 | 38.49 | -5.12 | 0.218 |
| Thermaceae | 9.93 | 11.72 | 23.37 | 12.25 | 29.19 | 37.72 | -0.81 | 0.273 |
| Blastochloridaceae | 23.10 | 14.61 | 27.71 | 10.08 | 40.00 | 37.72 | -0.42 | 0.530 |
| Symbiodiniaceae | 36.26 | 26.50 | 27.54 | 23.42 | 32.46 | 37.72 | -0.05 | 0.844 |
| Weeksellaceae | 21.59 | 35.68 | 52.58 | 35.68 | 35.73 | 37.72 | 0.01 | 0.981 |
| Chloropicaceae | 19.21 | 37.38 | 18.36 | 44.05 | 37.74 | 37.20 | -0.67 | 0.130 |
| Sporomusaceae | 17.05 | 28.54 | 41.39 | 17.99 | 39.50 | 37.20 | -0.12 | 0.806 |
| Dictyobacteraceae | 16.84 | 14.95 | 18.03 | 14.58 | 31.20 | 36.68 | -0.73 | 0.242 |
| Tepidisphaeraceae | 10.58 | 14.10 | 38.89 | 30.25 | 40.51 | 36.43 | -0.75 | 0.239 |
| Ruaniaceae | 23.10 | 27.01 | 42.90 | 26.68 | 28.18 | 36.17 | 0.03 | 0.928 |
| Thiotrichaceae | 17.27 | 14.95 | 23.03 | 14.89 | 26.67 | 35.65 | -0.48 | 0.351 |
| Reyranellaceae | 135.34 | 110.60 | 186.27 | 288.66 | 205.56 | 295.80 | -0.87 | 0.034 |
| Mycosphaerellaceae | 6.91 | 7.81 | 6.51 | 14.27 | 12.08 | 12.66 | -0.88 | 0.003 |
| Peronosporaceae | 231.62 | 270.97 | 117.50 | 476.50 | 349.22 | 314.40 | -0.88 | 0.062 |
| Microcoleaceae | 13.17 | 11.89 | 20.70 | 16.13 | 33.71 | 34.62 | -0.88 | 0.153 |
| Chlamydiaceae | 17.70 | 30.24 | 65.93 | 56.77 | 34.47 | 34.10 | -0.14 | 0.829 |
| Tsukamurellaceae | 17.48 | 20.05 | 24.03 | 15.98 | 35.73 | 33.84 | -0.47 | 0.332 |
| Saprolegniaceae | 47.27 | 34.32 | 24.87 | 32.26 | 40.26 | 33.58 | 0.00 | 0.988 |
| Desulfobulbaceae | 12.09 | 15.97 | 20.53 | 17.06 | 28.93 | 33.33 | -0.71 | 0.158 |
| Puniceicoccaceae | 12.30 | 11.72 | 21.36 | 15.36 | 28.93 | 33.33 | -0.77 | 0.178 |
| Phycisphaeraceae | 9.50 | 13.93 | 24.03 | 16.91 | 27.17 | 33.33 | -0.71 | 0.197 |
| Bartonellaceae | 28.06 | 23.27 | 32.88 | 23.27 | 21.89 | 33.33 | 0.10 | 0.697 |
| Bifidobacteriaceae | 23.74 | 30.58 | 25.87 | 21.10 | 29.44 | 33.33 | -0.06 | 0.786 |
| Pycnococcaceae | 23.53 | 38.56 | 21.03 | 46.07 | 42.27 | 33.07 | -0.55 | 0.138 |
| Halieaceae | 15.97 | 17.67 | 25.20 | 16.13 | 28.68 | 32.81 | -0.40 | 0.353 |
| Shewanellaceae | 36.91 | 54.87 | 32.88 | 78.02 | 78.25 | 73.11 | -0.88 | 0.030 |
| Streptococcaceae | 29.57 | 53.18 | 51.74 | 38.31 | 24.66 | 32.29 | 0.50 | 0.226 |
| Fimbriimonadaceae | 14.46 | 3.91 | 8.85 | 4.81 | 39.00 | 32.03 | -1.48 | 0.257 |
| Maricaulaceae | 22.23 | 32.45 | 28.54 | 25.28 | 40.26 | 32.03 | -0.23 | 0.420 |
| Roseiflexaceae | 14.68 | 4.08 | 17.53 | 7.29 | 15.85 | 32.03 | -0.60 | 0.502 |
| Volvocaceae | 23.31 | 25.31 | 16.02 | 26.52 | 34.72 | 31.52 | -0.52 | 0.066 |
| Oscillatoriaceae | 10.79 | 15.29 | 16.69 | 14.27 | 27.17 | 31.52 | -0.77 | 0.183 |
| Clostridiales Family XVII. Incertae Sedis | 8.20 | 31.77 | 55.41 | 11.48 | 21.64 | 31.52 | 0.56 | 0.544 |
| Erwiniaceae | 4510.35 | 37.55 | 908.31 | 74.14 | 27.17 | 31.26 | 5.36 | 0.324 |
| Stellaceae | 19.43 | 13.42 | 24.54 | 16.60 | 25.41 | 31.00 | -0.35 | 0.383 |
| Desulfuromonadaceae | 10.15 | 13.08 | 15.52 | 10.55 | 33.97 | 30.74 | -0.96 | 0.236 |
| Vicinamibacteraceae | 10.15 | 6.12 | 14.35 | 10.86 | 29.69 | 29.45 | -1.19 | 0.159 |
| Ustilaginaceae | 5.61 | 5.10 | 2.00 | 4.65 | 24.15 | 29.19 | -2.19 | 0.179 |
| Myxotrichaceae | 3.89 | 2.72 | 0.33 | 2.48 | 25.91 | 28.68 | -3.04 | 0.180 |
| Mamiellaceae | 88.93 | 163.26 | 62.76 | 235.61 | 188.20 | 159.40 | -0.89 | 0.081 |
| Rickettsiaceae | 22.67 | 32.96 | 17.19 | 35.68 | 38.75 | 28.42 | -0.50 | 0.156 |
| Spirochaetaceae | 10.15 | 14.44 | 24.87 | 9.00 | 20.13 | 28.42 | -0.22 | 0.726 |
| Pseudoalteromonadaceae | 26.98 | 22.60 | 23.37 | 27.92 | 22.90 | 27.90 | -0.11 | 0.424 |
| Thraustochytriaceae | 58.50 | 39.92 | 39.06 | 32.26 | 29.19 | 27.64 | 0.63 | 0.120 |
| Sneathiellaceae | 12.30 | 11.89 | 24.20 | 17.06 | 25.66 | 27.64 | -0.54 | 0.233 |
| Tepidamorphaceae | 14.25 | 13.93 | 26.04 | 12.10 | 25.16 | 27.64 | -0.26 | 0.601 |
| Ktedonobacteraceae | 11.22 | 11.55 | 13.85 | 8.53 | 24.15 | 27.38 | -0.71 | 0.311 |
| Antricoccaceae | 11.01 | 32.28 | 17.19 | 17.84 | 20.63 | 26.87 | -0.11 | 0.830 |
| Lichenihabitantaceae | 9.71 | 12.40 | 17.19 | 10.08 | 20.63 | 26.61 | -0.54 | 0.345 |
| Aeromonadaceae | 22.23 | 26.33 | 29.88 | 23.73 | 22.64 | 26.35 | 0.11 | 0.497 |
| Oscillochloridaceae | 10.58 | 5.95 | 14.19 | 9.15 | 23.65 | 26.09 | -0.94 | 0.211 |
| Parvularculaceae | 15.33 | 12.91 | 18.86 | 10.24 | 23.90 | 26.09 | -0.35 | 0.477 |
| Armatimonadaceae | 8.85 | 8.15 | 22.87 | 16.75 | 32.71 | 25.83 | -0.92 | 0.151 |
| Propylenellaceae | 12.09 | 8.66 | 21.86 | 10.55 | 20.38 | 25.58 | -0.41 | 0.479 |
| Trimorphomycetaceae | 0.22 | 0.17 | 0.33 | 0.16 | 35.73 | 25.06 | -6.40 | 0.197 |
| Dietziaceae | 11.66 | 18.35 | 17.19 | 14.74 | 21.89 | 24.80 | -0.38 | 0.270 |
| Acanthamoebidae | 26.33 | 216.27 | 31.04 | 30.56 | 25.41 | 24.54 | 1.77 | 0.412 |
| Elioraeaceae | 10.58 | 11.89 | 14.52 | 9.46 | 24.41 | 24.28 | -0.65 | 0.289 |
| Heliobacteriaceae | 11.01 | 24.46 | 24.37 | 12.56 | 15.35 | 24.28 | 0.20 | 0.679 |
| Bacteroidaceae | 22.23 | 13.08 | 18.53 | 13.34 | 22.39 | 24.03 | -0.15 | 0.668 |
| Microbulbiferaceae | 14.68 | 9.00 | 16.36 | 12.72 | 20.38 | 23.77 | -0.51 | 0.238 |
| Pythiaceae | 93.90 | 25.82 | 18.53 | 25.75 | 31.45 | 23.77 | 0.77 | 0.510 |
| Cavosteliaceae | 28.06 | 76.28 | 41.39 | 64.68 | 28.18 | 23.77 | 0.32 | 0.643 |
| Vulgatibacteraceae | 4.53 | 2.55 | 3.17 | 2.33 | 20.63 | 23.51 | -2.18 | 0.210 |
| Cordycipitaceae | 6.26 | 7.48 | 6.51 | 11.32 | 8.55 | 23.51 | -1.10 | 0.235 |
| Candidatus Competibacteraceae | 15.33 | 10.02 | 15.02 | 10.86 | 29.94 | 23.51 | -0.67 | 0.288 |
| Afifellaceae | 14.46 | 13.59 | 21.03 | 11.79 | 23.40 | 23.25 | -0.25 | 0.534 |
| Neisseriaceae | 22.23 | 19.37 | 19.86 | 16.60 | 24.41 | 23.25 | -0.06 | 0.747 |
| Thermoleophilaceae | 7.99 | 1.19 | 2.67 | 0.62 | 18.37 | 22.73 | -1.82 | 0.276 |
| Piscirickettsiaceae | 15.33 | 12.06 | 15.86 | 7.45 | 22.64 | 22.22 | -0.27 | 0.611 |
| Ectocarpaceae | 31.08 | 14.61 | 8.85 | 10.86 | 23.15 | 22.22 | -0.04 | 0.947 |
| Pseudobacteriovoracaceae | 1.73 | 2.89 | 2.34 | 4.50 | 20.88 | 21.96 | -2.77 | 0.140 |
| Pseudanabaenaceae | 7.99 | 8.83 | 12.02 | 11.01 | 16.61 | 21.96 | -0.78 | 0.148 |
| Chlorobiaceae | 8.85 | 9.51 | 13.85 | 11.48 | 17.11 | 21.96 | -0.65 | 0.171 |
| Dunaliellaceae | 15.76 | 20.39 | 10.35 | 22.03 | 20.38 | 21.96 | -0.47 | 0.173 |
| Streblomastigidae | 28.28 | 33.98 | 12.02 | 55.84 | 49.06 | 34.88 | -0.91 | 0.073 |
| Sanguibacteraceae | 10.36 | 9.00 | 16.02 | 11.63 | 13.84 | 21.44 | -0.41 | 0.359 |
| Caulerpaceae | 1.30 | 1.02 | 2.50 | 2.95 | 2.26 | 3.88 | -0.92 | 0.095 |
| Longimicrobiaceae | 4.53 | 8.49 | 11.85 | 8.69 | 22.90 | 20.93 | -1.08 | 0.163 |

| | | | | | | | | |
|---|---|---|---|---|---|---|---|---|
| Syntrophaceae | 5.18 | 6.29 | 8.68 | 4.19 | 22.39 | 20.93 | -1.24 | 0.257 |
| Bradymonadaceae | 4.75 | 3.23 | 4.67 | 2.79 | 14.59 | 20.41 | -1.58 | 0.247 |
| Thalassiosiraceae | 170.53 | 10.87 | 9.01 | 9.93 | 24.66 | 20.41 | 1.79 | 0.488 |
| Plasmodiophoridae | 40.15 | 7.98 | 8.51 | 8.84 | 22.90 | 20.15 | 0.13 | 0.900 |
| Nitriliruptoraceae | 6.69 | 12.91 | 9.01 | 10.86 | 14.34 | 19.89 | -0.66 | 0.170 |
| Holophagaceae | 37.56 | 3.40 | 3.17 | 3.72 | 18.12 | 19.89 | 0.08 | 0.953 |
| Malasseziaceae | 24.82 | 20.05 | 11.18 | 32.88 | 64.91 | 19.63 | -1.07 | 0.264 |
| Prolixibacteraceae | 15.97 | 12.91 | 15.52 | 9.31 | 19.12 | 19.63 | -0.11 | 0.757 |
| Ascobolaceae | 0.86 | 1.70 | 1.00 | 2.79 | 2.26 | 1.81 | -0.95 | 0.046 |
| Glomerellaceae | 7.77 | 11.72 | 7.01 | 18.46 | 9.06 | 19.12 | -0.82 | 0.163 |
| Methanosarcinaceae | 6.04 | 7.81 | 11.85 | 5.58 | 37.99 | 19.12 | -1.29 | 0.319 |
| Gallionellaceae | 15.76 | 11.89 | 16.19 | 11.48 | 22.64 | 19.12 | -0.28 | 0.452 |
| Beutenbergiaceae | 8.63 | 10.87 | 8.51 | 6.98 | 10.82 | 19.12 | -0.40 | 0.497 |
| Azonexaceae | 27.41 | 16.31 | 18.69 | 18.77 | 19.88 | 19.12 | 0.11 | 0.691 |
| Bacteriovoracaceae | 6.48 | 5.27 | 5.17 | 3.88 | 11.83 | 18.86 | -1.03 | 0.307 |
| Rhodobiaceae | 11.22 | 9.00 | 13.02 | 6.82 | 14.34 | 18.60 | -0.26 | 0.600 |
| Sporolactobacillaceae | 24.82 | 109.58 | 63.59 | 29.32 | 16.61 | 18.34 | 1.62 | 0.208 |
| Cohaesibacteraceae | 19.21 | 23.61 | 30.88 | 16.91 | 13.33 | 18.08 | 0.61 | 0.116 |
| Kallotenuaceae | 11.66 | 5.78 | 8.68 | 6.82 | 14.34 | 18.08 | -0.59 | 0.324 |
| Thermoguttaceae | 3.89 | 5.27 | 16.19 | 9.31 | 19.62 | 17.57 | -0.88 | 0.235 |
| Bionectriaceae | 3.67 | 10.02 | 2.50 | 22.49 | 1.76 | 17.57 | -1.37 | 0.305 |
| Spongiibacteraceae | 9.71 | 6.80 | 10.18 | 6.05 | 16.61 | 17.57 | -0.59 | 0.346 |
| Zavarziniaceae | 9.50 | 5.78 | 10.18 | 5.74 | 13.59 | 17.57 | -0.54 | 0.393 |
| Ancalomicrobiaceae | 14.46 | 16.99 | 21.36 | 10.08 | 21.13 | 17.57 | 0.11 | 0.747 |
| Simkaniaceae | 8.63 | 5.27 | 7.01 | 4.34 | 14.34 | 17.31 | -0.78 | 0.328 |
| Trypanosomatidae | 19.43 | 14.78 | 12.85 | 16.13 | 39.25 | 17.05 | -0.62 | 0.380 |
| Selenomonadaceae | 5.40 | 8.49 | 15.86 | 5.27 | 13.59 | 17.05 | -0.27 | 0.683 |
| Mucoraceae | 11.22 | 23.61 | 20.53 | 20.32 | 13.84 | 17.05 | 0.11 | 0.761 |
| Chaetomiaceae | 29.36 | 7.48 | 6.01 | 12.87 | 7.30 | 17.05 | 0.20 | 0.834 |
| Syntrophobacteraceae | 4.10 | 4.42 | 11.85 | 8.69 | 16.61 | 16.79 | -1.05 | 0.121 |
| Desulfallaceae | 4.10 | 8.83 | 19.53 | 5.89 | 16.86 | 16.79 | -0.28 | 0.708 |
| Sterolibacteriaceae | 18.78 | 10.70 | 13.19 | 7.76 | 19.88 | 16.79 | -0.06 | 0.901 |
| Acidithiobacillaceae | 7.12 | 8.32 | 12.35 | 7.76 | 13.33 | 16.53 | -0.44 | 0.349 |
| Peptostreptococcaceae | 5.61 | 20.22 | 35.89 | 6.82 | 12.33 | 16.53 | 0.79 | 0.430 |
| Vallicoccaceae | 7.56 | 17.84 | 10.85 | 11.79 | 15.10 | 16.53 | -0.26 | 0.529 |
| Herpotrichiellaceae | 10.15 | 5.95 | 12.02 | 8.53 | 22.39 | 16.28 | -0.75 | 0.251 |
| Veillonellaceae | 5.18 | 10.19 | 14.52 | 9.15 | 14.09 | 16.02 | -0.39 | 0.412 |
| Bacillariaceae | 28.06 | 7.65 | 8.35 | 8.07 | 18.87 | 16.02 | 0.04 | 0.964 |
| Syntrophomonadaceae | 4.32 | 9.17 | 24.87 | 6.05 | 16.35 | 16.02 | 0.00 | 0.998 |
| Candidatus Brocadiaceae | 7.12 | 5.10 | 8.18 | 5.12 | 14.59 | 15.76 | -0.80 | 0.272 |
| Balneolaceae | 8.42 | 13.42 | 14.19 | 8.07 | 16.86 | 15.76 | -0.18 | 0.667 |
| Sulfuricellaceae | 10.36 | 6.29 | 10.68 | 8.69 | 12.83 | 15.50 | -0.44 | 0.262 |
| Breoghaniaceae | 13.38 | 13.08 | 18.69 | 8.07 | 18.87 | 15.24 | 0.10 | 0.803 |
| Dacrymycetaceae | 37.99 | 64.22 | 26.04 | 101.13 | 77.49 | 70.27 | -0.96 | 0.053 |
| Hapalosiphonaceae | 6.04 | 6.63 | 7.18 | 9.00 | 10.82 | 14.98 | -0.81 | 0.102 |
| Calotrichaceae | 5.18 | 7.81 | 9.18 | 8.53 | 15.60 | 14.73 | -0.81 | 0.113 |
| Colwelliaceae | 8.85 | 4.93 | 9.51 | 5.89 | 10.06 | 14.73 | -0.40 | 0.459 |
| Prevotellaceae | 7.56 | 6.46 | 8.35 | 9.31 | 10.06 | 14.47 | -0.60 | 0.129 |
| Cryptobasidiaceae | 0.86 | 0.68 | 1.34 | 0.93 | 5.79 | 14.47 | -2.88 | 0.263 |
| Treponemataceae | 5.40 | 6.97 | 9.01 | 5.12 | 10.06 | 14.47 | -0.47 | 0.421 |
| Flammeovirgaceae | 24.61 | 16.99 | 17.19 | 10.24 | 17.11 | 14.21 | 0.50 | 0.151 |
| Chthonomonadaceae | 3.02 | 2.38 | 5.67 | 3.41 | 14.59 | 14.21 | -1.54 | 0.188 |
| Neocallimastigaceae | 14.46 | 14.44 | 8.01 | 17.06 | 15.35 | 14.21 | -0.34 | 0.268 |
| Coleofasciculaceae | 13.60 | 8.32 | 10.18 | 9.46 | 15.60 | 14.21 | -0.29 | 0.381 |
| Tetrahymenidae | 11.01 | 15.12 | 8.18 | 12.10 | 13.33 | 14.21 | -0.21 | 0.475 |
| Robiginitomaculaceae | 10.58 | 6.46 | 13.52 | 7.60 | 15.60 | 14.21 | -0.29 | 0.517 |
| Gloeobacteraceae | 4.53 | 2.72 | 6.68 | 2.79 | 17.86 | 13.95 | -1.31 | 0.263 |
| Dictyosteliaceae | 14.25 | 20.90 | 17.36 | 18.92 | 14.34 | 13.95 | 0.15 | 0.521 |
| Pseudoxanthobacteraceae | 10.58 | 13.08 | 22.53 | 12.56 | 12.83 | 13.95 | 0.23 | 0.596 |
| Haematococcaceae | 6.48 | 13.25 | 5.34 | 11.32 | 13.59 | 13.69 | -0.62 | 0.203 |
| Polyporaceae | 12.52 | 24.46 | 20.53 | 13.96 | 13.59 | 13.69 | 0.48 | 0.263 |
| Synergistaceae | 4.10 | 6.12 | 11.02 | 5.27 | 11.07 | 13.69 | -0.50 | 0.416 |
| Salpingoecidae | 18.78 | 15.12 | 9.51 | 15.36 | 15.35 | 13.43 | -0.02 | 0.938 |
| Euzebyaceae | 6.26 | 7.48 | 10.68 | 8.22 | 12.83 | 13.18 | -0.49 | 0.192 |
| Ophiostomataceae | 4.53 | 6.29 | 3.17 | 9.62 | 4.28 | 13.18 | -0.95 | 0.228 |
| Waddliaceae | 6.48 | 4.59 | 6.01 | 2.79 | 12.58 | 13.18 | -0.74 | 0.374 |
| Cyanidiaceae | 19.86 | 14.44 | 11.02 | 20.78 | 18.62 | 13.18 | -0.21 | 0.520 |
| Salinisphaeraceae | 9.07 | 9.00 | 10.18 | 6.82 | 12.08 | 13.18 | -0.18 | 0.584 |
| Egicoccaceae | 5.61 | 23.95 | 12.52 | 14.58 | 10.57 | 13.18 | 0.14 | 0.838 |
| Wenzhouxiangellaceae | 6.91 | 5.95 | 9.51 | 4.65 | 13.59 | 12.92 | -0.48 | 0.421 |
| Physalacriaceae | 7.77 | 14.10 | 8.35 | 11.79 | 9.31 | 12.92 | -0.17 | 0.618 |
| Eubacteriaceae | 5.18 | 14.61 | 16.86 | 6.67 | 10.82 | 12.92 | 0.27 | 0.641 |
| Idiomarinaceae | 9.93 | 9.00 | 11.52 | 4.81 | 13.59 | 12.92 | -0.04 | 0.930 |
| Ceratobasidiaceae | 52.67 | 84.43 | 35.05 | 140.07 | 101.14 | 98.17 | -0.98 | 0.048 |
| Prochlorococcaceae | 4.75 | 4.93 | 6.84 | 7.76 | 12.33 | 12.66 | -0.99 | 0.059 |
| Aerococcaceae | 4.53 | 14.61 | 10.85 | 8.22 | 11.07 | 12.66 | -0.09 | 0.853 |
| Coriobacteriaceae | 3.02 | 5.27 | 6.84 | 4.65 | 10.82 | 12.40 | -0.88 | 0.207 |
| Lewinellaceae | 6.91 | 8.32 | 8.01 | 6.36 | 8.05 | 12.40 | -0.21 | 0.580 |
| Rhodothermaceae | 5.18 | 4.59 | 7.84 | 4.50 | 9.56 | 12.14 | -0.57 | 0.335 |
| Reticulibacteraceae | 9.71 | 4.93 | 6.18 | 5.58 | 11.32 | 12.14 | -0.48 | 0.343 |
| Campylobacteraceae | 6.69 | 8.66 | 11.02 | 6.67 | 12.83 | 12.14 | -0.26 | 0.497 |
| Synchytriaceae | 16.19 | 11.04 | 5.67 | 9.31 | 12.33 | 12.14 | -0.04 | 0.935 |
| Crocinitomicaceae | 70.37 | 76.45 | 39.89 | 133.09 | 144.92 | 93.00 | -0.99 | 0.039 |
| Desulfocapsaceae | 3.24 | 6.80 | 7.34 | 6.67 | 12.33 | 11.63 | -0.82 | 0.122 |
| Acytosteliaceae | 17.48 | 19.88 | 14.52 | 16.44 | 11.07 | 11.63 | 0.41 | 0.140 |
| Thiobacillaceae | 7.34 | 5.44 | 8.01 | 6.05 | 10.82 | 11.63 | -0.45 | 0.277 |
| Halobacteroidaceae | 2.81 | 13.42 | 16.86 | 6.82 | 4.53 | 11.63 | 0.53 | 0.528 |
| Cryptococcaceae | 16.41 | 30.07 | 14.52 | 51.65 | 36.23 | 35.13 | -1.01 | 0.047 |
| Thermogemmatisporaceae | 5.18 | 6.97 | 7.84 | 6.36 | 10.57 | 11.37 | -0.50 | 0.211 |
| Chytriomycetaceae | 10.58 | 11.21 | 7.01 | 7.60 | 12.33 | 11.37 | -0.12 | 0.691 |
| Trueperaceae | 4.75 | 3.57 | 15.36 | 3.10 | 7.30 | 11.37 | 0.12 | 0.894 |
| Plectosphaerellaceae | 6.48 | 6.63 | 3.17 | 12.56 | 6.29 | 11.11 | -0.88 | 0.123 |
| Orbiliaceae | 3.67 | 4.76 | 3.34 | 2.79 | 8.81 | 11.11 | -0.95 | 0.278 |
| Candidatus Paracaedibacteraceae | 8.63 | 1.36 | 3.84 | 3.57 | 19.37 | 11.11 | -1.30 | 0.278 |
| Ktedonosporobacteraceae | 3.24 | 4.76 | 5.17 | 2.02 | 6.29 | 11.11 | -0.56 | 0.514 |
| Eggerthellaceae | 3.45 | 6.12 | 6.51 | 1.40 | 7.80 | 11.11 | -0.34 | 0.678 |
| Geosiphonaceae | 17.92 | 3.06 | 2.67 | 2.64 | 7.30 | 11.11 | 0.17 | 0.887 |
| Ajellomycetaceae | 4.32 | 7.48 | 11.68 | 5.12 | 6.29 | 11.11 | 0.06 | 0.915 |
| Porphyridiaceae | 6.69 | 8.66 | 5.67 | 8.53 | 10.32 | 10.85 | -0.50 | 0.065 |
| Thermodesulfobacteriaceae | 1.73 | 3.40 | 4.84 | 1.71 | 8.05 | 10.85 | -1.05 | 0.320 |

| | | | | | | | | |
|---|---|---|---|---|---|---|---|---|
| Spizellomycetaceae | 16.19 | 4.25 | 2.84 | 3.41 | 9.81 | 10.85 | -0.05 | 0.959 |
| Monodopsidaceae | 9.71 | 8.32 | 7.51 | 9.93 | 11.32 | 10.59 | -0.32 | 0.062 |
| Pelomyxidae | 13.17 | 17.84 | 8.18 | 11.79 | 10.57 | 10.59 | 0.25 | 0.535 |
| Microcystaceae | 4.10 | 6.12 | 5.67 | 5.27 | 9.06 | 10.33 | -0.63 | 0.185 |
| Aphanothecaceae | 5.40 | 5.27 | 8.51 | 4.65 | 13.08 | 10.33 | -0.55 | 0.360 |
| Terasakiellaceae | 7.56 | 4.93 | 7.51 | 2.95 | 13.59 | 10.33 | -0.43 | 0.547 |
| Pavlovaceae | 12.95 | 11.89 | 8.01 | 9.93 | 12.33 | 10.33 | 0.01 | 0.961 |
| Oxytrichidae | 26.33 | 23.44 | 11.35 | 38.16 | 43.78 | 41.85 | -1.02 | 0.033 |
| Legeriomycetaceae | 8.85 | 9.68 | 5.17 | 10.55 | 11.83 | 10.08 | -0.45 | 0.159 |
| Amorphothecaceae | 0.22 | 0.17 | 0.00 | 0.78 | 7.55 | 10.08 | -5.58 | 0.163 |
| Deferribacteraceae | 3.45 | 3.40 | 5.84 | 2.95 | 6.29 | 10.08 | -0.61 | 0.402 |
| Erysipelotrichaceae | 3.02 | 9.85 | 6.84 | 7.45 | 6.79 | 10.08 | -0.30 | 0.540 |
| Helicobacteraceae | 3.02 | 6.12 | 8.68 | 4.50 | 5.79 | 10.08 | -0.19 | 0.737 |
| Teratosphaeriaceae | 6.91 | 9.85 | 7.01 | 12.87 | 9.81 | 9.82 | -0.45 | 0.107 |
| Aphanizomenonaceae | 6.48 | 4.76 | 6.34 | 6.82 | 7.04 | 9.82 | -0.43 | 0.158 |
| Nitrospinaceae | 3.02 | 3.40 | 7.01 | 3.72 | 12.33 | 9.82 | -0.95 | 0.244 |
| Tribonemataceae | 12.30 | 11.89 | 6.68 | 6.98 | 9.31 | 9.82 | 0.24 | 0.489 |
| Thermoascaceae | 9.93 | 9.00 | 16.02 | 15.67 | 8.05 | 9.82 | 0.06 | 0.889 |
| Umbelopsidaceae | 6.04 | 7.65 | 4.34 | 6.98 | 12.08 | 9.56 | -0.67 | 0.126 |
| Ardenticatenaceae | 2.37 | 1.87 | 4.34 | 2.02 | 11.32 | 9.56 | -1.42 | 0.232 |
| Prochlorotrichaceae | 5.40 | 3.06 | 6.68 | 4.19 | 6.79 | 9.56 | -0.44 | 0.398 |
| Vahlkampfiidae | 15.33 | 11.38 | 10.35 | 13.65 | 9.56 | 9.56 | 0.18 | 0.522 |
| Reichenbachiellaceae | 7.12 | 9.85 | 7.34 | 4.03 | 7.80 | 9.56 | 0.19 | 0.633 |
| Fomitopsidaceae | 4.53 | 6.12 | 7.18 | 3.26 | 6.79 | 9.56 | -0.14 | 0.786 |
| Chrysochromulinaceae | 10.58 | 6.29 | 6.34 | 6.67 | 8.30 | 9.56 | -0.08 | 0.805 |
| Kiloniellaceae | 9.07 | 7.65 | 9.35 | 4.65 | 11.07 | 9.56 | 0.04 | 0.908 |
| Erysiphaceae | 12.74 | 15.63 | 15.52 | 32.88 | 21.64 | 34.88 | -1.03 | 0.060 |
| Hypoxylaceae | 3.24 | 3.91 | 2.67 | 6.36 | 3.02 | 9.30 | -0.93 | 0.242 |
| Natrialbaceae | 4.53 | 4.42 | 4.17 | 4.03 | 6.29 | 9.30 | -0.58 | 0.291 |
| Pelagibacteraceae | 4.96 | 2.21 | 5.17 | 2.95 | 7.30 | 9.30 | -0.66 | 0.338 |
| Acidiferrobacteraceae | 5.40 | 4.42 | 6.01 | 2.48 | 9.56 | 9.30 | -0.43 | 0.512 |
| Gigasporaceae | 10.58 | 6.12 | 5.67 | 6.98 | 8.05 | 9.30 | -0.12 | 0.729 |
| Saccotheciaceae | 3.89 | 13.59 | 9.68 | 10.24 | 10.32 | 9.30 | -0.14 | 0.781 |
| Pichiaceae | 8.85 | 6.12 | 5.17 | 7.91 | 8.05 | 9.04 | -0.31 | 0.276 |
| Syntrophorhabdaceae | 4.32 | 2.04 | 4.01 | 1.55 | 8.05 | 9.04 | -0.85 | 0.362 |
| Halothiobacillaceae | 3.02 | 3.23 | 4.17 | 2.02 | 5.28 | 9.04 | -0.65 | 0.434 |
| Ahrensiaceae | 7.34 | 6.63 | 14.52 | 7.91 | 8.30 | 9.04 | 0.17 | 0.712 |
| Theileriidae | 29.14 | 32.79 | 16.36 | 60.65 | 54.09 | 45.47 | -1.03 | 0.015 |
| Glomeraceae | 294.86 | 436.79 | 182.76 | 780.37 | 605.60 | 523.91 | -1.06 | 0.035 |
| Acaulosporaceae | 5.61 | 8.49 | 2.67 | 9.62 | 7.55 | 8.78 | -0.63 | 0.203 |
| Porticoccaceae | 4.75 | 2.55 | 3.00 | 2.95 | 7.55 | 8.78 | -0.90 | 0.228 |
| Herpetosiphonaceae | 1.51 | 3.74 | 4.84 | 2.48 | 6.29 | 8.78 | -0.80 | 0.316 |
| Perkinsidae | 9.28 | 7.31 | 5.17 | 8.53 | 6.79 | 8.78 | -0.15 | 0.600 |
| Agaricaceae | 7.12 | 12.57 | 6.01 | 13.03 | 7.80 | 8.78 | -0.20 | 0.642 |
| Halobacteriaceae | 3.45 | 6.29 | 5.84 | 3.26 | 6.04 | 8.78 | -0.21 | 0.678 |
| Vitrellaceae | 12.09 | 6.29 | 6.18 | 7.45 | 10.82 | 8.78 | -0.14 | 0.729 |
| Candidatus Ozemobacteraceae | 0.43 | 1.02 | 1.34 | 2.02 | 2.01 | 1.81 | -1.07 | 0.054 |
| Kordiimonadaceae | 4.32 | 4.93 | 5.01 | 5.58 | 8.81 | 8.53 | -0.69 | 0.101 |
| Methanotrichaceae | 1.51 | 1.87 | 2.17 | 1.24 | 6.29 | 8.53 | -1.53 | 0.245 |
| Marinifilaceae | 6.04 | 8.83 | 8.35 | 8.38 | 10.57 | 8.53 | -0.24 | 0.275 |
| Sphaerobolaceae | 6.04 | 14.27 | 6.51 | 15.51 | 13.08 | 8.53 | -0.47 | 0.369 |
| Haloarculaceae | 2.81 | 7.14 | 6.34 | 6.51 | 5.54 | 8.53 | -0.34 | 0.428 |
| Caedimonadaceae | 7.34 | 1.19 | 3.34 | 1.24 | 6.79 | 8.53 | -0.48 | 0.612 |
| Coprobacillaceae | 4.32 | 9.00 | 6.51 | 6.20 | 2.77 | 8.53 | 0.18 | 0.737 |
| Terrimicrobiaceae | 0.43 | 0.17 | 1.50 | 1.24 | 5.28 | 8.27 | -2.81 | 0.169 |
| Calditrichaceae | 2.37 | 2.21 | 3.51 | 2.33 | 4.78 | 8.27 | -0.93 | 0.292 |
| Pyrinomonadaceae | 2.81 | 1.02 | 5.34 | 2.02 | 4.78 | 8.27 | -0.72 | 0.428 |
| Psathyrellaceae | 4.96 | 12.40 | 3.51 | 10.24 | 9.56 | 8.27 | -0.43 | 0.478 |
| Methanobacteriaceae | 3.45 | 3.91 | 11.52 | 2.17 | 10.57 | 8.27 | -0.15 | 0.854 |
| Coniochaetaceae | 6.04 | 3.06 | 1.50 | 7.91 | 5.79 | 8.53 | -1.07 | 0.081 |
| Desulfarculaceae | 1.51 | 2.38 | 3.67 | 3.10 | 5.54 | 8.01 | -1.14 | 0.154 |
| Tulasnellaceae | 3.89 | 5.44 | 3.84 | 4.50 | 8.05 | 8.01 | -0.64 | 0.159 |
| Jonesiaceae | 6.48 | 9.34 | 7.51 | 4.50 | 6.04 | 8.01 | 0.33 | 0.295 |
| Egibacteraceae | 4.10 | 4.08 | 3.00 | 3.10 | 6.04 | 8.01 | -0.62 | 0.296 |
| Carnobacteriaceae | 4.96 | 37.72 | 21.70 | 11.32 | 6.79 | 8.01 | 1.30 | 0.309 |
| Silvanigrellaceae | 3.67 | 2.21 | 3.84 | 1.86 | 6.54 | 8.01 | -0.76 | 0.352 |
| Rhabdaerophilaceae | 3.02 | 4.08 | 9.01 | 4.81 | 8.05 | 8.01 | -0.37 | 0.509 |
| Acidaminococcaceae | 3.89 | 9.51 | 8.85 | 4.65 | 5.79 | 8.01 | 0.27 | 0.576 |
| Noelaerhabdaceae | 14.03 | 9.68 | 8.18 | 9.93 | 12.33 | 8.01 | 0.08 | 0.815 |
| Saccharomycetaceae | 7.99 | 9.00 | 5.17 | 8.38 | 4.78 | 8.01 | 0.07 | 0.847 |
| Saccharospirillaceae | 6.69 | 6.80 | 10.01 | 3.26 | 7.55 | 7.75 | 0.34 | 0.422 |
| Chloroflexaceae | 2.37 | 2.38 | 5.84 | 0.93 | 7.04 | 7.75 | -0.57 | 0.535 |
| Blastocystidae | 11.22 | 7.81 | 6.34 | 8.22 | 6.29 | 7.75 | 0.19 | 0.558 |
| Tetrabaenaceae | 2.59 | 4.76 | 2.50 | 3.72 | 8.05 | 7.49 | -0.97 | 0.133 |
| Lipomycetaceae | 7.12 | 10.53 | 11.52 | 8.22 | 6.29 | 7.49 | 0.41 | 0.207 |
| Halanaerobiaceae | 4.32 | 51.99 | 45.90 | 36.14 | 4.28 | 7.49 | 1.09 | 0.381 |
| Patellariaceae | 2.37 | 6.29 | 15.86 | 4.50 | 2.52 | 7.49 | 0.76 | 0.500 |
| Suillaceae | 9.71 | 15.29 | 7.01 | 17.37 | 11.57 | 7.49 | -0.19 | 0.716 |
| Sordariaceae | 11.44 | 3.40 | 2.67 | 6.36 | 6.29 | 7.49 | -0.20 | 0.785 |
| Diversisporaceae | 29.57 | 40.60 | 16.69 | 72.75 | 56.86 | 56.58 | -1.10 | 0.022 |
| Bangiaceae | 5.18 | 4.93 | 3.67 | 5.58 | 6.79 | 6.98 | -0.49 | 0.044 |
| Halteriidae | 2.37 | 4.08 | 3.34 | 3.41 | 8.30 | 6.98 | -0.93 | 0.170 |
| Halobacteriovoraceae | 2.16 | 2.55 | 2.84 | 2.17 | 7.55 | 6.98 | -1.15 | 0.214 |
| Methanocellaceae | 1.30 | 0.68 | 2.34 | 0.78 | 7.80 | 6.98 | -1.85 | 0.230 |
| Hyaloscyphaceae | 1.73 | 1.19 | 1.50 | 1.09 | 5.28 | 6.98 | -1.59 | 0.231 |
| Xylariaceae | 3.89 | 3.57 | 3.84 | 8.53 | 3.27 | 6.98 | -0.73 | 0.251 |
| Rikenellaceae | 3.24 | 3.57 | 4.84 | 3.26 | 7.04 | 6.98 | -0.57 | 0.270 |
| Paraglomeraceae | 3.24 | 5.27 | 3.34 | 3.57 | 5.03 | 6.98 | -0.40 | 0.362 |
| Cavenderiaceae | 6.91 | 8.83 | 6.51 | 7.14 | 5.79 | 6.98 | 0.16 | 0.411 |
| Botryosphaeriaceae | 37.78 | 4.25 | 9.18 | 7.91 | 4.78 | 6.98 | 1.38 | 0.421 |
| Anaplasmataceae | 5.18 | 2.21 | 2.00 | 0.78 | 6.04 | 6.98 | -0.55 | 0.550 |
| Francisellaceae | 5.61 | 1.19 | 3.17 | 1.09 | 5.28 | 6.98 | -0.42 | 0.634 |
| Porphyromonadaceae | 3.89 | 5.10 | 6.34 | 5.27 | 8.81 | 6.72 | -0.44 | 0.226 |
| Desulfomicrobiaceae | 1.51 | 2.21 | 4.84 | 2.64 | 6.04 | 6.72 | -0.85 | 0.235 |
| Rhabdochlamydiaceae | 2.59 | 1.53 | 4.17 | 1.86 | 6.29 | 6.72 | -0.84 | 0.297 |
| Haliscomenobacteraceae | 2.81 | 2.72 | 6.68 | 3.41 | 5.03 | 6.72 | -0.31 | 0.577 |
| Metschnikowiaceae | 6.04 | 10.87 | 5.84 | 9.31 | 9.56 | 6.72 | -0.17 | 0.650 |
| Pucciniaceae | 4.10 | 6.46 | 10.85 | 6.20 | 5.54 | 6.72 | 0.21 | 0.670 |
| Fusobacteriaceae | 3.67 | 6.12 | 5.17 | 2.48 | 3.77 | 6.72 | 0.21 | 0.675 |
| Mortierellaceae | 53.96 | 70.67 | 41.23 | 107.18 | 111.21 | 147.51 | -1.14 | 0.017 |

| | | | | | | | | |
|---|---|---|---|---|---|---|---|---|
| Candidatus Nanopelagicaceae | 1.30 | 2.55 | 1.84 | 3.10 | 4.03 | 6.46 | -1.26 | 0.106 |
| Hymenochaetaceae | 5.83 | 6.80 | 4.01 | 9.15 | 8.05 | 6.46 | -0.51 | 0.107 |
| Merismopediaceae | 1.51 | 3.40 | 3.84 | 3.41 | 7.04 | 6.46 | -0.95 | 0.123 |
| Hahellaceae | 5.83 | 4.08 | 3.17 | 4.03 | 7.04 | 6.46 | -0.42 | 0.287 |
| Thermosporotrichaceae | 2.81 | 3.91 | 5.17 | 2.64 | 6.79 | 6.46 | -0.42 | 0.439 |
| Pneumocystidaceae | 7.12 | 4.25 | 2.50 | 6.05 | 4.28 | 6.46 | -0.27 | 0.566 |
| Clostridiales Family XVI. Incertae Sedis | 2.81 | 4.93 | 10.35 | 4.03 | 3.52 | 6.46 | 0.37 | 0.620 |
| Minwuiaceae | 4.75 | 5.27 | 5.84 | 2.33 | 5.28 | 6.46 | 0.17 | 0.680 |
| Ferrimonadaceae | 5.61 | 3.23 | 3.67 | 2.64 | 5.03 | 6.46 | -0.18 | 0.710 |
| Reticulomyxidae | 12.52 | 4.76 | 2.84 | 2.33 | 8.05 | 6.46 | 0.26 | 0.769 |
| Akkermansiaceae | 2.59 | 3.74 | 5.17 | 2.02 | 4.28 | 6.46 | -0.15 | 0.796 |
| Parameciidae | 5.83 | 4.76 | 4.34 | 4.03 | 4.78 | 6.46 | -0.03 | 0.899 |
| Notoacmeibacteraceae | 4.75 | 4.08 | 7.68 | 4.81 | 4.78 | 6.46 | 0.04 | 0.910 |
| Naemateliaceae | 9.93 | 0.51 | 0.83 | 0.31 | 5.79 | 6.46 | -0.16 | 0.913 |
| Usitatibacteraceae | 3.89 | 2.21 | 3.67 | 1.71 | 8.81 | 6.20 | -0.78 | 0.381 |
| Endozoicomonadaceae | 3.89 | 3.40 | 5.67 | 3.10 | 8.05 | 6.20 | -0.42 | 0.430 |
| Sarocladiaceae | 0.86 | 35.68 | 0.67 | 10.24 | 0.75 | 6.20 | 1.11 | 0.628 |
| Basidiobolaceae | 6.69 | 7.14 | 5.51 | 6.51 | 5.79 | 6.20 | 0.06 | 0.639 |
| Powellomycetaceae | 5.61 | 4.25 | 2.50 | 5.89 | 7.30 | 5.94 | -0.63 | 0.112 |
| Acanthopleuribacteraceae | 2.37 | 1.02 | 2.50 | 2.17 | 6.29 | 5.94 | -1.29 | 0.154 |
| Desulfohalobiaceae | 2.59 | 1.87 | 4.17 | 2.33 | 6.29 | 5.94 | -0.75 | 0.261 |
| Methanomicrobiaceae | 2.37 | 1.87 | 2.17 | 1.40 | 4.78 | 5.94 | -0.92 | 0.297 |
| Hyellaceae | 3.24 | 2.21 | 3.34 | 2.64 | 3.77 | 5.94 | -0.49 | 0.347 |
| Saprospiraceae | 4.53 | 5.61 | 8.01 | 5.89 | 8.81 | 5.94 | -0.19 | 0.587 |
| Psychromonadaceae | 3.67 | 2.89 | 8.85 | 3.41 | 3.02 | 5.94 | 0.32 | 0.662 |
| Eubacteriales Family XIII. Incertae Sedis | 1.73 | 6.63 | 6.01 | 3.57 | 6.79 | 5.94 | -0.18 | 0.743 |
| Albuginaceae | 9.93 | 8.15 | 5.34 | 8.22 | 7.80 | 5.94 | 0.09 | 0.767 |
| Stentoridae | 7.77 | 6.63 | 4.17 | 5.74 | 7.80 | 5.94 | -0.07 | 0.822 |
| Thermoflexaceae | 1.94 | 2.04 | 3.51 | 1.71 | 1.01 | 5.94 | -0.21 | 0.829 |
| Ulvaceae | 4.96 | 4.76 | 5.01 | 7.45 | 7.04 | 5.68 | -0.45 | 0.074 |
| Mollisiaceae | 0.22 | 1.70 | 1.84 | 2.02 | 4.53 | 5.68 | -1.70 | 0.104 |
| Tannerellaceae | 4.10 | 4.25 | 4.51 | 4.50 | 5.03 | 5.68 | -0.24 | 0.138 |
| Piptocephalidaceae | 4.53 | 4.25 | 2.84 | 5.43 | 4.03 | 5.68 | -0.38 | 0.186 |
| Nitrosopumilaceae | 2.16 | 1.36 | 1.84 | 1.40 | 7.30 | 5.68 | -1.42 | 0.228 |
| Raperosteliaceae | 7.34 | 7.65 | 5.67 | 6.82 | 4.78 | 5.68 | 0.26 | 0.257 |
| Cryomorphaceae | 2.59 | 2.89 | 4.01 | 3.57 | 3.52 | 5.68 | -0.43 | 0.272 |
| Atopobiaceae | 1.08 | 1.87 | 3.34 | 1.24 | 6.29 | 5.68 | -1.07 | 0.283 |
| Cunninghamellaceae | 4.32 | 6.46 | 3.34 | 6.20 | 6.04 | 5.68 | -0.35 | 0.300 |
| Halorubraceae | 1.51 | 3.57 | 3.17 | 1.55 | 5.28 | 5.68 | -0.60 | 0.404 |
| Sporocadaceae | 2.81 | 4.76 | 2.67 | 7.29 | 2.01 | 5.68 | -0.55 | 0.427 |
| Arcobacteraceae | 3.02 | 2.89 | 3.34 | 3.10 | 3.02 | 5.68 | -0.35 | 0.433 |
| Lachnaceae | 5.61 | 2.55 | 3.67 | 2.48 | 8.30 | 5.68 | -0.48 | 0.476 |
| Endogonaceae | 5.18 | 3.40 | 3.00 | 4.03 | 3.77 | 5.68 | -0.22 | 0.518 |
| Chroococcaceae | 2.37 | 2.04 | 4.01 | 0.93 | 4.53 | 5.68 | -0.40 | 0.604 |
| Segniliparaceae | 2.81 | 4.42 | 4.17 | 1.71 | 5.79 | 5.68 | -0.21 | 0.711 |
| Arthrodermataceae | 1.73 | 3.40 | 6.01 | 3.26 | 3.27 | 5.68 | -0.13 | 0.823 |
| Thermoproteaceae | 1.08 | 1.02 | 0.33 | 1.24 | 2.01 | 2.33 | -1.20 | 0.064 |
| Phaffomycetaceae | 4.32 | 3.40 | 3.00 | 4.19 | 6.29 | 5.43 | -0.57 | 0.087 |
| Fastidiosibacteraceae | 3.45 | 1.53 | 2.67 | 2.48 | 5.28 | 5.43 | -0.79 | 0.189 |
| Rarobacteraceae | 2.16 | 2.04 | 2.84 | 2.02 | 7.04 | 5.43 | -1.04 | 0.234 |
| Oleiphilaceae | 3.45 | 1.02 | 2.50 | 1.09 | 5.79 | 5.43 | -0.82 | 0.370 |
| Gigartinaceae | 3.89 | 6.12 | 2.84 | 4.65 | 5.03 | 5.43 | -0.23 | 0.518 |
| Sutterellaceae | 10.79 | 3.40 | 8.18 | 5.12 | 6.79 | 5.43 | 0.37 | 0.523 |
| Microthyriaceae | 4.10 | 1.36 | 0.50 | 0.62 | 3.52 | 5.43 | -0.68 | 0.536 |
| Pleioneaceae | 3.02 | 1.53 | 3.67 | 2.79 | 2.26 | 5.43 | -0.35 | 0.559 |
| Mariprofundaceae | 4.32 | 4.42 | 4.84 | 2.79 | 3.77 | 5.43 | 0.18 | 0.566 |
| Dysgonomonadaceae | 4.32 | 3.40 | 7.18 | 3.41 | 7.55 | 5.43 | -0.14 | 0.778 |
| Apusomonadidae | 9.71 | 7.48 | 5.17 | 6.98 | 11.32 | 5.43 | -0.09 | 0.847 |
| Tolypothrichaceae | 2.37 | 5.61 | 3.67 | 1.40 | 5.79 | 5.43 | -0.11 | 0.861 |
| Melampsoraceae | 4.96 | 9.00 | 3.84 | 15.05 | 14.84 | 11.37 | -1.21 | 0.019 |
| Salsipaludibacteraceae | 3.02 | 2.04 | 2.17 | 2.48 | 6.04 | 5.17 | -0.92 | 0.175 |
| Tepidiformaceae | 0.65 | 1.53 | 1.50 | 0.31 | 6.04 | 5.17 | -1.65 | 0.279 |
| Pleosporaceae | 5.40 | 2.55 | 6.01 | 2.79 | 1.01 | 5.17 | 0.64 | 0.361 |
| Gracilariaceae | 4.96 | 6.29 | 2.50 | 5.89 | 5.03 | 5.17 | -0.23 | 0.558 |
| Haloferacaceae | 2.16 | 3.40 | 7.68 | 2.33 | 2.52 | 5.17 | 0.40 | 0.611 |
| Spirulinaceae | 2.16 | 3.40 | 2.17 | 0.78 | 3.77 | 5.17 | -0.33 | 0.667 |
| Sarcocystidae | 4.96 | 5.27 | 4.51 | 4.81 | 5.03 | 5.17 | -0.03 | 0.739 |
| Dipodascaceae | 9.28 | 5.27 | 3.51 | 6.51 | 7.04 | 5.17 | -0.05 | 0.910 |
| Bruguierivoracaceae | 4.10 | 7.31 | 7.68 | 9.31 | 5.03 | 5.17 | -0.03 | 0.942 |
| Trichocoleusaceae | 1.08 | 1.70 | 2.84 | 2.02 | 5.54 | 4.91 | -1.15 | 0.158 |
| Bodonidae | 2.81 | 3.23 | 3.51 | 7.14 | 15.10 | 4.91 | -1.51 | 0.198 |
| Thermithiobacillaceae | 1.30 | 1.19 | 1.50 | 0.78 | 3.77 | 4.91 | -1.25 | 0.277 |
| Blastocladiaceae | 4.32 | 4.42 | 3.67 | 4.81 | 3.52 | 4.91 | -0.09 | 0.620 |
| Lyophyllaceae | 4.32 | 5.44 | 8.85 | 8.38 | 8.05 | 4.91 | -0.20 | 0.632 |
| Kytococcaceae | 2.59 | 4.59 | 3.51 | 3.41 | 2.52 | 4.91 | -0.02 | 0.958 |
| Emcibacteraceae | 3.89 | 2.55 | 3.34 | 3.88 | 7.80 | 4.65 | -0.74 | 0.203 |
| Methylacidiphilaceae | 1.73 | 2.04 | 3.51 | 2.33 | 4.28 | 4.65 | -0.63 | 0.221 |
| Melioribacteraceae | 0.65 | 0.17 | 3.17 | 1.71 | 3.02 | 4.65 | -1.23 | 0.228 |
| Candidatus Scalinduaceae | 1.08 | 2.21 | 3.84 | 1.24 | 5.28 | 4.65 | -0.65 | 0.425 |
| Limnochordaceae | 2.37 | 7.81 | 62.59 | 12.56 | 4.28 | 4.65 | 1.76 | 0.468 |
| Roseivirgaceae | 5.18 | 5.61 | 5.01 | 3.10 | 6.04 | 4.65 | 0.20 | 0.516 |
| Magnetococcaceae | 2.37 | 0.85 | 3.34 | 0.62 | 3.52 | 4.65 | -0.42 | 0.630 |
| Marivirgaceae | 2.37 | 5.61 | 2.67 | 2.48 | 5.03 | 4.65 | -0.19 | 0.720 |
| Geoglossaceae | 1.08 | 1.36 | 1.17 | 2.02 | 3.27 | 3.10 | -1.22 | 0.050 |
| Sigmoideomycetaceae | 3.67 | 2.55 | 2.17 | 2.95 | 4.53 | 4.39 | -0.50 | 0.163 |
| Aggregatilineaceae | 2.37 | 1.70 | 4.51 | 3.57 | 4.03 | 4.39 | -0.48 | 0.310 |
| Rubricoccaceae | 2.16 | 2.72 | 3.84 | 2.17 | 6.79 | 4.39 | -0.62 | 0.369 |
| Tepidanaerobacteraceae | 0.43 | 12.91 | 57.92 | 5.74 | 2.01 | 4.39 | 2.55 | 0.376 |
| Amoebophilaceae | 1.94 | 37.38 | 2.67 | 0.31 | 1.76 | 4.39 | 2.70 | 0.418 |
| Amoebophryaceae | 5.83 | 6.12 | 4.01 | 4.19 | 5.54 | 4.39 | 0.18 | 0.485 |
| Zymomonadaceae | 1.30 | 5.95 | 4.17 | 2.17 | 3.27 | 4.39 | 0.21 | 0.750 |
| Acholeplasmataceae | 1.30 | 2.72 | 3.34 | 1.55 | 2.26 | 4.39 | -0.16 | 0.800 |
| Suessiaceae | 5.83 | 4.08 | 3.67 | 4.03 | 5.28 | 4.39 | -0.01 | 0.958 |
| Trichomonascaceae | 1.73 | 2.72 | 2.84 | 2.95 | 3.27 | 4.13 | -0.51 | 0.110 |
| Pyramimonadaceae | 1.51 | 1.36 | 0.33 | 2.17 | 2.01 | 4.13 | -1.38 | 0.113 |
| Plasmodiidae | 7.99 | 5.44 | 5.01 | 3.41 | 5.03 | 4.13 | 0.55 | 0.159 |
| Togniniaceae | 1.73 | 1.02 | 0.50 | 5.27 | 1.01 | 4.13 | -1.68 | 0.196 |
| Onygenaceae | 1.51 | 2.55 | 2.50 | 2.17 | 3.52 | 4.13 | -0.58 | 0.197 |
| Methanoregulaceae | 0.65 | 1.53 | 2.00 | 1.55 | 6.79 | 4.13 | -1.58 | 0.204 |
| Orbaceae | 1.94 | 2.38 | 1.67 | 1.86 | 3.02 | 4.13 | -0.59 | 0.261 |

| | | | | | | | | |
|---|---|---|---|---|---|---|---|---|
| Methylothermaceae | 0.86 | 2.04 | 2.67 | 1.09 | 4.28 | 4.13 | -0.77 | 0.345 |
| Trichosporonaceae | 3.67 | 4.25 | 3.34 | 3.72 | 2.77 | 4.13 | 0.08 | 0.689 |
| Neomegalonemataceae | 2.37 | 1.19 | 2.84 | 1.71 | 1.76 | 4.13 | -0.25 | 0.696 |
| Rubritaleaceae | 1.30 | 1.36 | 2.34 | 2.79 | 5.54 | 3.62 | -1.26 | 0.088 |
| Diaporthaceae | 1.94 | 3.23 | 2.84 | 7.45 | 2.77 | 3.88 | -0.81 | 0.285 |
| Salisaetaceae | 1.30 | 1.02 | 2.67 | 1.40 | 2.52 | 3.88 | -0.64 | 0.354 |
| Helotiaceae | 1.94 | 2.89 | 2.50 | 1.86 | 4.28 | 3.88 | -0.45 | 0.357 |
| Leptosphaeriaceae | 0.43 | 1.70 | 3.67 | 2.17 | 2.77 | 3.88 | -0.60 | 0.415 |
| Pseudocohnilembidae | 4.32 | 1.87 | 2.17 | 2.33 | 5.03 | 3.88 | -0.43 | 0.432 |
| Kangiellaceae | 3.89 | 2.04 | 1.67 | 1.40 | 5.28 | 3.88 | -0.48 | 0.507 |
| Entamoebidae | 2.37 | 3.23 | 1.50 | 2.48 | 2.26 | 3.88 | -0.28 | 0.515 |
| Pleurotaceae | 5.18 | 3.06 | 2.17 | 2.79 | 5.28 | 3.88 | -0.20 | 0.678 |
| Thermosediminibacteraceae | 1.51 | 3.91 | 7.18 | 3.57 | 3.02 | 3.88 | 0.27 | 0.709 |
| Gonapodyaceae | 7.12 | 5.78 | 4.84 | 5.89 | 6.79 | 3.88 | 0.10 | 0.738 |
| Phaeodactylaceae | 2.59 | 3.57 | 2.84 | 3.41 | 2.26 | 3.88 | -0.09 | 0.761 |
| Phanerochaetaceae | 2.59 | 3.40 | 3.84 | 2.95 | 2.52 | 3.88 | 0.07 | 0.779 |
| Lichtheimiaceae | 4.10 | 5.61 | 3.67 | 4.65 | 5.28 | 3.88 | -0.05 | 0.851 |
| Hydrogenophilaceae | 2.16 | 2.21 | 6.01 | 2.02 | 3.77 | 3.88 | 0.10 | 0.878 |
| Sulfurovaceae | 1.94 | 3.06 | 3.00 | 1.55 | 2.26 | 3.88 | 0.06 | 0.901 |
| Gelatoporiaceae | 3.67 | 6.80 | 3.17 | 11.48 | 12.33 | 10.08 | -1.31 | 0.012 |
| Pyriculariaceae | 1.08 | 1.36 | 0.50 | 3.88 | 1.51 | 3.62 | -1.62 | 0.102 |
| Wallemiaceae | 1.94 | 1.87 | 1.34 | 2.33 | 4.78 | 3.62 | -1.06 | 0.111 |
| Algiphilaceae | 1.73 | 0.85 | 1.00 | 1.09 | 3.27 | 3.62 | -1.16 | 0.198 |
| Sporidiobolaceae | 4.32 | 4.25 | 2.84 | 5.89 | 5.54 | 3.62 | -0.40 | 0.238 |
| Ceraceosoraceae | 1.30 | 0.51 | 0.33 | 0.47 | 1.51 | 3.62 | -1.39 | 0.341 |
| Cafeteriaceae | 5.18 | 4.25 | 3.34 | 2.79 | 5.03 | 3.62 | 0.16 | 0.629 |
| Eimeriidae | 1.51 | 2.89 | 3.34 | 2.17 | 2.77 | 3.62 | -0.15 | 0.715 |
| Symbiobacteriaceae | 3.89 | 12.91 | 53.08 | 42.97 | 3.02 | 3.62 | 0.49 | 0.754 |
| Thermostichaceae | 1.94 | 2.55 | 2.50 | 2.17 | 1.76 | 3.62 | -0.11 | 0.780 |
| Hoehnelomycetaceae | 5.40 | 0.85 | 1.50 | 2.17 | 3.27 | 3.62 | -0.23 | 0.792 |
| Desertifilaceae | 1.30 | 4.08 | 5.51 | 3.57 | 4.78 | 3.62 | -0.14 | 0.803 |
| Podosporaceae | 5.83 | 1.53 | 1.00 | 3.10 | 0.50 | 3.62 | 0.21 | 0.846 |
| Muribaculaceae | 1.94 | 2.89 | 2.17 | 3.10 | 2.52 | 3.36 | -0.36 | 0.158 |
| Thermomicrobiaceae | 0.65 | 0.68 | 1.17 | 0.78 | 2.52 | 3.36 | -1.41 | 0.206 |
| Chrysiogenaceae | 0.86 | 0.85 | 1.17 | 0.78 | 2.52 | 3.36 | -1.21 | 0.239 |
| Oculatellaceae | 1.30 | 1.36 | 2.50 | 1.71 | 2.52 | 3.36 | -0.56 | 0.264 |
| Candidatus Methanoperedenaceae | 1.73 | 0.68 | 1.34 | 1.09 | 2.26 | 3.36 | -0.84 | 0.270 |
| Filobasidiaceae | 2.81 | 1.36 | 1.34 | 1.71 | 3.02 | 3.36 | -0.56 | 0.286 |
| Lentisphaeraceae | 0.22 | 0.85 | 1.50 | 1.40 | 0.75 | 3.36 | -1.10 | 0.344 |
| Candidatus Thermochlorobacteriaceae | 0.86 | 1.19 | 2.84 | 1.24 | 3.02 | 3.36 | -0.64 | 0.368 |
| Mycoplasmataceae | 1.30 | 3.06 | 2.50 | 2.17 | 3.02 | 3.36 | -0.32 | 0.427 |
| Marasmiaceae | 3.67 | 4.25 | 2.00 | 3.72 | 5.03 | 3.36 | -0.29 | 0.438 |
| Chaetosphaeriaceae | 4.32 | 1.87 | 1.34 | 4.81 | 2.26 | 3.36 | -0.47 | 0.458 |
| Casimicrobiaceae | 1.08 | 0.68 | 1.34 | 0.78 | 1.01 | 3.36 | -0.73 | 0.498 |
| Phototrophicaceae | 1.51 | 1.87 | 2.00 | 0.78 | 3.27 | 3.36 | -0.46 | 0.511 |
| Vallitaleaceae | 1.51 | 3.57 | 4.51 | 3.10 | 2.01 | 3.36 | 0.18 | 0.731 |
| Granulosicoccaceae | 1.94 | 1.53 | 4.17 | 1.55 | 3.77 | 3.36 | -0.18 | 0.762 |
| Elaphomycetaceae | 1.51 | 3.57 | 4.01 | 2.33 | 4.28 | 3.36 | -0.13 | 0.776 |
| Pluteaceae | 4.32 | 3.40 | 3.00 | 3.72 | 3.27 | 3.36 | 0.05 | 0.790 |
| Didymosphaeriaceae | 2.81 | 1.36 | 1.50 | 0.78 | 1.26 | 3.36 | 0.07 | 0.926 |
| Hypocreaceae | 5.18 | 7.48 | 7.18 | 15.82 | 13.08 | 21.18 | -1.34 | 0.042 |
| Andaluciidae | 3.89 | 2.89 | 3.67 | 1.71 | 2.52 | 3.10 | 0.51 | 0.114 |
| Fibrobacteraceae | 0.65 | 0.68 | 1.50 | 1.24 | 4.03 | 3.10 | -1.56 | 0.142 |
| Candidatus Eremiobacteraceae | 0.86 | 1.02 | 0.67 | 0.47 | 2.26 | 3.10 | -1.19 | 0.294 |
| Ferrovaceae | 2.59 | 1.70 | 2.17 | 1.86 | 4.53 | 3.10 | -0.56 | 0.320 |
| Parmeliaceae | 0.65 | 0.68 | 1.17 | 0.78 | 1.26 | 3.10 | -1.04 | 0.340 |
| Sphaerochaetaceae | 0.86 | 1.70 | 1.67 | 1.09 | 2.01 | 3.10 | -0.55 | 0.387 |
| Peniophoraceae | 0.43 | 1.02 | 1.34 | 1.55 | 0.50 | 3.10 | -0.89 | 0.410 |
| Gregarinidae | 2.59 | 3.91 | 1.67 | 3.26 | 3.77 | 3.10 | -0.31 | 0.423 |
| Catalimonadaceae | 1.94 | 5.61 | 4.51 | 3.10 | 3.27 | 3.10 | 0.35 | 0.511 |
| Caldicoprobacteraceae | 1.73 | 12.40 | 15.52 | 78.95 | 2.77 | 3.10 | -1.52 | 0.545 |
| Christensenellaceae | 0.86 | 3.06 | 4.51 | 2.02 | 1.26 | 3.10 | 0.40 | 0.605 |
| Cesiribacteraceae | 4.32 | 4.25 | 7.34 | 4.96 | 5.79 | 3.10 | 0.20 | 0.626 |
| Kosmotogaceae | 0.22 | 1.19 | 3.67 | 1.40 | 2.26 | 3.10 | -0.41 | 0.658 |
| Microsporaceae | 1.73 | 1.19 | 2.17 | 1.71 | 1.01 | 3.10 | -0.19 | 0.746 |
| Syntrophotaleaceae | 1.08 | 1.87 | 2.67 | 0.31 | 3.27 | 3.10 | -0.25 | 0.762 |
| Russulaceae | 1.08 | 3.23 | 2.50 | 3.41 | 1.01 | 3.10 | -0.14 | 0.823 |
| Fervidobacteriaceae | 1.08 | 1.87 | 3.34 | 0.93 | 2.01 | 3.10 | 0.06 | 0.933 |
| Prasiolaceae | 2.59 | 0.00 | 1.34 | 0.00 | 0.50 | 3.10 | 0.12 | 0.934 |
| Desulfonatronaceae | 1.73 | 1.19 | 2.00 | 3.72 | 3.77 | 5.17 | -1.36 | 0.017 |
| Zooshikellaceae | 1.30 | 1.02 | 1.50 | 1.55 | 2.26 | 2.84 | -0.80 | 0.113 |
| Catenariaceae | 4.53 | 2.72 | 3.00 | 1.71 | 2.26 | 2.84 | 0.59 | 0.170 |
| Microbotryaceae | 2.59 | 2.55 | 1.84 | 2.48 | 3.27 | 2.84 | -0.30 | 0.182 |
| Candidatus Babeliaceae | 9.71 | 2.04 | 9.85 | 4.34 | 3.77 | 2.84 | 0.98 | 0.302 |
| Microdochiaceae | 1.08 | 2.21 | 0.33 | 2.02 | 1.26 | 2.84 | -0.76 | 0.309 |
| Budviciaceae | 1.94 | 0.68 | 1.67 | 1.09 | 2.77 | 2.84 | -0.64 | 0.319 |
| Rhodothalassiaceae | 2.16 | 0.85 | 1.17 | 1.24 | 2.01 | 2.84 | -0.55 | 0.353 |
| Hydnaceae | 1.30 | 1.02 | 0.67 | 1.09 | 1.01 | 2.84 | -0.73 | 0.393 |
| Thermodesulfovibrionaceae | 1.94 | 1.02 | 0.67 | 0.31 | 3.77 | 2.84 | -0.93 | 0.405 |
| Thiovulaceae | 1.94 | 2.38 | 3.51 | 2.48 | 4.78 | 2.84 | -0.37 | 0.431 |
| Mycenaceae | 3.89 | 4.25 | 3.67 | 4.65 | 3.52 | 2.84 | 0.10 | 0.676 |
| Thermotogaceae | 1.51 | 3.23 | 8.01 | 2.02 | 5.03 | 2.84 | 0.37 | 0.688 |
| Sedimentisphaeraceae | 1.94 | 3.23 | 3.84 | 2.17 | 3.27 | 2.84 | 0.12 | 0.731 |
| Crenotrichaceae | 2.59 | 0.85 | 1.50 | 0.78 | 2.01 | 2.84 | -0.19 | 0.785 |
| Tilletiaceae | 3.45 | 5.44 | 2.84 | 4.96 | 3.77 | 2.84 | 0.02 | 0.963 |
| Brachyspiraceae | 5.61 | 1.02 | 1.00 | 0.78 | 3.77 | 2.84 | 0.05 | 0.966 |
| Olpidiaceae | 1.73 | 2.04 | 2.00 | 2.33 | 2.26 | 2.58 | -0.31 | 0.028 |
| Dermocarpellaceae | 1.08 | 1.19 | 1.67 | 1.40 | 1.76 | 2.58 | -0.54 | 0.226 |
| Acaryochloridaceae | 1.94 | 1.53 | 2.34 | 2.02 | 3.77 | 2.58 | -0.53 | 0.236 |
| Thermohalobacteraceae | 0.86 | 7.65 | 8.18 | 2.95 | 1.51 | 2.58 | 1.25 | 0.304 |
| Chaetocerotaceae | 3.89 | 3.57 | 3.00 | 2.33 | 3.77 | 2.58 | 0.27 | 0.329 |
| Hericiaceae | 2.16 | 3.06 | 1.50 | 3.26 | 2.52 | 2.58 | -0.31 | 0.362 |
| Tricholomataceae | 1.08 | 2.72 | 0.67 | 2.33 | 1.76 | 2.58 | -0.58 | 0.365 |
| Sulfolobaceae | 1.08 | 1.02 | 3.00 | 1.71 | 2.52 | 2.58 | -0.42 | 0.488 |
| Caulochytriaceae | 2.37 | 2.89 | 2.00 | 2.33 | 2.77 | 2.58 | -0.08 | 0.665 |
| Gottschalkiaceae | 0.43 | 2.72 | 2.67 | 0.93 | 1.01 | 2.58 | 0.36 | 0.667 |
| Naviculaceae | 3.45 | 3.06 | 5.01 | 2.64 | 5.03 | 2.58 | 0.17 | 0.697 |
| Woeseiaceae | 1.30 | 2.55 | 2.84 | 1.71 | 3.02 | 2.58 | -0.13 | 0.750 |
| Leptotrichiaceae | 2.16 | 3.23 | 4.01 | 2.79 | 3.52 | 2.58 | 0.08 | 0.803 |
| Pezizaceae | 2.81 | 1.87 | 0.50 | 1.09 | 1.76 | 2.58 | -0.07 | 0.921 |

| | | | | | | | | |
|---|---|---|---|---|---|---|---|---|
| Cardiobacteriaceae | 1.51 | 2.38 | 2.34 | 0.78 | 2.77 | 2.58 | 0.02 | 0.965 |
| Kickxellaceae | 6.26 | 4.42 | 3.84 | 6.05 | 6.04 | 2.58 | -0.02 | 0.972 |
| Stachybotryaceae | 2.81 | 4.42 | 1.50 | 9.77 | 4.53 | 9.30 | -1.44 | 0.079 |
| Oligoflexaceae | 4.75 | 20.22 | 12.18 | 25.75 | 44.03 | 32.55 | -1.46 | 0.037 |
| Schleiferiaceae | 0.65 | 1.87 | 1.50 | 2.48 | 4.78 | 2.33 | -1.25 | 0.130 |
| Ceratocystidaceae | 2.59 | 0.85 | 0.83 | 2.48 | 3.27 | 2.33 | -0.92 | 0.149 |
| Cryphonectriaceae | 0.86 | 1.36 | 0.83 | 2.79 | 1.01 | 2.33 | -1.00 | 0.189 |
| Gloniaceae | 0.86 | 0.85 | 1.00 | 0.93 | 1.76 | 2.33 | -0.89 | 0.197 |
| Thermococcaceae | 0.86 | 1.19 | 1.34 | 1.09 | 2.01 | 2.33 | -0.68 | 0.202 |
| Strophariaceae | 1.94 | 1.53 | 1.34 | 1.86 | 4.28 | 2.33 | -0.82 | 0.238 |
| Schizoporaceae | 1.30 | 2.55 | 3.34 | 0.93 | 1.01 | 2.33 | 0.75 | 0.267 |
| Tricladiaceae | 1.08 | 0.51 | 1.17 | 0.62 | 2.52 | 2.33 | -0.99 | 0.270 |
| Cryptosporidiidae | 4.10 | 3.06 | 2.50 | 4.96 | 4.28 | 2.33 | -0.26 | 0.536 |
| Ancylistaceae | 2.59 | 3.74 | 1.67 | 2.02 | 2.77 | 2.33 | 0.17 | 0.679 |
| Amanitaceae | 3.45 | 2.72 | 1.67 | 3.26 | 3.02 | 2.33 | -0.13 | 0.695 |
| Microscillaceae | 1.08 | 2.72 | 2.84 | 1.71 | 3.27 | 2.33 | -0.14 | 0.776 |
| Zhaonellaceae | 0.86 | 2.04 | 4.01 | 0.62 | 3.27 | 2.33 | 0.15 | 0.857 |
| Malawimonadidae | 0.65 | 0.00 | 0.00 | 0.47 | 4.53 | 2.07 | -3.45 | 0.209 |
| Pleuroascaceae | 0.43 | 0.68 | 1.00 | 0.31 | 3.02 | 2.07 | -1.35 | 0.300 |
| Flexibacteraceae | 1.73 | 4.76 | 4.17 | 1.71 | 3.27 | 2.07 | 0.60 | 0.332 |
| Punctulariaceae | 0.22 | 2.72 | 0.33 | 2.17 | 1.76 | 2.07 | -0.88 | 0.380 |
| Spiroplasmataceae | 1.51 | 1.36 | 1.17 | 1.24 | 1.51 | 2.07 | -0.25 | 0.405 |
| Claroideoglomeraceae | 3.67 | 4.08 | 3.51 | 4.34 | 2.77 | 2.07 | 0.29 | 0.414 |
| Thermonemataceae | 0.86 | 1.53 | 2.67 | 0.62 | 1.01 | 2.07 | 0.46 | 0.541 |
| Thermoanaerobacterales Family IV. Incertae Sedis | 0.65 | 2.38 | 3.67 | 1.86 | 1.01 | 2.07 | 0.44 | 0.581 |
| Pyronemataceae | 2.16 | 1.87 | 1.67 | 1.86 | 1.26 | 2.07 | 0.14 | 0.585 |
| Candidatus Uabimicrobiaceae | 1.94 | 1.19 | 1.84 | 0.78 | 3.02 | 2.07 | -0.24 | 0.701 |
| Desulfurobacteriaceae | 1.73 | 1.53 | 2.34 | 1.55 | 2.26 | 2.07 | -0.07 | 0.780 |
| Phycomycetaceae | 3.45 | 2.55 | 1.00 | 2.95 | 1.76 | 2.07 | 0.05 | 0.930 |
| Candidatus Midichloriaceae | 2.59 | 0.68 | 0.67 | 0.47 | 1.26 | 2.07 | 0.06 | 0.954 |
| Massarinaceae | 2.59 | 1.02 | 1.00 | 0.62 | 2.01 | 2.07 | -0.03 | 0.969 |
| Tuberaceae | 0.65 | 0.68 | 0.67 | 1.71 | 2.26 | 1.55 | -1.47 | 0.032 |
| Thyridiaceae | 2.37 | 1.53 | 0.33 | 5.27 | 2.26 | 4.39 | -1.49 | 0.085 |
| Hafniaceae | 6.48 | 3.23 | 5.67 | 2.95 | 3.02 | 1.81 | 0.98 | 0.107 |
| Eubacteriales Family XII. Incertae Sedis | 2.37 | 3.40 | 3.51 | 2.48 | 2.52 | 1.81 | 0.45 | 0.139 |
| Cladosporiaceae | 0.86 | 2.38 | 1.34 | 4.65 | 2.26 | 1.81 | -0.93 | 0.258 |
| Pontiellaceae | 0.43 | 1.70 | 1.84 | 1.40 | 3.02 | 1.81 | -0.65 | 0.319 |
| Schizosaccharomycetaceae | 5.18 | 4.93 | 2.67 | 4.03 | 3.52 | 1.81 | 0.45 | 0.338 |
| Babesiidae | 2.37 | 2.72 | 1.17 | 2.95 | 4.03 | 1.81 | -0.49 | 0.355 |
| Dissoconiaceae | 0.43 | 1.36 | 1.50 | 1.71 | 1.01 | 1.81 | -0.46 | 0.389 |
| Proteinivoraceae | 0.43 | 3.23 | 5.51 | 0.93 | 2.01 | 1.81 | 0.95 | 0.423 |
| Aquificaceae | 1.08 | 1.70 | 1.34 | 0.93 | 4.28 | 1.81 | -0.77 | 0.437 |
| Thermoflexibacteraceae | 0.65 | 3.57 | 3.34 | 1.55 | 1.51 | 1.81 | 0.63 | 0.441 |
| Turicibacteraceae | 0.65 | 4.42 | 2.17 | 1.24 | 1.26 | 1.81 | 0.75 | 0.468 |
| Chroococcidiopsidaceae | 0.22 | 1.87 | 2.34 | 1.24 | 3.02 | 1.81 | -0.46 | 0.546 |
| Dimargaritaceae | 2.59 | 3.23 | 1.67 | 2.17 | 2.52 | 1.81 | 0.20 | 0.557 |
| Archaeoglobaceae | 0.86 | 0.34 | 2.34 | 0.31 | 3.52 | 1.81 | -0.67 | 0.566 |
| Bernardetiaceae | 0.65 | 1.70 | 1.84 | 0.78 | 1.01 | 1.81 | 0.22 | 0.708 |
| Hydrogenothermaceae | 0.86 | 3.74 | 3.51 | 2.17 | 3.02 | 1.81 | 0.21 | 0.737 |
| Polymastigidae | 2.16 | 2.04 | 1.17 | 1.86 | 2.01 | 1.81 | -0.08 | 0.769 |
| Petrotogaceae | 1.08 | 3.57 | 3.84 | 2.64 | 4.03 | 1.81 | 0.00 | 0.997 |
| Gemmatimonadaceae | 16.41 | 47.74 | 72.44 | 117.26 | 168.07 | 208.74 | -1.85 | 0.026 |
| Succinivibrionaceae | 1.73 | 1.70 | 2.34 | 1.40 | 1.26 | 1.55 | 0.45 | 0.116 |
| Elsinoaceae | 0.86 | 1.19 | 1.34 | 2.95 | 1.76 | 1.55 | -0.89 | 0.149 |
| Phaeosphaeriaceae | 1.94 | 1.70 | 2.34 | 1.86 | 1.26 | 1.55 | 0.36 | 0.162 |
| Choanephoraceae | 0.86 | 1.53 | 0.83 | 1.40 | 3.27 | 1.55 | -0.95 | 0.234 |
| Euglenaceae | 0.65 | 0.68 | 1.67 | 2.33 | 1.01 | 1.55 | -0.70 | 0.285 |
| Prasinococcaceae | 1.51 | 1.87 | 0.67 | 1.40 | 2.52 | 1.55 | -0.43 | 0.399 |
| Immundisolibacteraceae | 1.73 | 0.68 | 1.34 | 0.78 | 3.77 | 1.55 | -0.70 | 0.480 |
| Venturiaceae | 1.94 | 1.19 | 0.83 | 1.09 | 1.76 | 1.55 | -0.15 | 0.731 |
| Anaerohalosphaeraceae | 0.65 | 2.38 | 2.67 | 1.55 | 2.01 | 1.55 | 0.16 | 0.791 |
| Dermateaceae | 0.86 | 2.21 | 1.34 | 1.24 | 1.26 | 1.55 | 0.12 | 0.794 |
| Mangrovivirgaceae | 1.08 | 1.87 | 1.84 | 0.93 | 1.01 | 1.29 | 0.57 | 0.172 |
| Syncephalastraceae | 1.08 | 1.02 | 1.50 | 2.79 | 3.77 | 1.29 | -1.13 | 0.184 |
| Chamaesiphonaceae | 1.73 | 2.04 | 4.84 | 0.93 | 2.77 | 1.29 | 0.79 | 0.364 |
| Entomophthoraceae | 2.16 | 1.53 | 0.83 | 3.41 | 2.01 | 1.29 | -0.57 | 0.384 |
| Koleobacteraceae | 0.43 | 2.38 | 3.17 | 0.93 | 1.26 | 1.29 | 0.78 | 0.414 |
| Natranaerobiaceae | 0.65 | 2.04 | 1.84 | 0.93 | 1.01 | 1.29 | 0.49 | 0.427 |
| Arenicellaceae | 1.08 | 1.36 | 2.17 | 1.55 | 1.01 | 1.29 | 0.26 | 0.538 |
| Paludibacteraceae | 1.73 | 1.87 | 2.00 | 1.09 | 5.03 | 1.29 | -0.40 | 0.684 |
| Aliterellaceae | 0.86 | 1.19 | 2.50 | 1.71 | 1.01 | 1.29 | 0.19 | 0.759 |
| Physciaceae | 0.43 | 1.19 | 2.00 | 1.55 | 1.26 | 1.29 | -0.18 | 0.762 |
| Ambisporaceae | 3.89 | 2.89 | 2.50 | 4.03 | 4.78 | 1.29 | -0.12 | 0.826 |
| Neolectaceae | 1.73 | 1.36 | 1.34 | 1.09 | 1.26 | 1.03 | 0.39 | 0.092 |
| Defluviitaleaceae | 1.73 | 1.53 | 5.17 | 1.24 | 0.75 | 1.03 | 1.48 | 0.267 |
| Xylanivirgaceae | 0.65 | 5.10 | 3.67 | 1.71 | 1.26 | 1.03 | 1.24 | 0.301 |
| Saccharomycodaceae | 4.75 | 2.72 | 0.50 | 2.02 | 0.75 | 1.03 | 1.07 | 0.377 |
| Magnaporthaceae | 1.51 | 1.19 | 0.33 | 3.72 | 0.75 | 1.03 | -0.86 | 0.483 |
| Desulfosalsimonadaceae | 1.73 | 2.04 | 0.67 | 2.17 | 0.50 | 1.03 | 0.26 | 0.727 |
| Palmophyllaceae | 2.16 | 1.87 | 1.00 | 2.33 | 1.76 | 1.03 | -0.03 | 0.955 |
| Bacillales Family X. Incertae Sedis | 1.51 | 2.21 | 3.34 | 2.17 | 0.75 | 0.78 | 0.93 | 0.191 |
| Trichomonadidae | 1.51 | 2.55 | 1.34 | 1.24 | 1.26 | 0.78 | 0.72 | 0.194 |
| Omphalotaceae | 2.59 | 2.38 | 1.50 | 1.09 | 2.77 | 0.78 | 0.48 | 0.445 |
| Mixiaceae | 1.94 | 1.53 | 0.83 | 2.02 | 2.01 | 0.78 | -0.16 | 0.768 |
| Boletaceae | 1.51 | 1.53 | 1.34 | 1.24 | 2.77 | 0.52 | -0.05 | 0.947 |
| Tritrichomonadidae | 2.16 | 1.53 | 1.00 | 0.78 | 1.76 | 0.26 | 0.75 | 0.322 |

RPM, reads per million.
