## Supplementary material for "Horizontally acquired *IbACS* gene modulates rhizosphere microbiota and contributes to sweet potato growth": Table S2

**Table S2.** Complete list of rhizosphere microbial genera detected in the IbACS-KO8 and control sweet potato lines, with normalized read counts, fold-change values, and statistical significance.

| Family | EV-1 (RPM) | EV-2 (RPM) | EV-3 (RPM) | KO8-1 (RPM) | KO8-2 (RPM) | KO8-3 (RPM) | log2FC (EV/KO8) | *p* value |
|---|---|---|---|---|---|---|---|---|
| Unclassified | 489400.86 | 598043.02 | 464774.83 | 620334.90 | 702897.31 | 651802.32 | -0.35 | 0.054 |
| cannot be assigned to a (non-viral) family | 157504.13 | 79579.76 | 123058.60 | 98529.26 | 78148.48 | 91367.50 | 0.43 | 0.305 |
| *Streptomyces* | 27994.95 | 48668.39 | 26791.52 | 32210.17 | 35487.99 | 34449.44 | 0.02 | 0.955 |
| *Rhizobium* | 22015.45 | 29854.11 | 45873.80 | 19072.01 | 10730.51 | 14909.79 | 1.13 | 0.116 |
| *Mesorhizobium* | 8213.98 | 3373.20 | 7159.28 | 4522.71 | 9863.80 | 14516.64 | -0.62 | 0.374 |
| *Bradyrhizobium* | 2319.25 | 5150.96 | 5663.42 | 2782.08 | 8297.48 | 9355.73 | -0.64 | 0.366 |
| *Saccharopolyspora* | 7425.43 | 5579.12 | 4825.54 | 6404.49 | 4497.15 | 7539.85 | -0.05 | 0.874 |
| *Sphingomonas* | 4878.74 | 9521.95 | 12065.78 | 5592.98 | 11139.94 | 6936.98 | 0.16 | 0.746 |
| *Paenibacillus* | 8645.87 | 27380.64 | 9316.30 | 11854.30 | 4001.63 | 6461.61 | 1.02 | 0.339 |
| *Shinella* | 636.56 | 1384.78 | 2178.86 | 1182.28 | 4604.25 | 6333.60 | -1.53 | 0.217 |
| *Mycobacterium* | 849.48 | 1293.55 | 6017.52 | 926.79 | 4130.76 | 6023.97 | -0.44 | 0.685 |
| *Nocardioides* | 7031.69 | 1958.69 | 6786.87 | 2374.07 | 3470.41 | 5454.67 | 0.48 | 0.483 |
| *Bosea* | 2062.55 | 4965.58 | 10320.31 | 6154.91 | 2125.90 | 4488.05 | 0.44 | 0.611 |
| *Pseudonocardia* | 2334.65 | 3072.84 | 2518.01 | 5108.01 | 3298.74 | 4259.08 | -0.68 | 0.079 |
| *Phyllobacterium* | 264.08 | 327.19 | 405.84 | 348.03 | 1866.11 | 4119.62 | -2.67 | 0.246 |
| *Stenotrophomonas* | 117399.50 | 830.53 | 22398.86 | 16168.57 | 2396.82 | 3752.23 | 2.66 | 0.386 |
| *Leifsonia* | 17110.92 | 13660.93 | 41011.13 | 25942.77 | 2341.37 | 3264.89 | 1.18 | 0.311 |
| *Devosia* | 2915.92 | 6897.10 | 8333.28 | 1801.23 | 3051.10 | 3118.66 | 1.19 | 0.164 |
| *Pseudolabrys* | 7126.66 | 4039.20 | 9166.97 | 5212.08 | 3164.28 | 3005.48 | 0.84 | 0.172 |
| *Paraburkholderia* | 1124.39 | 454.99 | 300.68 | 467.99 | 4547.03 | 2831.67 | -2.06 | 0.231 |
| *Microbacterium* | 871.16 | 2246.24 | 6023.40 | 1370.01 | 1541.25 | 2728.11 | 0.70 | 0.532 |
| *Nitrobacter* | 419.75 | 280.89 | 322.35 | 144.42 | 1800.02 | 2631.58 | -2.16 | 0.246 |
| *Hyphomicrobium* | 246.73 | 154.80 | 378.12 | 234.31 | 2024.87 | 2620.65 | -2.64 | 0.196 |
| *Candidatus* Nitrosocosmicus | 36.86 | 7.01 | 11.09 | 4.05 | 1897.76 | 2614.15 | -6.36 | 0.196 |
| *Rhodococcus* | 825.19 | 6265.79 | 7535.89 | 514.42 | 1384.77 | 2482.49 | 1.74 | 0.234 |
| *Gemmata* | 831.70 | 329.24 | 1143.10 | 293.20 | 2151.72 | 2333.66 | -1.05 | 0.335 |
| *Lysobacter* | 669.74 | 131.73 | 1038.78 | 92.69 | 1988.15 | 2156.73 | -1.20 | 0.354 |
| *Escherichia* | 1412.11 | 2701.06 | 1082.46 | 2875.09 | 2340.61 | 2154.13 | -0.50 | 0.280 |
| *Amycolatopsis* | 1795.43 | 3299.73 | 2554.13 | 2577.06 | 1702.29 | 2140.08 | 0.25 | 0.470 |
| *Sphingobium* | 365.11 | 784.06 | 2004.83 | 1059.99 | 1547.07 | 2036.78 | -0.56 | 0.442 |
| *Pseudomonas* | 1181.20 | 1506.95 | 1869.27 | 1442.92 | 1166.26 | 2027.68 | -0.02 | 0.940 |
| *Clostridium* | 1385.00 | 3043.45 | 1267.57 | 3061.72 | 2323.40 | 1989.95 | -0.37 | 0.454 |
| *Nocardia* | 441.43 | 780.13 | 577.51 | 569.10 | 1929.92 | 1839.04 | -1.27 | 0.189 |
| *Penicillium* | 523.60 | 1521.30 | 3146.08 | 862.14 | 1030.29 | 1828.11 | 0.48 | 0.598 |
| *Ensifer* | 3896.35 | 7310.91 | 9818.89 | 2986.94 | 340.05 | 1674.86 | 2.07 | 0.072 |
| *Sphingopyxis* | 236.11 | 4911.07 | 12233.93 | 6639.89 | 1322.23 | 1666.27 | 0.85 | 0.556 |
| *Afipia* | 1378.28 | 2256.83 | 4944.30 | 2710.89 | 1289.31 | 1538.00 | 0.63 | 0.454 |
| *Cellulomonas* | 135.73 | 123.53 | 359.64 | 134.13 | 971.54 | 1461.24 | -2.05 | 0.233 |
| *Simplicispira* | 5.85 | 45.45 | 349.73 | 28.51 | 1136.38 | 1403.74 | -2.68 | 0.224 |
| *Novosphingobium* | 604.48 | 3222.00 | 2066.65 | 1533.74 | 1226.26 | 1402.96 | 0.50 | 0.526 |
| *Asticcacaulis* | 1612.01 | 3033.71 | 2285.36 | 975.55 | 2883.48 | 1318.13 | 0.42 | 0.464 |
| *Actinomadura* | 859.01 | 2758.63 | 1321.33 | 1471.58 | 1009.02 | 1270.00 | 0.40 | 0.563 |
| *Acinetobacter* | 744.97 | 1295.26 | 537.20 | 1638.59 | 1286.27 | 1095.67 | -0.64 | 0.166 |
| *Brucella* | 868.99 | 190.85 | 373.25 | 1706.20 | 723.91 | 1062.36 | -1.28 | 0.132 |
| *Frigoriglobus* | 157.41 | 61.68 | 233.66 | 62.63 | 1039.91 | 1059.50 | -2.25 | 0.224 |
| *Mycolicibacterium* | 622.69 | 734.85 | 2401.94 | 598.85 | 570.97 | 1022.56 | 0.78 | 0.462 |
| *Dyadobacter* | 442.73 | 178.37 | 1192.99 | 562.09 | 882.16 | 960.11 | -0.41 | 0.596 |
| *Caulifigura* | 71.33 | 122.50 | 310.76 | 196.45 | 559.83 | 818.30 | -1.64 | 0.176 |
| *Marmoricola* | 1139.57 | 289.77 | 1521.22 | 244.43 | 405.63 | 801.39 | 1.02 | 0.305 |
| *Conexibacter* | 149.17 | 531.02 | 220.39 | 114.19 | 658.58 | 743.89 | -0.75 | 0.432 |
| *Frankia* | 387.45 | 459.94 | 438.09 | 515.66 | 590.72 | 714.49 | -0.50 | 0.076 |
| *Rhodoplanes* | 501.71 | 180.59 | 433.05 | 158.59 | 661.37 | 690.55 | -0.44 | 0.553 |
| *Rhodanobacter* | 2903.78 | 18644.45 | 14208.53 | 12977.38 | 1734.95 | 676.76 | 1.21 | 0.332 |
| *Singulisphaera* | 212.04 | 235.27 | 1096.57 | 329.96 | 462.10 | 663.75 | 0.09 | 0.931 |
| *Paludisphaera* | 150.90 | 1339.85 | 1602.52 | 644.81 | 461.59 | 644.24 | 0.82 | 0.422 |
| *Bacillus* | 522.95 | 3328.44 | 4725.76 | 2037.57 | 586.17 | 610.93 | 1.41 | 0.285 |
| *Urbifossiella* | 89.11 | 62.87 | 262.38 | 129.93 | 579.84 | 609.63 | -1.67 | 0.182 |
| *Fimbriiglobus* | 79.14 | 80.99 | 161.76 | 84.75 | 492.48 | 566.18 | -1.83 | 0.206 |
| *Luteibacter* | 22.77 | 73.47 | 73.24 | 63.41 | 528.44 | 559.41 | -2.76 | 0.177 |
| *Nonomuraea* | 408.91 | 913.22 | 573.31 | 583.27 | 443.11 | 557.33 | 0.26 | 0.561 |
| *Rhodopseudomonas* | 194.26 | 338.64 | 411.38 | 227.45 | 475.77 | 549.01 | -0.41 | 0.435 |
| *Actinoplanes* | 237.41 | 294.38 | 272.80 | 262.66 | 270.93 | 528.71 | -0.40 | 0.430 |
| *Flexivirga* | 71.55 | 56.55 | 109.02 | 34.12 | 191.42 | 518.56 | -1.65 | 0.358 |
| *Planctomyces* | 272.97 | 2795.71 | 5107.24 | 4679.28 | 485.14 | 513.10 | 0.52 | 0.696 |
| *Agrobacterium* | 1310.64 | 1460.47 | 2303.16 | 2005.94 | 356.26 | 507.63 | 0.82 | 0.310 |
| *Microlunatus* | 267.76 | 651.47 | 762.46 | 581.56 | 216.74 | 505.55 | 0.37 | 0.539 |
| *Rhizophagus* | 264.08 | 403.39 | 169.32 | 731.12 | 566.42 | 489.94 | -1.09 | 0.033 |
| *Cohnella* | 550.71 | 2514.48 | 1569.09 | 2549.64 | 302.83 | 479.27 | 0.47 | 0.663 |
| *Micromonospora* | 268.42 | 969.43 | 3244.86 | 1878.35 | 365.12 | 478.23 | 0.72 | 0.605 |
| *Aspergillus* | 184.29 | 398.95 | 710.38 | 243.34 | 330.68 | 447.27 | 0.34 | 0.623 |
| *Viruses* | 209.01 | 392.80 | 178.90 | 414.40 | 434.24 | 416.05 | -0.70 | 0.136 |
| *Kribbella* | 285.33 | 1235.97 | 1637.97 | 587.79 | 273.21 | 407.20 | 1.32 | 0.253 |
| *Delftia* | 21526.32 | 855.64 | 2904.87 | 2953.92 | 727.71 | 406.42 | 2.63 | 0.395 |
| *Burkholderia* | 371.62 | 259.87 | 253.65 | 234.15 | 498.30 | 395.75 | -0.35 | 0.417 |
| *Legionella* | 713.75 | 99.27 | 161.76 | 51.41 | 285.36 | 395.23 | 0.42 | 0.736 |
| *Klebsiella* | 298.55 | 335.56 | 207.79 | 453.03 | 369.68 | 392.11 | -0.53 | 0.062 |
| *Massilia* | 1190.74 | 2689.44 | 2173.65 | 570.66 | 358.54 | 388.21 | 2.20 | 0.066 |
| *Bdellovibrio* | 40.98 | 59.80 | 49.72 | 52.19 | 206.11 | 381.96 | -2.09 | 0.228 |
| *Thermomonas* | 38.81 | 9.74 | 27.38 | 7.17 | 371.96 | 373.12 | -3.31 | 0.205 |
| *Corynebacterium* | 215.95 | 429.53 | 230.97 | 405.52 | 382.08 | 354.12 | -0.38 | 0.327 |
| *Kaistia* | 264.30 | 425.94 | 513.51 | 276.21 | 189.14 | 346.06 | 0.57 | 0.215 |
| *Anatilimnocola* | 64.18 | 83.21 | 187.63 | 166.23 | 301.31 | 344.75 | -1.28 | 0.080 |
| *Planifilum* | 152.64 | 1369.07 | 1333.59 | 510.52 | 138.25 | 344.23 | 1.52 | 0.257 |
| *Gordonia* | 94.96 | 145.74 | 135.39 | 121.05 | 191.93 | 343.71 | -0.80 | 0.288 |
| *Altererythrobacter* | 160.88 | 417.06 | 527.79 | 594.80 | 440.07 | 342.67 | -0.32 | 0.532 |
| *Pseudaminobacter* | 287.28 | 72.61 | 153.03 | 83.97 | 225.86 | 340.07 | -0.34 | 0.665 |
| *Agromyces* | 184.94 | 160.95 | 393.24 | 289.46 | 153.44 | 334.09 | -0.07 | 0.898 |
| *Schlesneria* | 960.48 | 1588.96 | 2145.60 | 4616.50 | 249.66 | 333.57 | -0.15 | 0.918 |
| *Solirubrobacter* | 238.71 | 158.38 | 215.35 | 141.61 | 285.11 | 331.74 | -0.31 | 0.497 |
| *Arthrobacter* | 392.87 | 329.41 | 509.31 | 370.62 | 303.84 | 330.18 | 0.29 | 0.284 |
| *Capsulimonas* | 79.57 | 6.32 | 16.29 | 9.50 | 393.73 | 329.14 | -2.84 | 0.216 |
| *Aeromicrobium* | 119.46 | 83.38 | 121.28 | 70.57 | 209.91 | 326.28 | -0.90 | 0.330 |
| *Limnoglobus* | 59.41 | 55.36 | 122.46 | 66.52 | 260.04 | 326.02 | -1.46 | 0.212 |
| *Methylobacterium* | 278.82 | 386.13 | 377.28 | 232.59 | 307.39 | 324.72 | 0.27 | 0.253 |
| *Sinorhizobium* | 216.60 | 375.88 | 530.14 | 339.78 | 238.01 | 322.38 | 0.32 | 0.505 |
| *Aquihabitans* | 45.96 | 18.45 | 23.69 | 154.39 | 243.84 | 320.56 | -3.03 | 0.044 |

| | | | | | | | | |
|---|---|---|---|---|---|---|---|---|
| *Labilithrix* | 42.06 | 10.93 | 14.78 | 8.88 | 273.46 | 319.78 | -3.15 | 0.206 |
| *Microvirga* | 205.32 | 215.11 | 408.86 | 177.76 | 273.97 | 314.83 | 0.11 | 0.802 |
| *Reyranella* | 135.94 | 111.23 | 187.46 | 289.92 | 206.87 | 297.92 | -0.87 | 0.034 |
| *Aquisphaera* | 87.38 | 195.63 | 383.83 | 148.78 | 218.51 | 297.14 | 0.01 | 0.993 |
| *Caulobacter* | 181.69 | 184.18 | 221.06 | 142.55 | 341.32 | 295.84 | -0.41 | 0.398 |
| *Phenylobacterium* | 169.33 | 320.70 | 275.82 | 277.30 | 333.22 | 293.76 | -0.24 | 0.418 |
| *Allocatelliglobosispora* | 7.37 | 10.08 | 7.90 | 7.01 | 57.48 | 291.42 | -3.81 | 0.335 |
| *Bordetella* | 2092.90 | 102.51 | 284.05 | 71.97 | 250.42 | 289.33 | 2.02 | 0.431 |
| *Rhabdothermincola* | 69.38 | 111.23 | 55.43 | 27.26 | 198.01 | 287.77 | -1.12 | 0.350 |
| *Pirellula* | 45.10 | 160.09 | 315.63 | 274.50 | 261.31 | 286.47 | -0.66 | 0.328 |
| *Nocardiopsis* | 186.89 | 548.62 | 295.81 | 325.13 | 208.89 | 282.83 | 0.34 | 0.580 |
| *Plasmopara* | 156.97 | 241.42 | 91.72 | 442.29 | 319.29 | 279.97 | -1.09 | 0.050 |
| *Chlorella* | 51.38 | 95.00 | 152.19 | 98.15 | 142.81 | 276.84 | -0.79 | 0.315 |
| *Pseudoxanthomonas* | 111.66 | 58.60 | 85.00 | 58.58 | 229.40 | 273.46 | -1.14 | 0.257 |
| *Pseudorhodoplanes* | 184.94 | 75.86 | 184.78 | 69.17 | 254.98 | 273.46 | -0.42 | 0.545 |
| *Luteimonas* | 122.50 | 36.73 | 60.81 | 34.90 | 278.27 | 271.64 | -1.41 | 0.265 |
| *Vibrio* | 197.08 | 266.02 | 170.67 | 340.09 | 281.06 | 271.38 | -0.49 | 0.078 |
| *Blastopirellula* | 54.42 | 100.98 | 239.03 | 186.48 | 214.97 | 259.67 | -0.74 | 0.245 |
| *Actinomycetospora* | 197.73 | 212.89 | 239.87 | 190.37 | 207.88 | 256.29 | -0.01 | 0.961 |
| *Sorangium* | 73.72 | 19.31 | 40.15 | 21.50 | 259.28 | 254.99 | -2.01 | 0.227 |
| *Tautonia* | 84.12 | 154.45 | 444.47 | 129.30 | 187.62 | 247.70 | 0.28 | 0.760 |
| *Prauserella* | 187.11 | 275.42 | 219.04 | 218.73 | 181.80 | 246.40 | 0.08 | 0.734 |
| *Rhizopus* | 5717.80 | 68.00 | 1162.08 | 1075.88 | 186.10 | 236.25 | 2.21 | 0.404 |
| *Sporichthya* | 29.49 | 21.87 | 20.66 | 17.45 | 198.51 | 228.45 | -2.62 | 0.200 |
| *Aestuariivirga* | 330.86 | 1255.11 | 2347.85 | 1001.88 | 129.64 | 227.93 | 1.53 | 0.280 |
| *Aminobacter* | 140.93 | 76.20 | 136.06 | 80.08 | 162.30 | 223.77 | -0.40 | 0.479 |
| *Zavarzinella* | 50.30 | 43.06 | 109.86 | 71.66 | 180.53 | 222.46 | -1.22 | 0.171 |
| *Saccharomonospora* | 197.73 | 249.62 | 216.19 | 219.35 | 170.66 | 221.94 | 0.12 | 0.486 |
| *Streptacidiphilus* | 123.58 | 250.82 | 161.60 | 179.47 | 165.60 | 213.36 | -0.06 | 0.868 |
| *Saccharothrix* | 145.48 | 208.27 | 175.20 | 162.64 | 166.35 | 210.50 | -0.03 | 0.892 |
| *Herbaspirillum* | 151.34 | 317.96 | 716.94 | 366.26 | 81.03 | 206.07 | 0.86 | 0.414 |
| *Roseomonas* | 87.38 | 109.52 | 182.26 | 152.83 | 166.35 | 200.61 | -0.45 | 0.241 |
| *Kitasatospora* | 154.80 | 259.87 | 163.44 | 191.15 | 174.20 | 200.09 | 0.03 | 0.911 |
| *Sciscionella* | 136.59 | 185.38 | 185.11 | 166.23 | 150.91 | 197.49 | -0.02 | 0.914 |
| *Blastococcus* | 125.53 | 163.17 | 141.10 | 145.82 | 152.18 | 196.71 | -0.20 | 0.333 |
| *Nakamurella* | 87.59 | 112.76 | 139.25 | 88.49 | 154.45 | 195.92 | -0.37 | 0.413 |
| *Chthoniobacter* | 14.31 | 10.25 | 25.20 | 16.05 | 211.68 | 195.40 | -3.09 | 0.185 |
| *Enterococcus* | 125.32 | 203.49 | 105.66 | 278.40 | 218.51 | 194.62 | -0.67 | 0.095 |
| *Brevundimonas* | 162.83 | 80.30 | 101.63 | 65.43 | 175.22 | 192.80 | -0.33 | 0.570 |
| *Actinokineospora* | 133.77 | 189.31 | 167.98 | 151.74 | 145.09 | 188.90 | 0.02 | 0.934 |
| *Actinopolyspora* | 184.94 | 158.55 | 139.59 | 146.29 | 125.34 | 186.04 | 0.08 | 0.719 |
| *Neorhizobium* | 271.88 | 491.72 | 676.28 | 369.53 | 154.71 | 185.52 | 1.02 | 0.163 |
| *Actinoallomurus* | 208.36 | 428.51 | 256.00 | 234.46 | 135.21 | 185.00 | 0.69 | 0.228 |
| *Mucilaginibacter* | 488.91 | 215.79 | 423.31 | 376.07 | 239.28 | 184.74 | 0.50 | 0.344 |
| *Gimesia* | 86.73 | 132.58 | 307.40 | 281.67 | 161.04 | 184.22 | -0.25 | 0.691 |
| *Methylovirgula* | 34.91 | 30.92 | 44.18 | 30.69 | 174.71 | 182.65 | -1.82 | 0.201 |
| *Achromobacter* | 5756.18 | 1108.00 | 1425.81 | 1993.79 | 122.55 | 182.39 | 1.85 | 0.316 |
| *Tardiphaga* | 60.27 | 95.85 | 114.39 | 62.00 | 130.15 | 181.35 | -0.46 | 0.437 |
| *Ramlibacter* | 291.40 | 355.38 | 880.21 | 309.09 | 193.45 | 179.53 | 1.16 | 0.267 |
| *Opitutus* | 56.59 | 14.35 | 26.04 | 23.84 | 130.15 | 178.49 | -1.78 | 0.223 |
| *Kutzneria* | 135.07 | 166.41 | 147.15 | 142.86 | 132.68 | 177.45 | -0.01 | 0.938 |
| *Azospirillum* | 133.77 | 331.46 | 174.36 | 118.87 | 172.18 | 167.04 | 0.48 | 0.423 |
| *Pelagibacterium* | 75.23 | 106.27 | 174.03 | 63.09 | 137.24 | 166.26 | -0.04 | 0.936 |
| *Streptosporangium* | 102.34 | 264.83 | 135.39 | 168.72 | 131.67 | 164.18 | 0.11 | 0.824 |
| *Microbispora* | 116.21 | 258.50 | 152.02 | 167.63 | 137.24 | 161.32 | 0.18 | 0.685 |
| *Acidothermus* | 17.78 | 23.75 | 24.52 | 12.93 | 158.51 | 160.54 | -2.33 | 0.211 |
| *Micromonas* | 89.11 | 163.51 | 62.82 | 235.55 | 188.38 | 159.50 | -0.89 | 0.082 |
| *Gottfriedia* | 1.73 | 6.49 | 4.70 | 2.96 | 50.89 | 155.59 | -4.02 | 0.283 |
| *Rubinisphaera* | 72.42 | 117.38 | 277.67 | 222.16 | 140.27 | 155.33 | -0.15 | 0.821 |
| *Variovorax* | 133.99 | 197.51 | 232.32 | 169.81 | 151.42 | 154.55 | 0.25 | 0.414 |
| *Enterobacter* | 76.54 | 56.21 | 44.01 | 82.72 | 94.70 | 154.29 | -0.91 | 0.130 |
| *Castellaniella* | 23.20 | 11.79 | 22.85 | 13.55 | 123.82 | 153.51 | -2.33 | 0.209 |
| *Catellatospora* | 48.78 | 45.11 | 58.62 | 43.62 | 72.92 | 151.43 | -0.81 | 0.354 |
| *Cupriavidus* | 181.26 | 153.09 | 185.62 | 129.15 | 138.25 | 151.17 | 0.31 | 0.059 |
| *Thermomonospora* | 92.15 | 312.67 | 170.83 | 192.24 | 117.23 | 149.61 | 0.33 | 0.616 |
| *Aureliella* | 27.75 | 74.15 | 411.89 | 177.44 | 171.17 | 149.09 | 0.05 | 0.969 |
| *Polyangium* | 28.84 | 10.76 | 16.80 | 12.46 | 155.47 | 148.05 | -2.48 | 0.202 |
| *Methylocapsa* | 33.39 | 37.08 | 53.42 | 39.26 | 130.91 | 147.01 | -1.36 | 0.191 |
| *Lentzea* | 98.87 | 140.61 | 135.06 | 116.06 | 107.11 | 146.23 | 0.02 | 0.924 |
| *Batrachochytrium* | 250.20 | 193.58 | 197.71 | 246.30 | 150.40 | 143.63 | 0.25 | 0.438 |
| *Minicystis* | 19.95 | 9.40 | 13.77 | 12.93 | 144.33 | 142.85 | -2.80 | 0.187 |
| *Trebonia* | 45.53 | 135.83 | 69.04 | 77.27 | 107.11 | 142.59 | -0.38 | 0.488 |
| *Iamia* | 26.23 | 10.76 | 12.93 | 17.92 | 103.31 | 141.28 | -2.39 | 0.190 |
| *Jatrophihabitans* | 23.42 | 36.39 | 38.97 | 22.12 | 128.63 | 140.50 | -1.56 | 0.229 |
| *Phycicoccus* | 212.04 | 54.67 | 95.75 | 38.48 | 118.50 | 140.50 | 0.29 | 0.722 |
| *Nitratireductor* | 127.49 | 101.49 | 185.79 | 100.95 | 106.09 | 139.98 | 0.26 | 0.475 |
| *Chelatococcus* | 81.96 | 101.66 | 153.03 | 85.06 | 122.04 | 139.72 | -0.04 | 0.906 |
| *Niastella* | 155.67 | 21.02 | 13.77 | 10.59 | 86.34 | 137.12 | -0.29 | 0.820 |
| *Frateuria* | 323.92 | 598.17 | 701.82 | 363.61 | 757.33 | 136.34 | 0.37 | 0.603 |
| *Staphylococcus* | 91.50 | 142.84 | 172.01 | 252.38 | 136.48 | 136.08 | -0.37 | 0.443 |
| *Lacipirellula* | 31.00 | 104.05 | 141.77 | 80.54 | 95.96 | 135.82 | -0.17 | 0.767 |
| *Hoeflea* | 124.02 | 160.78 | 229.96 | 130.08 | 98.50 | 135.56 | 0.50 | 0.241 |
| *Bauldia* | 70.46 | 32.63 | 67.53 | 29.60 | 113.94 | 135.04 | -0.71 | 0.385 |
| *Pandoraea* | 485.01 | 6258.44 | 5084.90 | 5341.70 | 182.31 | 134.00 | 1.06 | 0.452 |
| *Acidisphaera* | 42.93 | 57.41 | 44.51 | 74.78 | 164.84 | 133.48 | -1.36 | 0.098 |
| *Telmatocola* | 22.55 | 28.19 | 59.46 | 51.57 | 116.98 | 133.48 | -1.45 | 0.108 |
| *Luteolibacter* | 11.06 | 10.25 | 16.63 | 20.10 | 73.68 | 133.22 | -2.58 | 0.193 |
| *Thermogemmata* | 19.30 | 14.86 | 33.09 | 21.81 | 132.93 | 131.92 | -2.09 | 0.183 |
| *Aureimonas* | 120.55 | 169.66 | 247.94 | 162.64 | 114.70 | 131.14 | 0.40 | 0.367 |
| *Patulibacter* | 27.97 | 24.43 | 35.11 | 17.60 | 104.32 | 129.32 | -1.52 | 0.247 |
| *Flavobacterium* | 162.18 | 59.97 | 158.74 | 62.94 | 64.31 | 126.97 | 0.58 | 0.356 |
| *Lignipirellula* | 18.65 | 42.37 | 99.44 | 88.18 | 106.60 | 126.71 | -1.00 | 0.141 |
| *Roseiarcus* | 16.26 | 21.70 | 27.88 | 14.18 | 113.18 | 125.93 | -1.94 | 0.218 |
| *Dyella* | 261.26 | 1111.76 | 907.76 | 852.63 | 474.00 | 125.93 | 0.65 | 0.454 |
| *Geodermatophilus* | 75.02 | 113.28 | 90.88 | 76.34 | 107.61 | 124.37 | -0.14 | 0.619 |
| *Paracoccus* | 96.48 | 147.62 | 205.27 | 118.71 | 109.89 | 124.11 | 0.35 | 0.412 |
| *Aquamicrobium* | 86.51 | 53.14 | 83.32 | 51.88 | 77.99 | 121.77 | -0.17 | 0.707 |
| *Rhodopirellula* | 40.33 | 96.19 | 208.13 | 169.03 | 111.66 | 121.25 | -0.22 | 0.744 |
| *Actinospica* | 35.56 | 44.08 | 38.64 | 27.89 | 65.07 | 120.73 | -0.85 | 0.359 |
| *Kibdelosporangium* | 77.84 | 123.02 | 98.10 | 93.16 | 91.66 | 117.87 | -0.02 | 0.943 |

| | | | | | | | | |
|---|---|---|---|---|---|---|---|---|
| *Noviherbaspirillum* | 257.79 | 74.15 | 116.91 | 106.40 | 91.15 | 116.83 | 0.51 | 0.505 |
| *Paraclostridium* | 53.12 | 97.39 | 40.82 | 164.82 | 133.19 | 116.57 | -1.12 | 0.031 |
| *Sphaerisporangium* | 76.75 | 206.05 | 115.91 | 123.23 | 102.29 | 115.79 | 0.22 | 0.668 |
| *Chlamydomonas* | 69.81 | 95.68 | 52.07 | 129.93 | 131.92 | 115.27 | -0.79 | 0.037 |
| *Actinophytocola* | 86.51 | 131.22 | 116.41 | 95.65 | 83.56 | 113.96 | 0.19 | 0.441 |
| *Symmachiella* | 55.94 | 68.68 | 188.14 | 120.27 | 113.44 | 113.96 | -0.15 | 0.808 |
| *Modestobacter* | 69.81 | 110.37 | 82.14 | 106.56 | 97.74 | 113.70 | -0.28 | 0.259 |
| *Methylocystis* | 39.89 | 36.73 | 57.45 | 32.09 | 132.68 | 112.66 | -1.05 | 0.258 |
| *Baekduia* | 42.50 | 86.28 | 71.56 | 32.09 | 81.03 | 110.06 | -0.15 | 0.790 |
| *Archangium* | 42.71 | 17.09 | 20.49 | 20.41 | 89.13 | 109.80 | -1.45 | 0.223 |
| *Maioricimonas* | 41.84 | 87.48 | 184.95 | 144.88 | 83.05 | 109.28 | -0.10 | 0.879 |
| *Sandaracinus* | 17.13 | 5.98 | 8.73 | 6.85 | 94.95 | 108.50 | -2.72 | 0.202 |
| *Planctomicrobium* | 40.54 | 84.40 | 176.55 | 153.92 | 86.60 | 108.50 | -0.21 | 0.747 |
| *Acidovorax* | 166.73 | 39.98 | 78.78 | 43.62 | 91.41 | 108.50 | 0.23 | 0.761 |
| *Mycobacteroides* | 54.64 | 103.37 | 107.17 | 65.59 | 88.11 | 108.50 | 0.02 | 0.963 |
| *Xanthomonas* | 603.18 | 72.78 | 186.79 | 140.05 | 111.41 | 108.24 | 1.26 | 0.408 |
| *Chelativorans* | 79.14 | 50.74 | 93.73 | 53.75 | 85.58 | 107.98 | -0.14 | 0.718 |
| *Ilumatobacter* | 38.81 | 13.16 | 17.47 | 13.40 | 86.09 | 107.72 | -1.58 | 0.245 |
| *Tuwongella* | 23.42 | 19.82 | 49.22 | 30.69 | 104.32 | 107.20 | -1.39 | 0.175 |
| *Actinomarinicola* | 23.42 | 10.42 | 10.08 | 8.88 | 82.04 | 105.90 | -2.16 | 0.221 |
| *Sphingosinicella* | 71.98 | 34.17 | 51.57 | 35.21 | 100.52 | 105.90 | -0.61 | 0.352 |
| *Demequina* | 29.05 | 24.26 | 49.89 | 31.16 | 109.13 | 103.82 | -1.24 | 0.196 |
| *Cryobacterium* | 135.29 | 145.06 | 370.39 | 236.80 | 72.16 | 103.82 | 0.66 | 0.444 |
| *Enhydrobacter* | 68.95 | 30.75 | 71.56 | 101.42 | 69.38 | 103.30 | -0.68 | 0.119 |
| *Pedobacter* | 86.94 | 73.81 | 121.11 | 74.93 | 141.54 | 102.00 | -0.18 | 0.640 |
| *Ochrobactrum* | 87.38 | 38.10 | 51.57 | 159.68 | 78.24 | 101.47 | -0.94 | 0.145 |
| *Novibacillus* | 58.32 | 296.26 | 448.84 | 78.05 | 53.93 | 101.21 | 1.78 | 0.235 |
| *Allosaccharopolyspora* | 90.19 | 84.40 | 69.04 | 86.15 | 53.93 | 99.91 | 0.02 | 0.939 |
| *Listeria* | 51.82 | 122.50 | 56.11 | 147.84 | 109.89 | 98.61 | -0.63 | 0.211 |
| *Brevibacillus* | 86.51 | 980.71 | 1195.51 | 748.72 | 68.36 | 98.61 | 1.30 | 0.340 |
| *Parachlamydia* | 39.89 | 28.19 | 45.35 | 45.80 | 120.78 | 97.57 | -1.22 | 0.145 |
| *Gandjariella* | 77.62 | 93.12 | 78.11 | 82.10 | 74.95 | 97.31 | -0.03 | 0.840 |
| *Halopolyspora* | 96.27 | 77.74 | 65.85 | 79.61 | 58.24 | 96.79 | 0.03 | 0.906 |
| *Nostoc* | 39.89 | 86.79 | 80.80 | 76.49 | 76.47 | 96.53 | -0.27 | 0.455 |
| *Jiangella* | 59.41 | 93.63 | 89.87 | 68.24 | 69.63 | 94.97 | 0.06 | 0.819 |
| *Gemmatirosa* | 5.64 | 18.79 | 30.40 | 51.41 | 77.23 | 94.71 | -2.03 | 0.027 |
| *Corallococcus* | 61.14 | 41.01 | 40.82 | 52.03 | 94.44 | 94.71 | -0.75 | 0.132 |
| *Rhizoctonia* | 49.43 | 81.67 | 34.10 | 138.03 | 98.24 | 93.41 | -1.00 | 0.052 |
| *Chondromyces* | 21.46 | 7.01 | 12.26 | 11.22 | 86.09 | 91.85 | -2.21 | 0.194 |
| *Planctopirus* | 89.11 | 134.63 | 248.61 | 277.46 | 79.00 | 91.07 | 0.08 | 0.924 |
| *Thalassoglobus* | 36.86 | 77.74 | 173.86 | 158.75 | 76.72 | 90.55 | -0.18 | 0.808 |
| *Anaeromyxobacter* | 19.73 | 15.89 | 20.33 | 15.11 | 88.62 | 90.29 | -1.79 | 0.204 |
| *Coccomyxa* | 50.95 | 60.31 | 44.85 | 69.95 | 81.03 | 90.03 | -0.63 | 0.020 |
| *Actinopolymorpha* | 81.09 | 122.50 | 82.31 | 85.06 | 78.49 | 89.77 | 0.18 | 0.510 |
| *Trinickia* | 334.33 | 19.65 | 21.17 | 12.31 | 320.05 | 89.25 | -0.17 | 0.919 |
| *Pasteurella* | 43.36 | 72.44 | 30.91 | 131.95 | 101.28 | 88.47 | -1.13 | 0.031 |
| *Ralstonia* | 447.50 | 1965.18 | 1096.90 | 1251.14 | 71.66 | 87.42 | 1.31 | 0.301 |
| *Alicyclobacillus* | 55.94 | 86.45 | 79.96 | 52.35 | 71.66 | 87.42 | 0.07 | 0.801 |
| *Gemmatimonas* | 6.72 | 21.87 | 31.24 | 49.23 | 70.90 | 86.90 | -1.79 | 0.025 |
| *Hansschlegelia* | 26.67 | 17.60 | 40.48 | 18.54 | 56.97 | 86.90 | -0.94 | 0.320 |
| *Salmonella* | 75.02 | 108.32 | 57.11 | 127.44 | 97.23 | 85.86 | -0.37 | 0.300 |
| *Candidatus* Sulfopaludibacter | 9.76 | 9.23 | 16.13 | 11.53 | 71.91 | 85.60 | -2.27 | 0.188 |
| *Caballeronia* | 42.71 | 25.80 | 39.48 | 26.48 | 93.94 | 85.60 | -0.93 | 0.261 |
| *Ancylobacter* | 56.81 | 58.95 | 93.56 | 49.23 | 79.25 | 85.60 | -0.03 | 0.929 |
| *Dactylosporangium* | 43.80 | 53.99 | 52.91 | 50.63 | 56.21 | 84.56 | -0.34 | 0.324 |
| *Xanthobacter* | 50.95 | 58.43 | 87.18 | 49.07 | 84.32 | 84.56 | -0.15 | 0.682 |
| *Labilibacter* | 45.31 | 79.28 | 46.70 | 98.77 | 73.94 | 84.30 | -0.58 | 0.109 |
| *Sporocytophaga* | 31.00 | 120.28 | 87.52 | 47.83 | 412.47 | 83.78 | -1.19 | 0.475 |
| *Paenarthrobacter* | 58.11 | 52.11 | 65.68 | 88.49 | 75.45 | 83.52 | -0.49 | 0.012 |
| *Halosaccharopolyspora* | 80.00 | 73.47 | 55.60 | 63.41 | 49.63 | 83.26 | 0.09 | 0.743 |
| *Hephaestia* | 648.27 | 112.08 | 88.69 | 45.65 | 199.02 | 83.00 | 1.38 | 0.444 |
| *Ciceribacter* | 109.92 | 254.57 | 339.99 | 218.73 | 61.28 | 79.62 | 0.97 | 0.247 |
| *Antrihabitans* | 17.78 | 19.48 | 22.17 | 16.20 | 75.96 | 79.10 | -1.53 | 0.210 |
| *Actinocorallia* | 46.40 | 149.50 | 90.71 | 85.68 | 66.85 | 78.58 | 0.31 | 0.600 |
| *Actinocatenispora* | 50.30 | 112.59 | 70.22 | 74.47 | 62.54 | 78.58 | 0.11 | 0.783 |
| *Nitrospira* | 78.70 | 20.50 | 38.80 | 29.44 | 64.82 | 77.28 | -0.31 | 0.646 |
| *Actinomyces* | 23.42 | 35.88 | 35.11 | 47.67 | 58.24 | 77.02 | -0.95 | 0.056 |
| *Aquibium* | 59.62 | 45.45 | 74.08 | 40.51 | 54.44 | 76.76 | 0.06 | 0.860 |
| *Acetobacter* | 57.24 | 71.76 | 51.91 | 72.13 | 72.42 | 76.24 | -0.29 | 0.150 |
| *Chitinophaga* | 48.78 | 45.11 | 68.87 | 48.61 | 94.95 | 76.24 | -0.43 | 0.301 |
| *Kinneretia* | 22.11 | 48.35 | 66.69 | 46.58 | 54.19 | 76.24 | -0.37 | 0.450 |
| *Actinoalloteichus* | 57.02 | 72.96 | 60.81 | 65.12 | 58.49 | 75.20 | -0.06 | 0.718 |
| *Synechococcus* | 39.68 | 65.27 | 44.01 | 77.12 | 79.76 | 74.15 | -0.63 | 0.069 |
| *Glaciibacter* | 20.60 | 15.04 | 34.44 | 26.64 | 33.42 | 74.15 | -0.94 | 0.284 |
| *Bythopirellula* | 10.19 | 18.79 | 39.31 | 27.42 | 59.00 | 73.37 | -1.23 | 0.143 |
| *Halomonas* | 57.89 | 54.67 | 71.39 | 56.86 | 71.15 | 73.11 | -0.13 | 0.473 |
| *Planotetraspora* | 44.88 | 120.62 | 66.69 | 75.40 | 48.62 | 73.11 | 0.24 | 0.665 |
| *Starkeya* | 34.69 | 31.27 | 54.76 | 25.86 | 56.21 | 72.59 | -0.36 | 0.518 |
| *Beijerinckia* | 40.54 | 64.58 | 62.49 | 45.96 | 70.14 | 72.59 | -0.17 | 0.574 |
| *Martelella* | 102.34 | 181.79 | 243.57 | 112.32 | 58.74 | 72.33 | 1.12 | 0.133 |
| *Shewanella* | 36.21 | 54.33 | 31.92 | 78.05 | 77.99 | 71.81 | -0.90 | 0.028 |
| *Micractinium* | 9.76 | 23.58 | 36.79 | 22.75 | 37.47 | 71.81 | -0.91 | 0.298 |
| *Haloechinothrix* | 44.66 | 70.73 | 62.99 | 59.82 | 59.50 | 71.81 | -0.10 | 0.660 |
| *Roseimicrobium* | 11.49 | 10.76 | 24.52 | 16.51 | 55.20 | 71.29 | -1.61 | 0.180 |
| *Rubrivivax* | 24.93 | 27.51 | 39.48 | 20.41 | 46.84 | 71.29 | -0.59 | 0.404 |
| *Acrocarpospora* | 47.27 | 105.93 | 64.67 | 64.19 | 55.20 | 71.03 | 0.19 | 0.655 |
| *Calocera* | 36.86 | 63.73 | 25.53 | 100.17 | 77.23 | 70.51 | -0.97 | 0.052 |
| *Mycolicibacter* | 26.02 | 47.67 | 37.46 | 26.80 | 50.39 | 70.25 | -0.41 | 0.453 |
| *Streptomonospora* | 37.08 | 147.96 | 69.54 | 83.97 | 55.70 | 69.99 | 0.28 | 0.698 |
| *Miltoncostaea* | 7.59 | 3.93 | 7.56 | 5.45 | 34.94 | 69.73 | -2.53 | 0.243 |
| *Rhizomicrobium* | 118.38 | 88.84 | 56.27 | 62.63 | 114.95 | 69.73 | 0.09 | 0.833 |
| *Putridiphycobacter* | 36.86 | 65.61 | 26.37 | 112.32 | 86.85 | 69.47 | -1.06 | 0.053 |
| *Shigella* | 42.28 | 66.46 | 30.24 | 111.54 | 80.27 | 69.47 | -0.91 | 0.072 |
| *Catenulispora* | 29.27 | 50.74 | 36.62 | 38.32 | 62.79 | 68.69 | -0.54 | 0.199 |
| *Proteus* | 52.47 | 87.31 | 40.32 | 98.93 | 74.44 | 68.69 | -0.43 | 0.298 |
| *Haliangium* | 7.37 | 3.76 | 10.08 | 4.52 | 73.94 | 67.91 | -2.79 | 0.200 |
| *Aquicella* | 53.99 | 273.88 | 401.64 | 17.45 | 55.20 | 67.65 | 2.38 | 0.190 |
| *Roseimaritima* | 17.56 | 45.79 | 118.76 | 91.76 | 68.36 | 67.65 | -0.32 | 0.668 |
| *Croceibacterium* | 43.36 | 100.63 | 127.50 | 128.99 | 71.40 | 67.65 | 0.02 | 0.973 |
| *Myxococcus* | 24.50 | 18.79 | 25.53 | 22.90 | 69.63 | 66.87 | -1.21 | 0.182 |

| | | | | | | | | |
|---|---|---|---|---|---|---|---|---|
| *Curtobacterium* | 106.67 | 107.13 | 274.98 | 194.42 | 53.93 | 66.87 | 0.63 | 0.468 |
| *Tsuneonella* | 30.79 | 41.01 | 50.23 | 39.57 | 81.03 | 66.61 | -0.62 | 0.208 |
| *Pedococcus* | 76.10 | 16.74 | 36.28 | 12.00 | 47.10 | 66.61 | 0.04 | 0.962 |
| *Polaromonas* | 40.33 | 24.77 | 50.39 | 22.12 | 43.55 | 66.09 | -0.19 | 0.737 |
| *Cryptosporangium* | 34.69 | 62.02 | 54.43 | 43.31 | 51.40 | 65.83 | -0.09 | 0.781 |
| *Kineosporia* | 32.09 | 43.74 | 45.69 | 35.52 | 46.34 | 65.57 | -0.28 | 0.443 |
| *Allonocardiopsis* | 44.66 | 173.76 | 75.93 | 104.69 | 53.68 | 65.57 | 0.39 | 0.619 |
| *Rhodovulum* | 45.75 | 31.27 | 59.30 | 29.60 | 71.66 | 65.31 | -0.29 | 0.555 |
| *Leptospira* | 39.03 | 21.87 | 36.96 | 24.77 | 58.24 | 65.05 | -0.60 | 0.313 |
| *Deinococcus* | 27.75 | 32.12 | 50.06 | 31.63 | 60.52 | 64.53 | -0.51 | 0.287 |
| *Roseibium* | 44.01 | 55.87 | 102.13 | 51.57 | 62.03 | 63.75 | 0.19 | 0.690 |
| *Cellvibrio* | 42.71 | 8.88 | 7.39 | 5.76 | 26.33 | 63.49 | -0.69 | 0.588 |
| *Gellertiella* | 1016.42 | 2642.28 | 2910.41 | 2733.16 | 289.92 | 63.23 | 1.09 | 0.335 |
| *Bailinhaonella* | 32.52 | 122.33 | 61.31 | 69.17 | 45.32 | 62.45 | 0.29 | 0.675 |
| *Phytoactinopolyspora* | 32.74 | 59.80 | 50.56 | 44.24 | 51.40 | 61.93 | -0.14 | 0.642 |
| *Variibacter* | 47.05 | 39.98 | 61.65 | 18.85 | 65.83 | 61.93 | 0.02 | 0.967 |
| *Methylophilus* | 235.03 | 97.73 | 264.23 | 156.10 | 59.76 | 60.62 | 1.11 | 0.166 |
| *Hymenobacter* | 29.49 | 39.81 | 58.29 | 43.00 | 63.81 | 60.62 | -0.39 | 0.283 |
| *Labrys* | 32.52 | 59.46 | 64.67 | 33.49 | 51.65 | 60.62 | 0.10 | 0.789 |
| *Granulicella* | 16.04 | 15.38 | 23.85 | 16.20 | 66.59 | 59.58 | -1.36 | 0.204 |
| *Motilibacter* | 25.37 | 59.12 | 37.46 | 39.10 | 47.60 | 59.32 | -0.26 | 0.532 |
| *Stappia* | 48.78 | 35.37 | 58.62 | 62.32 | 57.48 | 58.80 | -0.32 | 0.215 |
| *Rubripirellula* | 15.61 | 39.64 | 93.56 | 83.35 | 58.24 | 58.80 | -0.43 | 0.542 |
| *Youhaiella* | 6.07 | 6.15 | 10.75 | 20.56 | 54.95 | 58.54 | -2.54 | 0.090 |
| *Phytohabitans* | 31.44 | 55.02 | 67.19 | 69.48 | 41.27 | 58.54 | -0.14 | 0.717 |
| *Pedosphaera* | 33.17 | 16.23 | 48.38 | 26.02 | 52.67 | 58.28 | -0.49 | 0.392 |
| *Plantactinospora* | 29.70 | 48.86 | 64.34 | 73.69 | 34.44 | 58.02 | -0.22 | 0.637 |
| *Candidatus* Microthrix | 10.41 | 5.98 | 5.38 | 8.41 | 41.78 | 57.76 | -2.31 | 0.185 |
| *Yinghuangia* | 26.67 | 52.62 | 37.12 | 36.45 | 43.30 | 57.76 | -0.24 | 0.515 |
| *Oleiharenicola* | 20.81 | 6.32 | 12.93 | 10.44 | 41.27 | 57.24 | -1.44 | 0.231 |
| *Diversispora* | 29.70 | 40.83 | 16.80 | 73.07 | 57.22 | 56.98 | -1.10 | 0.022 |
| *Cytophaga* | 264.08 | 39.13 | 19.32 | 8.88 | 33.17 | 56.98 | 1.71 | 0.443 |
| *Steroidobacter* | 21.46 | 25.29 | 34.94 | 31.63 | 51.40 | 56.72 | -0.77 | 0.109 |
| *Virgisporangium* | 26.67 | 42.03 | 39.64 | 34.74 | 34.44 | 56.46 | -0.21 | 0.550 |
| *Nitrosomonas* | 76.97 | 16.91 | 23.18 | 15.11 | 42.54 | 56.46 | 0.04 | 0.966 |
| *Leucobacter* | 52.25 | 47.67 | 125.82 | 79.30 | 38.99 | 55.68 | 0.38 | 0.582 |
| *Asanoa* | 28.40 | 50.06 | 52.24 | 46.27 | 35.95 | 55.68 | -0.08 | 0.815 |
| *Microtetraspora* | 28.62 | 80.30 | 55.60 | 58.73 | 37.73 | 55.16 | 0.12 | 0.810 |
| *Thalassospira* | 20.60 | 24.60 | 41.32 | 19.94 | 17.98 | 55.16 | -0.10 | 0.884 |
| *Paralcaligenes* | 20.16 | 561.09 | 647.90 | 643.41 | 38.99 | 54.64 | 0.74 | 0.590 |
| *Planosporangium* | 39.46 | 91.07 | 104.65 | 99.24 | 51.65 | 54.64 | 0.19 | 0.716 |
| *Gaiella* | 35.99 | 4.95 | 10.92 | 4.52 | 56.72 | 54.38 | -1.15 | 0.353 |
| *Prosthecobacter* | 33.39 | 18.11 | 45.19 | 34.27 | 42.54 | 54.12 | -0.44 | 0.311 |
| *Skermania* | 9.32 | 14.18 | 13.77 | 9.35 | 51.15 | 53.60 | -1.61 | 0.215 |
| *Stieleria* | 18.00 | 40.49 | 82.98 | 69.48 | 48.36 | 53.60 | -0.28 | 0.660 |
| *Gryllotalpicola* | 35.34 | 24.26 | 45.02 | 29.60 | 49.88 | 53.08 | -0.34 | 0.384 |
| *Tetrasphaera* | 43.80 | 32.80 | 44.18 | 26.80 | 42.54 | 53.08 | -0.02 | 0.956 |
| *Brevibacterium* | 33.39 | 53.31 | 61.14 | 47.67 | 40.77 | 52.82 | 0.07 | 0.822 |
| *Xylella* | 37.51 | 65.95 | 30.24 | 69.64 | 54.95 | 52.56 | -0.41 | 0.322 |
| *Hypericibacter* | 23.42 | 20.33 | 29.56 | 15.58 | 51.40 | 52.30 | -0.70 | 0.332 |
| *Crossiella* | 38.59 | 45.11 | 39.48 | 40.66 | 35.20 | 52.30 | -0.06 | 0.785 |
| *Nevskia* | 11.49 | 6.49 | 14.45 | 7.95 | 40.26 | 52.04 | -1.63 | 0.226 |
| *Rosistilla* | 14.09 | 34.34 | 79.79 | 64.19 | 47.10 | 52.04 | -0.35 | 0.612 |
| *Magnetospirillum* | 24.72 | 22.21 | 38.13 | 21.65 | 59.00 | 51.52 | -0.63 | 0.304 |
| *Thermobifida* | 37.29 | 100.98 | 239.20 | 72.75 | 37.47 | 51.52 | 1.22 | 0.350 |
| *Trebouxia* | 26.23 | 44.76 | 22.85 | 43.93 | 56.46 | 51.26 | -0.69 | 0.087 |
| *Methylocella* | 14.09 | 21.36 | 30.40 | 18.85 | 49.12 | 51.26 | -0.86 | 0.226 |
| *Nitrosospira* | 16.48 | 13.16 | 11.93 | 17.60 | 44.31 | 51.00 | -1.44 | 0.142 |
| *Priestia* | 55.72 | 91.92 | 127.66 | 42.37 | 39.50 | 50.74 | 1.05 | 0.145 |
| *Abditibacterium* | 2.38 | 3.59 | 68.03 | 35.99 | 79.25 | 50.74 | -1.17 | 0.304 |
| *Hamadaea* | 8.89 | 16.57 | 11.09 | 10.91 | 15.45 | 50.74 | -1.08 | 0.396 |
| *Thermacetogenium* | 0.65 | 0.68 | 2.52 | 0.47 | 48.87 | 50.48 | -4.69 | 0.190 |
| *Fusarium* | 31.65 | 31.27 | 23.18 | 42.22 | 28.36 | 50.48 | -0.49 | 0.205 |
| *Candidatus* Protochlamydia | 18.65 | 19.14 | 21.17 | 37.70 | 68.87 | 50.22 | -1.41 | 0.068 |
| *Leptolyngbya* | 23.42 | 24.26 | 34.77 | 30.22 | 53.68 | 50.22 | -0.70 | 0.127 |
| *Nannocystis* | 14.96 | 15.38 | 35.11 | 11.53 | 46.34 | 50.22 | -0.72 | 0.383 |
| *Pseudorhizobium* | 46.40 | 81.67 | 98.60 | 81.32 | 34.44 | 49.96 | 0.45 | 0.382 |
| *Glycomyces* | 37.08 | 51.94 | 46.87 | 64.65 | 36.71 | 49.44 | -0.15 | 0.626 |
| *Linnemannia* | 25.15 | 43.06 | 19.65 | 78.36 | 68.11 | 48.92 | -1.15 | 0.035 |
| *Rhizorhabdus* | 13.88 | 170.68 | 66.35 | 44.40 | 58.74 | 48.92 | 0.72 | 0.549 |
| *Methylosinus* | 16.69 | 19.82 | 29.23 | 18.38 | 45.32 | 48.66 | -0.77 | 0.241 |
| *Desertimonas* | 22.77 | 11.96 | 12.43 | 8.88 | 39.50 | 48.66 | -1.04 | 0.299 |
| *Marinobacter* | 43.15 | 31.78 | 34.44 | 25.24 | 38.23 | 48.66 | -0.03 | 0.914 |
| *Carbonactinospora* | 23.85 | 67.32 | 35.95 | 40.51 | 38.74 | 48.66 | -0.01 | 0.987 |
| *Mumia* | 31.87 | 144.20 | 39.64 | 23.99 | 30.64 | 48.14 | 1.07 | 0.408 |
| *Candidimonas* | 51.60 | 206.74 | 816.38 | 244.74 | 42.03 | 47.88 | 1.68 | 0.403 |
| *Vineibacter* | 18.43 | 12.30 | 16.97 | 17.45 | 35.95 | 47.62 | -1.08 | 0.175 |
| *Phreatobacter* | 32.09 | 28.02 | 55.43 | 28.98 | 43.80 | 47.62 | -0.06 | 0.885 |
| *Erythrobacter* | 38.16 | 77.91 | 102.30 | 76.96 | 62.29 | 47.09 | 0.23 | 0.642 |
| *Fodinicola* | 24.93 | 65.78 | 32.25 | 36.14 | 31.40 | 47.09 | 0.10 | 0.850 |
| *Posidoniimonas* | 8.24 | 17.26 | 32.25 | 24.61 | 40.77 | 46.57 | -0.95 | 0.133 |
| *Rhodoblastus* | 18.21 | 21.53 | 29.06 | 21.19 | 45.32 | 46.57 | -0.72 | 0.208 |
| *Alsobacter* | 28.84 | 24.60 | 33.76 | 26.33 | 41.53 | 46.05 | -0.38 | 0.272 |
| *Pimelobacter* | 26.88 | 11.62 | 79.29 | 12.77 | 22.28 | 46.05 | 0.54 | 0.629 |
| *Theileria* | 29.27 | 32.98 | 16.46 | 60.91 | 54.44 | 45.79 | -1.03 | 0.015 |
| *Talaromyces* | 12.58 | 18.28 | 30.40 | 10.28 | 28.61 | 45.79 | -0.47 | 0.547 |
| *Auxenochlorella* | 16.04 | 22.04 | 21.50 | 24.30 | 28.87 | 45.53 | -0.73 | 0.172 |
| *Verrucomicrobium* | 5.20 | 8.37 | 17.81 | 11.53 | 27.35 | 45.27 | -1.42 | 0.205 |
| *Croceicoccus* | 27.54 | 69.54 | 138.58 | 90.83 | 44.56 | 45.27 | 0.38 | 0.646 |
| *Rugosimonospora* | 23.42 | 50.57 | 30.57 | 34.74 | 35.95 | 45.27 | -0.15 | 0.699 |
| *Solimonas* | 39.68 | 18.28 | 42.33 | 24.15 | 35.70 | 45.27 | -0.07 | 0.879 |
| *Rhodopila* | 14.96 | 31.44 | 19.65 | 36.61 | 54.44 | 45.01 | -1.04 | 0.030 |
| *Pleomorphomonas* | 43.58 | 69.54 | 108.85 | 44.56 | 39.25 | 45.01 | 0.79 | 0.242 |
| *Georgenia* | 30.35 | 39.30 | 38.97 | 33.18 | 38.49 | 45.01 | -0.10 | 0.584 |
| *Aurantimonas* | 37.94 | 46.64 | 56.44 | 60.45 | 42.79 | 45.01 | -0.07 | 0.772 |
| *Terrabacter* | 64.61 | 25.80 | 33.09 | 23.37 | 51.65 | 44.75 | 0.05 | 0.935 |
| *Methyloceanibacter* | 81.09 | 14.52 | 59.30 | 28.04 | 38.49 | 44.49 | 0.48 | 0.536 |
| *Candidatus* Laterigemmans | 10.19 | 28.02 | 74.41 | 48.76 | 41.78 | 44.23 | -0.26 | 0.737 |
| *Methylopila* | 17.78 | 13.67 | 24.69 | 15.27 | 31.40 | 43.97 | -0.69 | 0.300 |
| *Serratia* | 1431.19 | 47.33 | 387.86 | 735.48 | 95.96 | 43.97 | 1.09 | 0.534 |

| | | | | | | | | |
|---|---|---|---|---|---|---|---|---|
| *Acidobacterium* | 9.97 | 7.69 | 17.64 | 8.57 | 52.67 | 43.71 | -1.57 | 0.223 |
| *Acidiphilium* | 14.31 | 16.40 | 18.98 | 16.05 | 35.95 | 43.45 | -0.94 | 0.200 |
| *Cystobacter* | 10.84 | 9.23 | 11.42 | 11.22 | 35.95 | 43.19 | -1.52 | 0.179 |
| *Pelotomaculum* | 11.49 | 14.69 | 21.17 | 9.19 | 39.25 | 43.19 | -0.95 | 0.301 |
| *Prosthecomicrobium* | 32.31 | 41.01 | 63.33 | 29.29 | 32.92 | 43.19 | 0.38 | 0.383 |
| *Vulcaniibacterium* | 18.00 | 5.64 | 11.93 | 4.99 | 50.64 | 42.93 | -1.47 | 0.273 |
| *Kocuria* | 35.34 | 55.02 | 80.97 | 157.81 | 36.97 | 42.67 | -0.47 | 0.640 |
| *Polystyrenella* | 16.69 | 28.36 | 65.18 | 50.16 | 38.23 | 42.41 | -0.25 | 0.688 |
| *Herbidospora* | 22.33 | 66.80 | 36.28 | 40.51 | 34.18 | 42.15 | 0.10 | 0.848 |
| *Brachybacterium* | 30.57 | 34.17 | 39.98 | 44.56 | 40.51 | 41.89 | -0.28 | 0.096 |
| *Rhodomicrobium* | 20.60 | 18.96 | 28.22 | 17.29 | 42.79 | 41.89 | -0.59 | 0.304 |
| *Fuerstiella* | 19.95 | 32.98 | 71.56 | 59.67 | 40.26 | 41.89 | -0.19 | 0.755 |
| *Aeoliella* | 3.69 | 7.18 | 21.00 | 14.64 | 31.40 | 41.63 | -1.46 | 0.131 |
| *Edaphobacter* | 18.43 | 23.92 | 30.24 | 21.50 | 42.79 | 41.63 | -0.54 | 0.247 |
| *Azorhizobium* | 27.97 | 28.36 | 52.24 | 26.33 | 36.46 | 41.37 | 0.06 | 0.881 |
| *Weizmannia* | 25.37 | 86.79 | 32.92 | 55.93 | 35.70 | 41.11 | 0.13 | 0.855 |
| *Embleya* | 19.51 | 54.50 | 29.06 | 26.48 | 35.70 | 41.11 | 0.00 | 0.997 |
| *Solihabitans* | 31.87 | 36.56 | 33.09 | 32.72 | 28.61 | 40.85 | -0.01 | 0.963 |
| *Stylonychia* | 25.80 | 22.55 | 10.92 | 37.08 | 42.79 | 40.59 | -1.02 | 0.034 |
| *Planomonospora* | 30.35 | 64.58 | 44.01 | 38.32 | 36.46 | 40.33 | 0.27 | 0.508 |
| *Labedaea* | 27.54 | 49.55 | 39.81 | 33.49 | 32.92 | 40.33 | 0.13 | 0.656 |
| *Allokutzneria* | 43.80 | 46.13 | 36.28 | 51.88 | 38.74 | 40.33 | -0.05 | 0.778 |
| *Williamsia* | 18.86 | 21.02 | 27.72 | 18.38 | 31.90 | 40.07 | -0.42 | 0.357 |
| *Actinocrispum* | 26.88 | 43.40 | 35.61 | 32.25 | 28.61 | 40.07 | 0.07 | 0.790 |
| *Guillardia* | 29.92 | 34.34 | 17.97 | 55.31 | 46.08 | 39.81 | -0.78 | 0.043 |
| *Rhodobacter* | 44.01 | 78.25 | 100.12 | 65.43 | 37.73 | 39.81 | 0.64 | 0.247 |
| *Ornithinimicrobium* | 22.11 | 34.17 | 39.31 | 24.77 | 34.44 | 39.81 | -0.05 | 0.875 |
| *Arsenicitalea* | 25.37 | 22.04 | 40.65 | 13.87 | 32.66 | 39.81 | 0.03 | 0.954 |
| *Longimycelium* | 27.32 | 31.61 | 29.73 | 31.31 | 23.80 | 39.55 | -0.09 | 0.709 |
| *Botrimarina* | 4.99 | 9.06 | 21.33 | 19.47 | 27.09 | 39.29 | -1.28 | 0.092 |
| *Duganella* | 61.14 | 44.76 | 59.30 | 39.10 | 55.96 | 39.29 | 0.30 | 0.247 |
| *Rubrobacter* | 18.00 | 21.19 | 21.33 | 15.89 | 33.68 | 39.29 | -0.55 | 0.311 |
| *Scenedesmus* | 30.35 | 37.25 | 22.34 | 37.70 | 43.04 | 39.03 | -0.41 | 0.136 |
| *Lichenibacterium* | 19.08 | 21.70 | 28.22 | 20.10 | 33.17 | 39.03 | -0.42 | 0.303 |
| *Tamaricihabitans* | 34.47 | 52.97 | 51.07 | 45.49 | 33.68 | 39.03 | 0.23 | 0.387 |
| *Meira* | 0.65 | 0.51 | 0.50 | 2.34 | 16.96 | 38.77 | -5.12 | 0.218 |
| *Dokdonella* | 12.58 | 16.40 | 15.29 | 12.46 | 51.91 | 38.77 | -1.22 | 0.232 |
| *Tessaracoccus* | 28.62 | 26.31 | 28.56 | 21.34 | 27.85 | 38.77 | -0.07 | 0.800 |
| *Terrihabitans* | 7.37 | 3.93 | 10.25 | 5.45 | 18.99 | 38.51 | -1.55 | 0.285 |
| *Novipirellula* | 14.96 | 25.29 | 69.88 | 49.23 | 49.12 | 38.51 | -0.31 | 0.652 |
| *Kineococcus* | 29.05 | 32.29 | 30.40 | 23.06 | 27.35 | 38.25 | 0.05 | 0.838 |
| *Blastochloris* | 23.20 | 14.69 | 27.88 | 10.13 | 40.26 | 37.99 | -0.42 | 0.530 |
| *Symbiodinium* | 36.42 | 26.48 | 27.72 | 23.52 | 32.66 | 37.99 | -0.05 | 0.837 |
| *Intrasporangium* | 34.26 | 27.51 | 35.78 | 25.24 | 23.80 | 37.73 | 0.17 | 0.526 |
| *Isoptericola* | 23.85 | 31.44 | 42.67 | 36.92 | 30.13 | 37.73 | -0.10 | 0.731 |
| *Umezawaea* | 27.54 | 39.13 | 34.27 | 33.18 | 31.40 | 37.73 | -0.02 | 0.915 |
| *Janibacter* | 34.47 | 30.92 | 56.44 | 21.34 | 33.42 | 37.47 | 0.40 | 0.360 |
| *Aquabacterium* | 18.86 | 14.35 | 25.03 | 12.31 | 18.99 | 37.47 | -0.24 | 0.700 |
| *Rhodoferax* | 87.38 | 25.63 | 51.40 | 24.46 | 30.38 | 37.21 | 0.84 | 0.309 |
| *Nordella* | 46.40 | 27.85 | 48.38 | 17.14 | 36.46 | 37.21 | 0.43 | 0.314 |
| *Alienimonas* | 8.89 | 19.82 | 37.80 | 30.53 | 25.83 | 37.21 | -0.49 | 0.402 |
| *Streptoalloteichus* | 32.31 | 39.64 | 35.28 | 33.03 | 31.65 | 37.21 | 0.07 | 0.543 |
| *Bremerella* | 11.92 | 23.07 | 53.75 | 39.10 | 33.68 | 37.21 | -0.31 | 0.629 |
| *Dissophora* | 1.95 | 1.88 | 1.51 | 2.34 | 6.84 | 36.95 | -3.11 | 0.337 |
| *Humisphaera* | 10.62 | 14.18 | 39.14 | 30.38 | 40.77 | 36.69 | -0.75 | 0.239 |
| *Sinosporangium* | 19.30 | 61.85 | 31.41 | 34.43 | 24.81 | 36.69 | 0.23 | 0.709 |
| *Pirellulimonas* | 4.12 | 10.93 | 20.16 | 18.54 | 30.89 | 36.43 | -1.29 | 0.075 |
| *Ginsengibacter* | 3.25 | 0.85 | 1.18 | 1.56 | 47.60 | 36.43 | -4.02 | 0.193 |
| *Comamonas* | 76.75 | 29.39 | 43.51 | 31.94 | 31.90 | 36.43 | 0.58 | 0.362 |
| *Azohydromonas* | 11.06 | 7.52 | 13.10 | 7.95 | 30.13 | 36.17 | -1.23 | 0.237 |
| *Luteimicrobium* | 8.02 | 9.91 | 18.65 | 13.87 | 15.95 | 35.65 | -0.84 | 0.302 |
| *Tumebacillus* | 28.62 | 38.61 | 45.69 | 25.39 | 32.41 | 35.65 | 0.27 | 0.335 |
| *Streblomastix* | 28.40 | 34.17 | 12.09 | 56.08 | 49.37 | 35.13 | -0.91 | 0.073 |
| *Chloropicon* | 18.65 | 36.56 | 17.81 | 42.22 | 36.71 | 35.13 | -0.64 | 0.144 |
| *Inquilinus* | 21.03 | 18.11 | 36.28 | 18.69 | 36.21 | 35.13 | -0.25 | 0.576 |
| *Litorilinea* | 10.41 | 7.86 | 15.12 | 4.52 | 22.79 | 34.87 | -0.90 | 0.391 |
| *Hydrogenophaga* | 31.00 | 27.00 | 54.59 | 24.61 | 33.42 | 34.87 | 0.28 | 0.533 |
| *Thermocrispum* | 22.77 | 30.92 | 31.08 | 27.73 | 28.61 | 34.87 | -0.10 | 0.579 |
| *Krasilnikoviella* | 14.53 | 12.81 | 12.09 | 19.32 | 17.22 | 34.61 | -0.85 | 0.191 |
| *Arenimonas* | 12.14 | 8.20 | 12.93 | 6.54 | 33.68 | 34.61 | -1.17 | 0.270 |
| *Actinotalea* | 19.73 | 17.43 | 23.85 | 16.83 | 24.05 | 34.61 | -0.31 | 0.455 |
| *Syntrophaceticus* | 0.87 | 1.71 | 5.21 | 1.56 | 30.13 | 34.35 | -3.08 | 0.198 |
| *Planobispora* | 20.38 | 61.68 | 31.24 | 37.55 | 22.79 | 34.35 | 0.26 | 0.674 |
| *Desulfosporosinus* | 11.27 | 18.96 | 20.16 | 14.96 | 28.87 | 34.09 | -0.63 | 0.247 |
| *Ostreococcus* | 23.85 | 38.10 | 18.31 | 36.30 | 36.21 | 34.09 | -0.41 | 0.275 |
| *Jiella* | 36.64 | 49.04 | 64.67 | 48.45 | 31.65 | 34.09 | 0.40 | 0.290 |
| *Sphaerobacter* | 11.27 | 5.13 | 7.06 | 3.12 | 21.78 | 34.09 | -1.33 | 0.318 |
| *Tsukamurella* | 17.56 | 20.16 | 24.19 | 16.05 | 35.95 | 34.09 | -0.47 | 0.332 |
| *Glomus* | 17.13 | 33.49 | 12.93 | 50.63 | 40.01 | 33.82 | -0.97 | 0.067 |
| *Soehngenia* | 16.69 | 31.27 | 19.65 | 32.87 | 34.18 | 33.82 | -0.58 | 0.129 |
| *Acidocella* | 14.09 | 20.33 | 24.52 | 15.73 | 36.21 | 33.82 | -0.54 | 0.303 |
| *Acidisoma* | 16.04 | 21.70 | 27.04 | 17.29 | 41.27 | 33.82 | -0.51 | 0.327 |
| *Enhygromyxa* | 6.29 | 5.13 | 10.25 | 3.27 | 29.88 | 33.56 | -1.62 | 0.254 |
| *Bartonella* | 28.19 | 23.41 | 33.09 | 23.37 | 22.03 | 33.56 | 0.10 | 0.697 |
| *Geminicoccus* | 22.98 | 28.87 | 33.43 | 33.49 | 30.89 | 33.30 | -0.20 | 0.302 |
| *Pycnococcus* | 23.63 | 38.78 | 21.17 | 45.96 | 42.54 | 33.04 | -0.54 | 0.142 |
| *Pyxidicoccus* | 13.88 | 7.52 | 11.59 | 7.63 | 37.22 | 33.04 | -1.24 | 0.244 |
| *Adhaeretor* | 5.64 | 10.42 | 20.16 | 11.53 | 22.03 | 33.04 | -0.88 | 0.258 |
| *Skermanella* | 22.11 | 17.60 | 28.56 | 19.01 | 35.70 | 33.04 | -0.36 | 0.356 |
| *Oligoflexus* | 4.77 | 20.33 | 12.26 | 25.86 | 44.31 | 32.78 | -1.46 | 0.037 |
| *Cryptococcus* | 15.18 | 29.05 | 13.61 | 50.79 | 35.20 | 32.52 | -1.03 | 0.057 |
| *Paludibaculum* | 4.77 | 3.59 | 7.22 | 5.76 | 35.95 | 32.26 | -2.25 | 0.176 |
| *Acuticoccus* | 16.48 | 21.53 | 30.40 | 17.14 | 22.28 | 32.00 | -0.06 | 0.876 |
| *Desulfovibrio* | 17.78 | 18.45 | 20.83 | 20.72 | 25.83 | 31.74 | -0.46 | 0.147 |
| *Pararhizobium* | 42.93 | 53.14 | 83.65 | 44.87 | 31.14 | 31.74 | 0.74 | 0.179 |
| *Hyalangium* | 9.54 | 9.57 | 8.90 | 6.08 | 34.94 | 31.74 | -1.38 | 0.244 |
| *Alcanivorax* | 12.14 | 15.89 | 15.79 | 13.87 | 21.78 | 31.74 | -0.62 | 0.265 |
| *Golovinomyces* | 11.92 | 13.33 | 14.28 | 30.38 | 20.76 | 31.48 | -1.06 | 0.047 |
| *Aquabacter* | 18.21 | 20.16 | 34.27 | 16.20 | 31.14 | 31.48 | -0.12 | 0.788 |
| *Salinibacterium* | 26.67 | 22.38 | 53.42 | 40.66 | 26.08 | 31.48 | 0.06 | 0.903 |

| | | | | | | | | |
|---|---|---|---|---|---|---|---|---|
| *Planctomonas* | 50.52 | 61.85 | 82.31 | 52.35 | 17.47 | 31.22 | 0.95 | 0.086 |
| *Stella* | 19.51 | 13.50 | 24.69 | 16.67 | 25.57 | 31.22 | -0.35 | 0.383 |
| *Cellulosimicrobium* | 16.48 | 17.26 | 30.24 | 15.58 | 37.22 | 31.22 | -0.39 | 0.448 |
| *Nesterenkonia* | 14.09 | 24.09 | 26.71 | 20.41 | 23.29 | 31.22 | -0.21 | 0.543 |
| *Roseiconus* | 8.02 | 24.60 | 61.82 | 48.92 | 35.20 | 31.22 | -0.29 | 0.711 |
| *Aldersonia* | 2.82 | 7.18 | 7.06 | 4.05 | 24.05 | 30.96 | -1.79 | 0.222 |
| *Photobacterium* | 18.86 | 21.87 | 21.17 | 32.25 | 26.33 | 30.70 | -0.53 | 0.020 |
| *Actinosynnema* | 20.16 | 25.97 | 22.17 | 21.03 | 22.54 | 30.70 | -0.12 | 0.604 |
| *Streptococcus* | 27.10 | 50.57 | 47.87 | 35.68 | 21.52 | 30.44 | 0.52 | 0.229 |
| *Rhodoligotrophos* | 20.16 | 20.33 | 38.13 | 17.76 | 25.32 | 30.44 | 0.10 | 0.820 |
| *Oricola* | 30.79 | 41.52 | 55.94 | 31.63 | 26.59 | 30.18 | 0.54 | 0.205 |
| *Rathayibacter* | 59.62 | 50.40 | 118.76 | 88.80 | 20.51 | 30.18 | 0.71 | 0.381 |
| *Salinarimonas* | 18.86 | 18.96 | 32.42 | 14.33 | 29.62 | 30.18 | -0.08 | 0.861 |
| *Oryzicola* | 27.10 | 13.84 | 22.85 | 11.68 | 22.54 | 29.92 | -0.01 | 0.989 |
| *Citrobacter* | 26.23 | 25.63 | 16.13 | 31.16 | 29.88 | 29.66 | -0.41 | 0.145 |
| *Luteitalea* | 10.19 | 6.15 | 14.45 | 10.91 | 29.88 | 29.66 | -1.19 | 0.159 |
| *Jongsikchunia* | 3.04 | 2.05 | 5.71 | 3.27 | 20.76 | 29.66 | -2.31 | 0.204 |
| *Candidatus Berkiella* | 20.81 | 3.59 | 3.19 | 1.09 | 13.17 | 29.66 | -0.67 | 0.624 |
| *Calycomorphotria* | 11.71 | 16.91 | 49.89 | 31.00 | 28.36 | 29.66 | -0.18 | 0.797 |
| *Humibacter* | 41.84 | 84.57 | 115.74 | 67.15 | 18.74 | 29.14 | 1.07 | 0.187 |
| *Pusillimonas* | 41.41 | 54.67 | 99.44 | 66.05 | 31.65 | 29.14 | 0.62 | 0.349 |
| *Oidiodendron* | 3.90 | 2.73 | 0.34 | 2.49 | 26.08 | 28.88 | -3.04 | 0.180 |
| *Gemmobacter* | 26.88 | 42.37 | 59.30 | 28.35 | 25.32 | 28.88 | 0.64 | 0.241 |
| *Ferrovibrio* | 8.02 | 9.23 | 17.30 | 10.75 | 18.99 | 28.88 | -0.76 | 0.269 |
| *Isosphaera* | 10.41 | 14.52 | 47.87 | 9.19 | 23.80 | 28.88 | 0.24 | 0.801 |
| *Ostreobium* | 14.31 | 29.39 | 15.12 | 37.23 | 36.46 | 28.62 | -0.80 | 0.078 |
| *Dictyobacter* | 14.53 | 13.67 | 16.46 | 13.55 | 25.57 | 28.62 | -0.60 | 0.233 |
| *Roseovarius* | 21.46 | 29.39 | 39.31 | 18.85 | 23.29 | 28.62 | 0.35 | 0.348 |
| *Pectobacterium* | 23.20 | 37.25 | 16.13 | 40.04 | 31.14 | 28.62 | -0.38 | 0.354 |
| *Qaidamihabitans* | 25.37 | 35.54 | 29.23 | 29.29 | 23.29 | 28.62 | 0.15 | 0.451 |
| *Knoellia* | 35.34 | 26.65 | 43.67 | 19.79 | 25.32 | 28.36 | 0.53 | 0.147 |
| *Longispora* | 12.36 | 21.19 | 14.95 | 16.98 | 19.24 | 28.36 | -0.41 | 0.290 |
| *Thermocatellispora* | 18.65 | 46.81 | 26.88 | 29.60 | 21.27 | 28.36 | 0.22 | 0.659 |
| *Enterovirga* | 18.43 | 42.54 | 57.62 | 27.57 | 24.56 | 28.10 | 0.56 | 0.378 |
| *Undibacterium* | 39.03 | 16.23 | 30.07 | 22.90 | 18.48 | 28.10 | 0.30 | 0.521 |
| *Pelagicola* | 20.16 | 50.23 | 20.33 | 42.69 | 35.95 | 28.10 | -0.23 | 0.661 |
| *Thermostaphylospora* | 16.48 | 55.70 | 22.85 | 28.82 | 19.75 | 28.10 | 0.31 | 0.668 |
| *Candidatus* Entotheonella | 8.67 | 8.03 | 13.44 | 8.26 | 29.12 | 27.84 | -1.11 | 0.221 |
| *Subtercola* | 48.57 | 38.10 | 101.29 | 65.90 | 16.71 | 27.84 | 0.77 | 0.356 |
| *Spongiactinospora* | 16.48 | 55.53 | 25.53 | 29.60 | 21.52 | 27.84 | 0.31 | 0.655 |
| *Ruania* | 16.91 | 19.14 | 34.10 | 22.28 | 18.74 | 27.84 | 0.03 | 0.946 |
| *Rhizocola* | 12.58 | 13.84 | 12.60 | 10.59 | 18.23 | 27.58 | -0.53 | 0.359 |
| *Saitoella* | 14.96 | 17.94 | 9.74 | 31.63 | 28.61 | 27.32 | -1.04 | 0.011 |
| *Ruegeria* | 22.98 | 25.80 | 37.96 | 24.93 | 23.55 | 27.32 | 0.20 | 0.511 |
| *Thermobispora* | 16.48 | 43.06 | 28.56 | 27.11 | 21.78 | 27.32 | 0.21 | 0.660 |
| *Pigmentiphaga* | 24.93 | 16.23 | 24.86 | 16.20 | 25.07 | 27.32 | -0.05 | 0.860 |
| *Alkalihalobacillus* | 35.34 | 229.80 | 115.74 | 52.50 | 21.78 | 27.06 | 1.91 | 0.238 |
| *Antricoccus* | 11.06 | 32.46 | 17.30 | 17.92 | 20.76 | 27.06 | -0.11 | 0.830 |
| *Mammaliicoccus* | 16.69 | 26.14 | 11.93 | 40.35 | 30.89 | 26.80 | -0.84 | 0.068 |
| *Janthinobacterium* | 39.24 | 27.34 | 38.47 | 21.50 | 23.55 | 26.80 | 0.55 | 0.087 |
| *Fimbriimonas* | 12.58 | 2.39 | 5.38 | 3.89 | 30.13 | 26.80 | -1.58 | 0.238 |
| *Geobacter* | 10.41 | 12.99 | 16.63 | 11.53 | 24.56 | 26.80 | -0.65 | 0.246 |
| *Lichenihabitans* | 9.76 | 12.47 | 17.30 | 10.13 | 20.76 | 26.80 | -0.54 | 0.345 |
| *Agaricicola* | 5.42 | 8.71 | 16.13 | 7.01 | 16.46 | 26.80 | -0.73 | 0.380 |
| *Murinocardiopsis* | 17.13 | 59.80 | 32.59 | 29.60 | 20.76 | 26.80 | 0.51 | 0.480 |
| *Lacunisphaera* | 8.24 | 2.39 | 2.69 | 3.89 | 18.99 | 26.54 | -1.89 | 0.206 |
| *Herbihabitans* | 21.03 | 29.05 | 24.36 | 24.77 | 28.87 | 26.54 | -0.11 | 0.518 |
| *Pontibacter* | 18.86 | 28.70 | 29.73 | 18.54 | 29.62 | 26.54 | 0.05 | 0.863 |
| *Xenophilus* | 14.31 | 8.03 | 13.94 | 23.99 | 59.76 | 26.28 | -1.60 | 0.164 |
| *Desulfuromonas* | 7.37 | 11.11 | 13.77 | 8.88 | 29.12 | 26.28 | -0.99 | 0.228 |
| *Armatimonas* | 8.89 | 8.20 | 23.01 | 16.83 | 32.92 | 26.02 | -0.92 | 0.151 |
| *Hondaea* | 47.48 | 34.34 | 30.91 | 29.29 | 25.83 | 26.02 | 0.48 | 0.168 |
| *Methyloferula* | 8.89 | 7.18 | 13.94 | 9.97 | 25.57 | 26.02 | -1.04 | 0.174 |
| *Phytophthora* | 51.60 | 22.72 | 18.98 | 26.95 | 22.54 | 26.02 | 0.31 | 0.623 |
| *Rhizobacter* | 15.83 | 12.99 | 27.55 | 12.93 | 23.80 | 26.02 | -0.15 | 0.742 |
| *Ideonella* | 11.71 | 11.11 | 23.52 | 9.35 | 14.18 | 26.02 | -0.10 | 0.876 |
| *Sphingobacterium* | 16.26 | 18.11 | 25.20 | 29.44 | 30.13 | 25.76 | -0.52 | 0.068 |
| *Propylenella* | 12.14 | 8.71 | 22.01 | 10.59 | 20.51 | 25.76 | -0.41 | 0.479 |
| *Stackebrandtia* | 15.39 | 25.63 | 17.47 | 18.07 | 20.51 | 25.76 | -0.14 | 0.644 |
| *Amnibacterium* | 64.83 | 8.88 | 28.56 | 19.01 | 20.76 | 25.50 | 0.65 | 0.531 |
| *Saitozyma* | 0.22 | 0.17 | 0.34 | 0.16 | 35.95 | 25.24 | -6.40 | 0.197 |
| *Spirosoma* | 27.10 | 222.97 | 2405.80 | 394.46 | 30.89 | 25.24 | 2.56 | 0.437 |
| *Blastomonas* | 7.37 | 15.04 | 26.20 | 15.89 | 15.70 | 25.24 | -0.22 | 0.692 |
| *Dietzia* | 11.71 | 18.45 | 17.30 | 14.80 | 22.03 | 24.98 | -0.38 | 0.270 |
| *Pseudoalteromonas* | 25.15 | 20.50 | 21.33 | 26.17 | 20.26 | 24.98 | -0.09 | 0.564 |
| *Aurantiacibacter* | 16.04 | 35.88 | 44.18 | 36.45 | 21.27 | 24.98 | 0.22 | 0.670 |
| *Sinomonas* | 36.64 | 42.37 | 60.98 | 40.51 | 28.36 | 24.72 | 0.58 | 0.163 |
| *Acanthamoeba* | 26.45 | 217.50 | 31.24 | 30.69 | 25.57 | 24.72 | 1.77 | 0.412 |
| *Pararhodospirillum* | 19.73 | 22.72 | 30.74 | 19.63 | 21.52 | 24.72 | 0.15 | 0.547 |
| *Marinitenerispora* | 16.91 | 58.95 | 27.04 | 33.65 | 22.79 | 24.72 | 0.34 | 0.629 |
| *Tomitella* | 9.76 | 22.38 | 29.40 | 22.90 | 20.76 | 24.72 | -0.15 | 0.732 |
| *Methylorubrum* | 15.39 | 23.07 | 28.89 | 17.92 | 22.03 | 24.72 | 0.06 | 0.849 |
| *Elioraea* | 10.62 | 11.96 | 14.61 | 9.50 | 24.56 | 24.46 | -0.65 | 0.289 |
| *Marinactinospora* | 15.83 | 54.16 | 24.02 | 27.57 | 20.76 | 24.46 | 0.37 | 0.607 |
| *Volvox* | 17.35 | 16.06 | 12.09 | 16.83 | 26.08 | 24.20 | -0.56 | 0.108 |
| *Dongia* | 12.14 | 3.08 | 8.06 | 4.83 | 18.99 | 24.20 | -1.04 | 0.291 |
| *Rubellimicrobium* | 18.21 | 29.73 | 41.49 | 21.19 | 25.83 | 24.20 | 0.33 | 0.462 |
| *Salipiger* | 15.61 | 19.48 | 16.46 | 21.50 | 22.28 | 23.94 | -0.39 | 0.025 |
| *Microbulbifer* | 14.74 | 9.06 | 16.46 | 12.77 | 20.51 | 23.94 | -0.51 | 0.238 |
| *Planoprotostelium* | 28.19 | 76.71 | 41.66 | 64.96 | 28.11 | 23.94 | 0.32 | 0.640 |
| *Qipengyuania* | 30.35 | 44.25 | 53.42 | 34.43 | 32.16 | 23.68 | 0.50 | 0.192 |
| *Vulgatibacter* | 4.55 | 2.56 | 3.19 | 2.34 | 20.76 | 23.68 | -2.18 | 0.210 |
| *Nitrolancea* | 4.55 | 2.73 | 5.88 | 1.40 | 16.96 | 23.42 | -1.66 | 0.281 |
| *Pseudarthrobacter* | 27.10 | 24.94 | 40.15 | 31.16 | 16.71 | 23.42 | 0.37 | 0.333 |
| *Afifella* | 14.53 | 13.67 | 21.17 | 11.84 | 23.55 | 23.42 | -0.25 | 0.534 |
| *Tianweitania* | 20.38 | 15.38 | 22.01 | 18.07 | 15.45 | 23.42 | 0.02 | 0.930 |
| *Herminiimonas* | 13.01 | 24.77 | 42.83 | 16.20 | 15.95 | 23.16 | 0.54 | 0.435 |
| *Thioclava* | 16.48 | 16.23 | 25.87 | 11.37 | 16.46 | 23.16 | 0.20 | 0.613 |
| *Acidipila* | 3.69 | 2.73 | 5.21 | 3.74 | 26.84 | 22.90 | -2.20 | 0.189 |
| *Thermoleophilum* | 8.02 | 1.20 | 2.69 | 0.62 | 18.48 | 22.90 | -1.82 | 0.276 |

| | | | | | | | | |
|---|---|---|---|---|---|---|---|---|
| *Neobacillus* | 40.11 | 222.11 | 195.19 | 125.72 | 21.52 | 22.64 | 1.43 | 0.237 |
| *Pelomonas* | 35.12 | 26.31 | 37.12 | 41.75 | 21.52 | 22.64 | 0.20 | 0.608 |
| *Schaalia* | 13.88 | 24.43 | 17.81 | 16.51 | 11.90 | 22.64 | 0.14 | 0.717 |
| *Solirhodobacter* | 7.59 | 10.25 | 25.03 | 6.70 | 18.99 | 22.64 | -0.17 | 0.815 |
| *Humibacillus* | 19.51 | 12.30 | 15.12 | 9.04 | 17.22 | 22.64 | -0.06 | 0.896 |
| *Alteraurantiacibacter* | 16.26 | 27.00 | 30.40 | 29.76 | 23.29 | 22.64 | -0.04 | 0.898 |
| *Pannonibacter* | 24.07 | 34.34 | 47.37 | 26.02 | 16.71 | 22.38 | 0.70 | 0.171 |
| *Chryseobacterium* | 15.39 | 29.05 | 39.98 | 26.95 | 24.56 | 22.38 | 0.19 | 0.672 |
| *Altericroceibacterium* | 10.19 | 26.14 | 31.75 | 23.99 | 17.98 | 22.38 | 0.08 | 0.867 |
| *Ectocarpus* | 31.22 | 14.69 | 8.90 | 10.91 | 23.29 | 22.38 | -0.04 | 0.947 |
| *Raphidocelis* | 12.14 | 16.06 | 10.58 | 20.25 | 20.51 | 22.12 | -0.70 | 0.028 |
| *Muricauda* | 16.91 | 20.33 | 14.78 | 31.78 | 31.14 | 22.12 | -0.71 | 0.053 |
| *Pseudobacteriovorax* | 1.73 | 2.90 | 2.35 | 4.52 | 21.02 | 22.12 | -2.77 | 0.140 |
| *Dunaliella* | 15.83 | 20.50 | 10.41 | 22.12 | 20.51 | 22.12 | -0.47 | 0.173 |
| *Protaetiibacter* | 6.29 | 8.71 | 19.15 | 12.15 | 19.24 | 22.12 | -0.65 | 0.266 |
| *Polynucleobacter* | 20.38 | 14.52 | 30.74 | 13.24 | 24.05 | 22.12 | 0.14 | 0.738 |
| *Defluviicoccus* | 6.29 | 4.27 | 11.09 | 4.99 | 16.71 | 21.86 | -1.01 | 0.279 |
| *Mycetocola* | 45.75 | 30.07 | 86.01 | 53.59 | 11.90 | 21.86 | 0.89 | 0.305 |
| *Mycoplana* | 13.66 | 33.49 | 46.03 | 25.86 | 15.19 | 21.86 | 0.57 | 0.399 |
| *Herbiconiux* | 43.80 | 35.20 | 107.51 | 70.42 | 16.46 | 21.86 | 0.78 | 0.419 |
| *Bifidobacterium* | 15.39 | 18.79 | 16.46 | 13.09 | 20.26 | 21.86 | -0.12 | 0.642 |
| *Candidatus* Accumulibacter | 12.36 | 7.52 | 10.08 | 11.37 | 21.02 | 21.60 | -0.85 | 0.122 |
| *Bryobacter* | 3.69 | 1.88 | 6.72 | 5.14 | 19.24 | 21.60 | -1.90 | 0.152 |
| *Brevifollis* | 4.55 | 6.66 | 17.13 | 12.15 | 20.26 | 21.60 | -0.93 | 0.160 |
| *Terriglobus* | 7.15 | 6.83 | 11.76 | 7.48 | 23.55 | 21.60 | -1.03 | 0.212 |
| *Sanguibacter* | 10.41 | 9.06 | 16.13 | 11.68 | 13.93 | 21.60 | -0.41 | 0.359 |
| *Lipingzhangella* | 14.96 | 38.44 | 21.17 | 21.50 | 14.69 | 21.60 | 0.37 | 0.515 |
| *Estrella* | 15.39 | 4.10 | 7.22 | 1.71 | 13.42 | 21.60 | -0.46 | 0.651 |
| *Salinispora* | 11.71 | 18.79 | 30.24 | 25.71 | 13.67 | 21.60 | -0.01 | 0.991 |
| *Aneurinibacillus* | 47.48 | 109.52 | 77.94 | 63.09 | 16.71 | 21.34 | 1.22 | 0.130 |
| *Bathycoccus* | 12.14 | 24.77 | 10.75 | 26.33 | 26.08 | 21.34 | -0.63 | 0.182 |
| *Thermoactinomyces* | 11.49 | 40.15 | 142.78 | 20.56 | 14.18 | 21.34 | 1.79 | 0.367 |
| *Actinocrinis* | 11.92 | 20.50 | 8.73 | 11.84 | 21.02 | 21.34 | -0.40 | 0.409 |
| *Thermasporomyces* | 11.06 | 13.84 | 11.59 | 10.59 | 13.93 | 21.34 | -0.33 | 0.431 |
| *Bacteroides* | 19.51 | 10.42 | 15.79 | 12.31 | 19.50 | 21.34 | -0.22 | 0.554 |
| *Sphingorhabdus* | 11.92 | 22.21 | 34.94 | 17.45 | 17.22 | 21.34 | 0.30 | 0.581 |
| *Camelimonas* | 11.92 | 14.52 | 15.12 | 7.48 | 16.46 | 21.34 | -0.12 | 0.794 |
| *Mariniblastus* | 3.90 | 7.86 | 16.63 | 16.20 | 15.45 | 21.08 | -0.89 | 0.151 |
| *Longimicrobium* | 4.55 | 8.54 | 11.93 | 8.72 | 23.04 | 21.08 | -1.08 | 0.163 |
| *Tistrella* | 6.07 | 7.01 | 13.61 | 7.95 | 10.89 | 21.08 | -0.58 | 0.406 |
| *Chryseolinea* | 192.53 | 16.57 | 15.79 | 6.54 | 16.96 | 21.08 | 2.34 | 0.414 |
| *Niallia* | 4.12 | 19.31 | 16.97 | 8.10 | 27.85 | 21.08 | -0.50 | 0.501 |
| *Nitrospirillum* | 13.01 | 17.94 | 18.65 | 14.49 | 20.00 | 21.08 | -0.16 | 0.503 |
| *Thioalkalivibrio* | 10.62 | 10.25 | 13.44 | 9.66 | 24.31 | 20.82 | -0.67 | 0.260 |
| *Spinactinospora* | 15.83 | 55.87 | 28.22 | 34.59 | 21.52 | 20.82 | 0.38 | 0.594 |
| *Promicromonospora* | 29.70 | 30.07 | 30.40 | 25.24 | 22.54 | 20.56 | 0.40 | 0.032 |
| *Hoyosella* | 10.41 | 12.13 | 11.09 | 11.37 | 19.50 | 20.56 | -0.61 | 0.173 |
| *Stigmatella* | 9.11 | 8.20 | 9.91 | 10.13 | 27.60 | 20.56 | -1.10 | 0.177 |
| *Tabrizicola* | 20.60 | 49.04 | 58.29 | 42.84 | 22.03 | 20.56 | 0.58 | 0.361 |
| *Moorella* | 2.17 | 8.20 | 12.93 | 2.65 | 23.55 | 20.56 | -1.00 | 0.362 |
| *Fluviicola* | 31.44 | 6.15 | 9.07 | 15.27 | 52.67 | 20.56 | -0.92 | 0.388 |
| *Thalassiosira* | 171.28 | 10.93 | 9.07 | 9.97 | 24.81 | 20.56 | 1.79 | 0.488 |
| *Allorhizocola* | 9.54 | 8.54 | 9.07 | 9.19 | 7.34 | 20.56 | -0.45 | 0.507 |
| *Helicosporidium* | 9.11 | 12.30 | 8.90 | 14.33 | 10.89 | 20.29 | -0.59 | 0.198 |
| *Virgibacillus* | 11.71 | 318.30 | 70.55 | 28.51 | 11.90 | 20.29 | 2.72 | 0.351 |
| *Aliidongia* | 12.79 | 10.25 | 16.63 | 7.32 | 27.35 | 20.29 | -0.47 | 0.482 |
| *Trichoderma* | 4.77 | 7.18 | 6.72 | 13.40 | 11.65 | 20.03 | -1.27 | 0.065 |
| *Nitriliruptor* | 6.72 | 12.99 | 9.07 | 10.91 | 14.43 | 20.03 | -0.66 | 0.170 |
| *Desulfotomaculum* | 3.47 | 4.95 | 5.54 | 2.03 | 19.75 | 20.03 | -1.58 | 0.259 |
| *Telmatospirillum* | 12.36 | 7.86 | 9.74 | 6.54 | 18.74 | 20.03 | -0.60 | 0.357 |
| *Malassezia* | 24.93 | 20.16 | 11.25 | 33.03 | 65.33 | 19.77 | -1.07 | 0.264 |
| *Acidimicrobium* | 3.90 | 3.76 | 3.53 | 2.96 | 16.96 | 19.51 | -1.82 | 0.209 |
| *Curvibacter* | 117.08 | 21.02 | 50.90 | 24.93 | 17.47 | 19.51 | 1.61 | 0.274 |
| *Ktedonobacter* | 7.59 | 8.03 | 9.07 | 5.45 | 16.71 | 19.51 | -0.75 | 0.318 |
| *Amaricoccus* | 11.27 | 10.76 | 16.80 | 10.28 | 11.39 | 19.51 | -0.08 | 0.836 |
| *Plasmodiophora* | 38.59 | 7.86 | 7.90 | 8.10 | 22.79 | 19.51 | 0.11 | 0.913 |
| *Colletotrichum* | 7.81 | 11.79 | 7.06 | 18.54 | 9.12 | 19.25 | -0.82 | 0.164 |
| *Nibricoccus* | 9.97 | 3.42 | 3.86 | 3.58 | 36.21 | 19.25 | -1.77 | 0.275 |
| *Sulfitobacter* | 15.61 | 25.29 | 30.07 | 17.76 | 18.23 | 19.25 | 0.36 | 0.341 |
| *Thiomonas* | 22.77 | 11.62 | 24.02 | 14.33 | 12.91 | 19.25 | 0.33 | 0.433 |
| *Spiractinospora* | 16.04 | 57.75 | 25.03 | 28.82 | 16.46 | 19.25 | 0.61 | 0.466 |
| *Quadrisphaera* | 10.62 | 17.43 | 18.14 | 14.02 | 15.95 | 19.25 | -0.09 | 0.744 |
| *Pseudovibrio* | 11.92 | 14.52 | 23.35 | 12.77 | 17.98 | 19.25 | -0.01 | 0.989 |
| *Bacteriovorax* | 6.50 | 5.30 | 5.21 | 3.89 | 11.65 | 18.99 | -1.02 | 0.312 |
| *Kroppenstedtia* | 8.02 | 29.39 | 66.69 | 12.93 | 13.42 | 18.99 | 1.20 | 0.371 |
| *Rhizorhapis* | 5.85 | 8.03 | 19.49 | 10.75 | 18.48 | 18.99 | -0.53 | 0.387 |
| *Leekyejoonella* | 10.84 | 13.33 | 17.81 | 10.13 | 16.46 | 18.99 | -0.12 | 0.739 |
| *Tistlia* | 11.92 | 10.59 | 14.28 | 6.23 | 15.45 | 18.99 | -0.14 | 0.773 |
| *Valsa* | 11.27 | 12.64 | 11.42 | 17.45 | 14.94 | 18.73 | -0.53 | 0.029 |
| *Pseudobythopirellula* | 3.25 | 7.18 | 11.42 | 10.59 | 16.46 | 18.73 | -1.07 | 0.077 |
| *Flectobacillus* | 11.49 | 15.55 | 10.58 | 30.07 | 18.23 | 18.73 | -0.83 | 0.114 |
| *Cumulibacter* | 14.31 | 23.24 | 14.28 | 14.49 | 16.21 | 18.73 | 0.07 | 0.820 |
| *Pseudooceanicola* | 11.49 | 12.81 | 21.67 | 12.77 | 16.21 | 18.73 | -0.05 | 0.884 |
| *Sneathiella* | 9.32 | 8.71 | 14.11 | 10.75 | 17.98 | 18.47 | -0.55 | 0.181 |
| *Ferruginibacter* | 5.20 | 2.90 | 5.38 | 1.56 | 26.08 | 18.47 | -1.77 | 0.272 |
| *Angustibacter* | 10.19 | 14.01 | 11.93 | 8.41 | 21.52 | 18.47 | -0.42 | 0.412 |
| *Luteipulveratus* | 10.41 | 11.62 | 11.42 | 8.41 | 10.13 | 18.47 | -0.14 | 0.741 |
| *Pseudochelatococcus* | 9.54 | 14.69 | 22.51 | 9.50 | 14.69 | 18.47 | 0.13 | 0.781 |
| *Cohaesibacter* | 19.30 | 23.75 | 31.08 | 16.98 | 13.42 | 18.21 | 0.61 | 0.116 |
| *Emericellopsis* | 1.30 | 4.61 | 0.67 | 27.42 | 4.30 | 18.21 | -2.92 | 0.161 |
| *Oceanobacillus* | 14.74 | 285.16 | 83.32 | 22.59 | 14.94 | 18.21 | 2.78 | 0.311 |
| *Kallotenue* | 11.71 | 5.81 | 8.73 | 6.85 | 14.43 | 18.21 | -0.59 | 0.324 |
| *Parvibaculum* | 16.48 | 10.25 | 12.60 | 9.66 | 24.05 | 18.21 | -0.40 | 0.432 |
| *Schumannella* | 20.81 | 21.02 | 56.27 | 40.51 | 13.67 | 18.21 | 0.44 | 0.588 |
| *Tenggerimyces* | 12.79 | 34.00 | 17.13 | 19.16 | 14.43 | 18.21 | 0.30 | 0.599 |
| *Microterricola* | 12.58 | 12.99 | 22.68 | 17.60 | 8.10 | 18.21 | 0.14 | 0.772 |
| *Aphanomyces* | 28.62 | 20.33 | 14.11 | 20.25 | 23.55 | 18.21 | 0.03 | 0.941 |
| *Pseudanabaena* | 6.72 | 7.01 | 10.58 | 8.72 | 13.67 | 17.95 | -0.73 | 0.171 |
| *Pantoea* | 4394.16 | 24.77 | 872.99 | 59.98 | 13.17 | 17.95 | 5.86 | 0.325 |
| *Klenkia* | 11.06 | 12.99 | 14.28 | 12.46 | 10.13 | 17.95 | -0.08 | 0.790 |

| | | | | | | | | |
|---|---|---|---|---|---|---|---|---|
| *Sulfobacillus* | 5.42 | 23.24 | 14.45 | 5.76 | 14.18 | 17.95 | 0.19 | 0.796 |
| *Alloactinosynnema* | 16.91 | 17.09 | 18.14 | 12.93 | 11.39 | 17.69 | 0.31 | 0.212 |
| *Thermogutta* | 3.90 | 5.30 | 16.29 | 9.35 | 19.75 | 17.69 | -0.88 | 0.235 |
| *Aeromonas* | 14.09 | 19.99 | 21.50 | 16.98 | 12.15 | 17.69 | 0.25 | 0.366 |
| *Zavarzinia* | 9.54 | 5.81 | 10.25 | 5.76 | 13.67 | 17.69 | -0.54 | 0.393 |
| *Chlamydia* | 11.27 | 16.06 | 19.49 | 19.79 | 13.42 | 17.69 | -0.12 | 0.679 |
| *Agrococcus* | 27.97 | 22.21 | 40.99 | 24.61 | 13.17 | 17.43 | 0.72 | 0.153 |
| *Criblamydia* | 4.99 | 4.78 | 9.57 | 4.99 | 15.45 | 17.43 | -0.97 | 0.247 |
| *Ammoniphilus* | 21.68 | 118.06 | 144.46 | 86.77 | 11.65 | 17.43 | 1.29 | 0.286 |
| *Oceanibaculum* | 7.81 | 7.18 | 10.08 | 7.17 | 12.41 | 17.43 | -0.56 | 0.310 |
| *Sporosarcina* | 9.76 | 137.03 | 47.87 | 26.95 | 14.94 | 17.43 | 1.71 | 0.354 |
| *Methylobrevis* | 9.11 | 10.93 | 15.79 | 7.79 | 15.70 | 17.43 | -0.19 | 0.664 |
| *Gorillibacterium* | 28.40 | 71.59 | 34.94 | 35.52 | 15.19 | 17.17 | 0.99 | 0.235 |
| *Hyphomonas* | 13.66 | 13.84 | 19.65 | 11.68 | 15.95 | 17.17 | 0.07 | 0.774 |
| *Collimonas* | 19.95 | 11.45 | 19.32 | 11.22 | 19.50 | 17.17 | 0.08 | 0.808 |
| *Arachidicoccus* | 31.44 | 3.08 | 5.21 | 2.03 | 15.19 | 17.17 | 0.21 | 0.872 |
| *Zhengella* | 15.61 | 18.62 | 24.52 | 10.91 | 6.08 | 16.91 | 0.79 | 0.114 |
| *Frigoribacterium* | 32.52 | 27.17 | 73.91 | 48.45 | 13.17 | 16.91 | 0.77 | 0.382 |
| *Thermoactinospora* | 15.18 | 34.68 | 17.47 | 18.54 | 13.17 | 16.91 | 0.47 | 0.419 |
| *Tepidamorphus* | 7.59 | 7.52 | 13.44 | 5.61 | 14.69 | 16.91 | -0.38 | 0.518 |
| *Allosalinactinospora* | 10.41 | 27.00 | 14.61 | 16.98 | 8.10 | 16.91 | 0.31 | 0.601 |
| *Acidithiobacillus* | 7.15 | 8.37 | 12.43 | 7.79 | 13.42 | 16.65 | -0.44 | 0.349 |
| *Methylomonas* | 11.49 | 12.13 | 14.95 | 11.53 | 14.43 | 16.65 | -0.14 | 0.505 |
| *Vallicoccus* | 7.59 | 17.94 | 10.92 | 11.84 | 15.19 | 16.65 | -0.26 | 0.529 |
| *Methyloligella* | 25.58 | 8.20 | 19.49 | 9.66 | 15.95 | 16.65 | 0.34 | 0.559 |
| *Algoriphagus* | 11.92 | 16.06 | 26.37 | 13.71 | 16.46 | 16.65 | 0.22 | 0.619 |
| *Azoarcus* | 13.88 | 11.62 | 18.31 | 11.22 | 15.19 | 16.65 | 0.03 | 0.925 |
| *Limnohabitans* | 14.74 | 14.86 | 21.67 | 14.18 | 13.93 | 16.39 | 0.21 | 0.430 |
| *Thermopolyspora* | 11.49 | 32.29 | 20.16 | 19.94 | 10.89 | 16.39 | 0.44 | 0.465 |
| *Gluconobacter* | 16.04 | 21.36 | 18.98 | 18.38 | 18.99 | 16.39 | 0.07 | 0.645 |
| *Siccirubricoccus* | 9.11 | 9.06 | 14.78 | 11.84 | 18.99 | 16.13 | -0.51 | 0.172 |
| *Nitrososphaera* | 10.19 | 1.71 | 6.55 | 1.56 | 16.71 | 16.13 | -0.90 | 0.410 |
| *Neochlamydia* | 3.69 | 8.54 | 9.91 | 2.65 | 17.98 | 16.13 | -0.73 | 0.427 |
| *Emticicia* | 184.51 | 9.40 | 15.79 | 4.36 | 29.62 | 16.13 | 2.07 | 0.452 |
| *Glaciihabitans* | 16.69 | 15.55 | 40.32 | 27.57 | 8.10 | 16.13 | 0.49 | 0.526 |
| *Chthonobacter* | 15.83 | 17.43 | 25.87 | 20.41 | 17.47 | 16.13 | 0.13 | 0.650 |
| *Coxiella* | 10.84 | 7.52 | 10.75 | 2.96 | 15.95 | 16.13 | -0.27 | 0.700 |
| *Rhodovastum* | 5.20 | 11.11 | 8.23 | 7.63 | 17.98 | 15.87 | -0.76 | 0.211 |
| *Smaragdicoccus* | 6.50 | 4.10 | 6.89 | 4.67 | 12.66 | 15.87 | -0.92 | 0.253 |
| *Pseudoclavibacter* | 15.18 | 16.91 | 33.93 | 21.50 | 13.42 | 15.87 | 0.38 | 0.495 |
| *Falsochrobactrum* | 11.71 | 11.96 | 18.14 | 13.24 | 14.69 | 15.87 | -0.07 | 0.791 |
| *Phragmitibacter* | 1.52 | 4.44 | 7.22 | 8.72 | 12.66 | 15.61 | -1.49 | 0.039 |
| *Heliomicrobium* | 8.02 | 18.79 | 14.61 | 9.35 | 9.37 | 15.61 | 0.27 | 0.567 |
| *Salinicoccus* | 9.11 | 21.87 | 11.76 | 14.96 | 15.70 | 15.61 | -0.11 | 0.792 |
| *Marinobacterium* | 10.41 | 9.40 | 13.10 | 4.21 | 14.43 | 15.61 | -0.06 | 0.917 |
| *Spirillospora* | 6.07 | 28.53 | 12.93 | 18.85 | 11.39 | 15.61 | 0.05 | 0.942 |
| *Allostreptomyces* | 9.32 | 22.55 | 11.93 | 17.29 | 9.87 | 15.61 | 0.03 | 0.945 |
| *Hartmannibacter* | 8.02 | 12.64 | 15.12 | 9.35 | 11.39 | 15.61 | -0.02 | 0.950 |
| *Geothrix* | 27.54 | 2.39 | 2.86 | 3.27 | 13.67 | 15.61 | 0.01 | 0.992 |
| *Monoraphidium* | 9.76 | 15.55 | 8.90 | 15.73 | 17.22 | 15.35 | -0.50 | 0.146 |
| *Falsiroseomonas* | 8.02 | 11.45 | 13.10 | 11.06 | 15.95 | 15.35 | -0.38 | 0.202 |
| *Breoghania* | 13.44 | 13.16 | 18.81 | 8.10 | 18.99 | 15.35 | 0.10 | 0.803 |
| *Segnochrobactrum* | 25.80 | 37.08 | 50.39 | 21.65 | 17.47 | 15.09 | 1.06 | 0.100 |
| *Marinomonas* | 9.54 | 9.06 | 13.10 | 7.79 | 10.38 | 15.09 | -0.07 | 0.848 |
| *Oxalicibacterium* | 6.50 | 16.40 | 34.10 | 17.14 | 24.05 | 15.09 | 0.02 | 0.979 |
| *Stakelama* | 25.80 | 28.19 | 32.92 | 19.01 | 28.36 | 14.83 | 0.48 | 0.164 |
| *Micavibrio* | 7.81 | 5.98 | 6.22 | 5.14 | 14.69 | 14.83 | -0.79 | 0.265 |
| *Crateriforma* | 5.20 | 16.91 | 32.59 | 21.65 | 15.45 | 14.83 | 0.07 | 0.920 |
| *Thauera* | 10.41 | 7.86 | 11.09 | 6.39 | 16.46 | 14.57 | -0.35 | 0.482 |
| *Pseudohoeflea* | 20.38 | 31.95 | 42.83 | 24.77 | 11.65 | 14.31 | 0.91 | 0.137 |
| *Planktothrix* | 7.15 | 6.15 | 6.89 | 7.63 | 12.15 | 14.31 | -0.75 | 0.139 |
| *Gracilibacillus* | 10.41 | 93.29 | 84.83 | 23.21 | 8.61 | 14.31 | 2.03 | 0.211 |
| *Thalassobaculum* | 7.15 | 7.86 | 12.93 | 8.57 | 13.93 | 14.31 | -0.40 | 0.319 |
| *Frondihabitans* | 27.32 | 24.94 | 58.96 | 39.88 | 15.45 | 14.31 | 0.68 | 0.375 |
| *Micrococcus* | 8.24 | 13.50 | 13.10 | 50.48 | 9.87 | 14.31 | -1.10 | 0.411 |
| *Larkinella* | 10.41 | 12.13 | 110.87 | 18.69 | 12.66 | 14.31 | 1.55 | 0.471 |
| *Tetrahymena* | 11.06 | 15.21 | 8.23 | 12.15 | 13.42 | 14.31 | -0.21 | 0.475 |
| *Allobranchiibius* | 7.15 | 8.20 | 8.40 | 4.83 | 10.38 | 14.31 | -0.31 | 0.559 |
| *Aliihoeflea* | 15.18 | 8.37 | 14.78 | 9.19 | 12.66 | 14.31 | 0.09 | 0.799 |
| *Paradevosia* | 1.95 | 32.12 | 4.20 | 1.71 | 16.96 | 14.31 | 0.22 | 0.880 |
| *Polymorphobacter* | 12.58 | 16.40 | 16.46 | 13.09 | 18.74 | 14.31 | -0.02 | 0.922 |
| *Scytonema* | 7.59 | 4.44 | 6.55 | 8.57 | 10.89 | 14.05 | -0.85 | 0.067 |
| *Fulvivirga* | 23.63 | 20.16 | 15.45 | 8.41 | 16.21 | 14.05 | 0.62 | 0.107 |
| *Gluconacetobacter* | 9.54 | 10.25 | 12.09 | 10.59 | 17.22 | 14.05 | -0.39 | 0.217 |
| *Rudaea* | 5.20 | 8.54 | 7.56 | 5.45 | 18.74 | 14.05 | -0.84 | 0.281 |
| *Methylobacter* | 8.46 | 7.18 | 12.77 | 8.57 | 15.95 | 14.05 | -0.44 | 0.294 |
| *Treponema* | 5.42 | 7.01 | 8.73 | 4.99 | 9.87 | 14.05 | -0.45 | 0.434 |
| *Pilimelia* | 3.04 | 10.42 | 8.06 | 7.63 | 7.34 | 14.05 | -0.43 | 0.463 |
| *Segetibacter* | 12.58 | 11.79 | 17.47 | 14.80 | 16.71 | 14.05 | -0.12 | 0.573 |
| *Pseudoxanthobacter* | 10.62 | 13.16 | 22.68 | 12.62 | 12.91 | 14.05 | 0.23 | 0.596 |
| *Cucumibacter* | 7.37 | 8.03 | 10.58 | 4.67 | 6.33 | 14.05 | 0.05 | 0.925 |
| *Advenella* | 8.46 | 7.18 | 15.12 | 6.08 | 10.13 | 14.05 | 0.02 | 0.962 |
| *Calothrix* | 4.77 | 7.69 | 8.73 | 8.10 | 14.18 | 13.79 | -0.77 | 0.111 |
| *Peribacillus* | 18.65 | 72.96 | 78.28 | 30.69 | 15.19 | 13.79 | 1.51 | 0.187 |
| *Thermaerobacter* | 2.82 | 8.71 | 41.32 | 5.76 | 7.60 | 13.79 | 0.96 | 0.551 |
| *Caenibius* | 7.37 | 16.91 | 19.82 | 11.68 | 16.21 | 13.79 | 0.08 | 0.853 |
| *Mitsuaria* | 8.67 | 5.13 | 5.54 | 4.99 | 14.43 | 13.53 | -0.77 | 0.268 |
| *Belnapia* | 10.19 | 10.08 | 12.77 | 10.13 | 18.23 | 13.53 | -0.34 | 0.338 |
| *Paeniglutamicibacter* | 8.24 | 9.91 | 11.09 | 9.81 | 10.38 | 13.53 | -0.21 | 0.356 |
| *Cereibacter* | 12.58 | 13.50 | 27.38 | 12.77 | 14.69 | 13.53 | 0.38 | 0.476 |
| *Flavihumibacter* | 13.01 | 6.83 | 12.77 | 6.70 | 14.18 | 13.53 | -0.08 | 0.860 |
| *Caenimonas* | 31.44 | 37.76 | 67.19 | 34.43 | 20.26 | 13.27 | 1.00 | 0.164 |
| *Euzebya* | 6.29 | 7.52 | 10.75 | 8.26 | 12.91 | 13.27 | -0.49 | 0.192 |
| *Haematococcus* | 6.50 | 12.99 | 5.38 | 11.22 | 13.42 | 13.27 | -0.61 | 0.202 |
| *Sphingosinithalassobacter* | 16.04 | 15.89 | 18.81 | 15.42 | 16.46 | 13.27 | 0.17 | 0.233 |
| *Methylocaldum* | 4.77 | 5.81 | 9.57 | 4.21 | 12.91 | 13.27 | -0.59 | 0.380 |
| *Haloactinospora* | 10.62 | 30.58 | 14.95 | 16.83 | 10.89 | 13.27 | 0.45 | 0.496 |
| *Schlegelella* | 5.42 | 8.37 | 10.92 | 7.48 | 9.12 | 13.27 | -0.27 | 0.504 |
| *Liberibacter* | 9.97 | 15.38 | 24.86 | 13.87 | 14.43 | 13.27 | 0.27 | 0.575 |
| *Egicoccus* | 5.64 | 24.09 | 12.60 | 14.64 | 10.63 | 13.27 | 0.14 | 0.838 |

| | | | | | | | | |
|---|---|---|---|---|---|---|---|---|
| *Dickeya* | 8.46 | 12.64 | 10.25 | 14.33 | 10.89 | 13.01 | -0.29 | 0.222 |
| *Roseococcus* | 8.67 | 8.54 | 9.91 | 7.32 | 15.95 | 13.01 | -0.42 | 0.351 |
| *Maritimibacter* | 8.02 | 8.54 | 13.94 | 5.14 | 8.36 | 13.01 | 0.20 | 0.675 |
| *Corticibacterium* | 9.11 | 8.20 | 13.44 | 6.85 | 8.36 | 13.01 | 0.12 | 0.747 |
| *Sediminibacterium* | 50.95 | 11.45 | 28.22 | 14.02 | 8.36 | 12.75 | 1.37 | 0.246 |
| *Propionibacterium* | 9.97 | 9.57 | 9.91 | 9.19 | 11.90 | 12.75 | -0.20 | 0.304 |
| *Sphaerimonospora* | 12.14 | 25.12 | 12.77 | 10.44 | 10.89 | 12.75 | 0.55 | 0.334 |
| *Pseudactinotalea* | 8.89 | 8.20 | 9.91 | 7.63 | 9.87 | 12.75 | -0.16 | 0.547 |
| *Mangrovihabitans* | 5.20 | 15.04 | 11.42 | 11.84 | 6.58 | 12.75 | 0.02 | 0.964 |
| *Metabacillus* | 16.91 | 71.42 | 53.42 | 25.39 | 9.87 | 12.49 | 1.57 | 0.182 |
| *Thermobacillus* | 11.71 | 45.62 | 73.07 | 24.61 | 8.61 | 12.49 | 1.51 | 0.249 |
| *Phytomonospora* | 6.72 | 13.16 | 7.73 | 9.35 | 10.63 | 12.49 | -0.23 | 0.518 |
| *Acidimangrovimonas* | 8.46 | 9.74 | 15.96 | 3.27 | 10.89 | 12.49 | 0.36 | 0.532 |
| *Methylococcus* | 7.81 | 6.66 | 11.93 | 7.95 | 9.87 | 12.49 | -0.20 | 0.564 |
| *Chromobacterium* | 14.09 | 6.15 | 9.91 | 7.32 | 10.89 | 12.49 | -0.02 | 0.954 |
| *Paracraurococcus* | 11.06 | 8.37 | 9.74 | 13.87 | 13.67 | 12.23 | -0.45 | 0.025 |
| *Reticulibacter* | 9.76 | 4.95 | 6.22 | 5.61 | 11.39 | 12.23 | -0.48 | 0.343 |
| *Desulforamulus* | 2.60 | 5.64 | 14.11 | 4.83 | 12.41 | 12.23 | -0.40 | 0.610 |
| *Sporomusa* | 4.77 | 9.74 | 12.26 | 5.61 | 12.41 | 12.23 | -0.18 | 0.732 |
| *Synchytrium* | 16.26 | 11.11 | 5.71 | 9.35 | 12.41 | 12.23 | -0.04 | 0.935 |
| *Oryzihumus* | 10.84 | 12.47 | 15.12 | 9.35 | 16.46 | 12.23 | 0.02 | 0.957 |
| *Pseudogymnoascus* | 8.89 | 5.30 | 5.71 | 8.10 | 10.63 | 11.97 | -0.62 | 0.088 |
| *Lysinibacillus* | 14.09 | 73.81 | 44.51 | 21.50 | 13.17 | 11.97 | 1.51 | 0.237 |
| *Lacisediminihabitans* | 24.50 | 20.33 | 75.42 | 47.67 | 7.85 | 11.97 | 0.83 | 0.469 |
| *Austwickia* | 5.20 | 9.74 | 9.57 | 6.23 | 11.39 | 11.97 | -0.27 | 0.513 |
| *Adhaeribacter* | 8.46 | 21.36 | 21.84 | 11.37 | 20.51 | 11.97 | 0.24 | 0.651 |
| *Effusibacillus* | 19.73 | 38.61 | 37.12 | 28.98 | 11.90 | 11.71 | 0.86 | 0.162 |
| *Methanosarcina* | 3.25 | 4.10 | 6.89 | 2.65 | 29.37 | 11.71 | -1.62 | 0.337 |
| *Geomonas* | 4.99 | 5.30 | 10.75 | 5.61 | 12.41 | 11.71 | -0.50 | 0.369 |
| *Ottowia* | 13.88 | 10.42 | 17.13 | 9.81 | 13.17 | 11.71 | 0.26 | 0.375 |
| *Ornithinicoccus* | 7.59 | 12.13 | 9.91 | 7.48 | 12.41 | 11.71 | -0.09 | 0.764 |
| *Melampsora* | 4.99 | 9.06 | 3.86 | 15.11 | 14.94 | 11.45 | -1.21 | 0.019 |
| *Armillaria* | 4.99 | 11.62 | 6.38 | 9.97 | 8.36 | 11.45 | -0.37 | 0.387 |
| *Labrenzia* | 8.46 | 11.45 | 15.45 | 9.19 | 9.62 | 11.45 | 0.23 | 0.495 |
| *Globisporangium* | 26.45 | 12.47 | 6.38 | 10.59 | 15.95 | 11.45 | 0.26 | 0.725 |
| *Pseudochrobactrum* | 16.91 | 26.82 | 36.96 | 20.72 | 8.10 | 11.19 | 1.01 | 0.132 |
| *Plantibacter* | 29.92 | 20.16 | 52.24 | 28.51 | 6.08 | 11.19 | 1.16 | 0.189 |
| *Actinorugispora* | 10.62 | 30.24 | 14.28 | 14.80 | 11.14 | 11.19 | 0.57 | 0.425 |
| *Lapillicoccus* | 9.76 | 5.64 | 9.57 | 6.54 | 9.62 | 11.19 | -0.13 | 0.701 |
| *Porphyridium* | 6.72 | 8.54 | 5.71 | 8.26 | 10.38 | 10.93 | -0.49 | 0.069 |
| *Agreia* | 15.83 | 15.55 | 33.93 | 21.81 | 5.06 | 10.93 | 0.79 | 0.308 |
| *Niveispirillum* | 5.20 | 6.83 | 9.57 | 6.39 | 11.39 | 10.93 | -0.41 | 0.314 |
| *Dechloromonas* | 15.18 | 13.67 | 11.93 | 15.27 | 13.42 | 10.93 | 0.04 | 0.818 |
| *Pararhodobacter* | 9.54 | 10.25 | 12.93 | 9.19 | 7.85 | 10.67 | 0.24 | 0.273 |
| *Trypanosoma* | 12.14 | 6.66 | 6.38 | 9.50 | 22.03 | 10.67 | -0.74 | 0.294 |
| *Fictibacillus* | 35.12 | 1075.88 | 2902.01 | 465.50 | 8.86 | 10.67 | 3.05 | 0.294 |
| *Plesiomonas* | 5.85 | 13.67 | 4.70 | 14.49 | 10.38 | 10.67 | -0.55 | 0.317 |
| *Clavibacter* | 16.26 | 16.40 | 41.99 | 25.08 | 10.89 | 10.67 | 0.68 | 0.408 |
| *Pythium* | 64.18 | 12.47 | 11.25 | 13.71 | 14.43 | 10.67 | 1.18 | 0.447 |
| *Fulvimarina* | 10.19 | 10.93 | 21.67 | 13.40 | 8.10 | 10.67 | 0.41 | 0.449 |
| *Pelagerythrobacter* | 9.97 | 18.45 | 27.88 | 21.03 | 8.86 | 10.67 | 0.47 | 0.463 |
| *Pelomyxa* | 13.23 | 17.94 | 8.23 | 11.84 | 10.63 | 10.67 | 0.25 | 0.535 |
| *Lutibaculum* | 6.50 | 6.49 | 12.77 | 6.54 | 10.63 | 10.67 | -0.11 | 0.798 |
| *Caldalkalibacillus* | 9.54 | 151.21 | 86.34 | 16.83 | 6.58 | 10.41 | 2.87 | 0.224 |
| *Dermacoccus* | 6.94 | 11.11 | 8.06 | 170.74 | 8.10 | 10.41 | -2.86 | 0.419 |
| *Porphyrobacter* | 7.15 | 14.01 | 17.13 | 11.68 | 9.12 | 10.41 | 0.30 | 0.509 |
| *Mucor* | 6.72 | 19.48 | 14.45 | 14.64 | 7.60 | 10.41 | 0.32 | 0.572 |
| *Methylotenera* | 20.81 | 5.81 | 10.58 | 9.19 | 11.14 | 10.41 | 0.28 | 0.675 |
| *Polymorphospora* | 3.69 | 13.16 | 18.98 | 24.15 | 6.33 | 10.41 | -0.19 | 0.821 |
| *Rufibacter* | 5.64 | 10.25 | 12.77 | 4.83 | 12.15 | 10.41 | 0.07 | 0.895 |
| *Pelagibius* | 9.76 | 9.40 | 13.94 | 7.17 | 14.94 | 10.41 | 0.03 | 0.944 |
| *Diacronema* | 12.36 | 11.62 | 7.73 | 9.35 | 12.15 | 10.41 | -0.01 | 0.973 |
| *Methylobacillus* | 15.83 | 5.81 | 14.11 | 14.80 | 10.63 | 10.41 | 0.00 | 0.994 |
| *Melghirimyces* | 5.42 | 34.85 | 41.99 | 11.53 | 7.34 | 10.15 | 1.50 | 0.253 |
| *Labedella* | 18.00 | 20.67 | 47.03 | 32.40 | 6.58 | 10.15 | 0.80 | 0.379 |
| *Homoserinimonas* | 12.14 | 11.28 | 22.34 | 18.85 | 8.61 | 10.15 | 0.28 | 0.600 |
| *Mangrovicella* | 7.81 | 12.13 | 12.43 | 12.62 | 5.82 | 10.15 | 0.18 | 0.641 |
| *Roseobacter* | 6.29 | 10.08 | 13.77 | 7.32 | 9.12 | 10.15 | 0.18 | 0.647 |
| *Pseudorhodobacter* | 6.29 | 6.49 | 18.14 | 10.13 | 8.86 | 10.15 | 0.09 | 0.893 |
| *Arsenicicoccus* | 8.67 | 11.79 | 9.41 | 6.70 | 8.10 | 9.89 | 0.28 | 0.257 |
| *Siculibacillus* | 8.89 | 10.42 | 11.09 | 3.74 | 8.61 | 9.89 | 0.45 | 0.279 |
| *Tribonema* | 12.36 | 11.96 | 6.72 | 7.01 | 9.37 | 9.89 | 0.24 | 0.489 |
| *Roseicella* | 8.02 | 8.54 | 8.73 | 6.39 | 12.15 | 9.89 | -0.17 | 0.599 |
| *Myceligenerans* | 6.72 | 7.86 | 12.26 | 9.66 | 8.36 | 9.89 | -0.06 | 0.858 |
| *Paecilomyces* | 9.97 | 9.06 | 16.13 | 15.58 | 8.10 | 9.89 | 0.07 | 0.875 |
| *Sinirhodobacter* | 8.89 | 10.08 | 16.29 | 7.63 | 6.33 | 9.63 | 0.58 | 0.225 |
| *Geobacillus* | 10.62 | 57.75 | 28.39 | 12.77 | 5.32 | 9.63 | 1.80 | 0.234 |
| *Naegleria* | 15.39 | 11.45 | 10.41 | 13.55 | 9.62 | 9.63 | 0.18 | 0.502 |
| *Saccharibacillus* | 13.23 | 23.41 | 16.97 | 12.77 | 7.34 | 9.37 | 0.86 | 0.095 |
| *Stenotrophobium* | 59.41 | 12.81 | 45.19 | 27.26 | 9.12 | 9.37 | 1.36 | 0.219 |
| *Naasia* | 28.19 | 31.78 | 86.34 | 53.12 | 6.84 | 9.37 | 1.08 | 0.350 |
| *Xylophilus* | 9.76 | 7.52 | 12.09 | 19.94 | 11.65 | 9.37 | -0.48 | 0.358 |
| *Alteromonas* | 8.24 | 9.23 | 10.41 | 6.70 | 10.13 | 9.37 | 0.09 | 0.668 |
| *Aureobasidium* | 3.90 | 13.67 | 9.74 | 10.28 | 10.38 | 9.37 | -0.14 | 0.781 |
| *Komagataeibacter* | 6.50 | 7.86 | 11.25 | 6.08 | 9.37 | 9.37 | 0.05 | 0.886 |
| *Paramesorhizobium* | 9.76 | 9.74 | 19.15 | 7.17 | 6.33 | 9.11 | 0.77 | 0.225 |
| *Diaminobutyricibacter* | 117.08 | 89.19 | 259.19 | 168.72 | 10.38 | 9.11 | 1.30 | 0.284 |
| *Methylibium* | 7.81 | 14.86 | 53.92 | 14.49 | 7.09 | 9.11 | 1.32 | 0.398 |
| *Rothia* | 6.07 | 10.08 | 13.61 | 8.26 | 5.57 | 9.11 | 0.38 | 0.418 |
| *Ahrensia* | 7.37 | 6.66 | 14.61 | 7.95 | 8.36 | 9.11 | 0.17 | 0.712 |
| *Xylanibacillus* | 14.53 | 20.16 | 30.91 | 14.96 | 4.05 | 8.85 | 1.24 | 0.104 |
| *Oleiagrimonas* | 7.37 | 41.69 | 32.25 | 30.53 | 19.24 | 8.85 | 0.47 | 0.570 |
| *Chitinimonas* | 31.44 | 3.42 | 3.36 | 3.89 | 7.09 | 8.85 | 0.95 | 0.580 |
| *Parasphingopyxis* | 7.15 | 6.83 | 10.92 | 7.95 | 10.89 | 8.85 | -0.15 | 0.594 |
| *Raineyella* | 7.59 | 7.69 | 10.75 | 8.72 | 7.09 | 8.85 | 0.08 | 0.722 |
| *Vitrella* | 12.14 | 6.32 | 6.22 | 7.48 | 10.89 | 8.85 | -0.14 | 0.729 |
| *Sphaerobolus* | 6.07 | 14.35 | 6.55 | 15.58 | 13.17 | 8.59 | -0.47 | 0.369 |
| *Spelaeicoccus* | 6.29 | 12.81 | 12.77 | 9.66 | 8.10 | 8.59 | 0.27 | 0.486 |
| *Glycocaulis* | 8.89 | 13.67 | 8.40 | 13.55 | 13.93 | 8.59 | -0.22 | 0.519 |
| *Cytobacillus* | 6.94 | 102.68 | 231.14 | 45.33 | 5.57 | 8.33 | 2.52 | 0.284 |

| | | | | | | | | |
|---|---|---|---|---|---|---|---|---|
| *Ohtaekwangia* | 103.85 | 4.61 | 3.53 | 1.40 | 3.80 | 8.33 | 3.05 | 0.428 |
| *Robertmurraya* | 9.54 | 29.39 | 18.14 | 25.55 | 15.45 | 8.33 | 0.21 | 0.752 |
| *Natronosporangium* | 4.55 | 13.67 | 18.81 | 12.93 | 8.10 | 8.07 | 0.35 | 0.602 |
| *Runella* | 5.20 | 5.13 | 27.55 | 9.66 | 8.36 | 8.07 | 0.54 | 0.651 |
| *Jannaschia* | 8.02 | 9.91 | 16.29 | 8.26 | 13.67 | 8.07 | 0.19 | 0.674 |
| *Emiliania* | 14.09 | 9.74 | 8.23 | 9.81 | 12.15 | 8.07 | 0.10 | 0.766 |
| *Salpingoeca* | 11.49 | 9.91 | 4.70 | 8.41 | 7.85 | 8.07 | 0.10 | 0.799 |
| *Azospira* | 33.61 | 5.98 | 7.73 | 3.58 | 10.38 | 7.81 | 1.12 | 0.442 |
| *Pelobium* | 9.97 | 9.23 | 15.12 | 10.28 | 11.90 | 7.81 | 0.19 | 0.552 |
| *Moraxella* | 2.38 | 3.59 | 27.88 | 6.23 | 4.56 | 7.81 | 0.86 | 0.603 |
| *Gulosibacter* | 4.34 | 7.35 | 15.79 | 8.88 | 6.84 | 7.81 | 0.22 | 0.739 |
| *Xinfangfangia* | 14.96 | 30.92 | 41.32 | 20.25 | 6.08 | 7.55 | 1.36 | 0.132 |
| *Lipomyces* | 7.15 | 10.59 | 11.59 | 8.26 | 6.33 | 7.55 | 0.41 | 0.207 |
| *Diaminobutyricimonas* | 15.61 | 12.13 | 29.06 | 18.69 | 6.08 | 7.55 | 0.81 | 0.283 |
| *Maritalea* | 6.72 | 8.71 | 13.44 | 5.14 | 8.61 | 7.55 | 0.44 | 0.340 |
| *Capillibacterium* | 5.85 | 17.77 | 268.77 | 16.05 | 4.81 | 7.55 | 3.36 | 0.413 |
| *Eoetvoesia* | 7.15 | 45.28 | 75.42 | 59.20 | 6.84 | 7.55 | 0.80 | 0.531 |
| *Suillus* | 9.76 | 15.38 | 7.06 | 17.45 | 11.65 | 7.55 | -0.19 | 0.716 |
| *Compostimonas* | 14.31 | 12.47 | 25.20 | 17.76 | 4.05 | 7.29 | 0.84 | 0.255 |
| *Anoxybacillus* | 7.81 | 67.32 | 28.89 | 16.20 | 4.56 | 7.29 | 1.89 | 0.281 |
| *Pseudorhodoferax* | 16.69 | 6.66 | 9.91 | 8.10 | 7.60 | 7.29 | 0.53 | 0.366 |
| *Acetivibrio* | 3.25 | 11.96 | 46.36 | 5.92 | 6.08 | 7.29 | 1.67 | 0.396 |
| *Tropicimonas* | 4.34 | 8.88 | 17.81 | 8.57 | 6.33 | 7.29 | 0.48 | 0.535 |
| *Chengkuizengella* | 7.37 | 16.91 | 17.81 | 9.04 | 3.29 | 6.76 | 1.14 | 0.134 |
| *Naumannella* | 7.15 | 12.13 | 13.94 | 7.32 | 7.09 | 6.76 | 0.65 | 0.185 |
| *Celeribacter* | 7.59 | 9.40 | 13.10 | 7.95 | 6.58 | 6.76 | 0.50 | 0.206 |
| *Caldibacillus* | 6.50 | 122.50 | 44.68 | 9.81 | 3.04 | 6.76 | 3.15 | 0.271 |
| *Erwinia* | 13.01 | 4.95 | 12.26 | 6.85 | 7.85 | 6.76 | 0.49 | 0.373 |
| *Marisediminicola* | 8.89 | 5.47 | 15.12 | 7.17 | 7.85 | 6.76 | 0.44 | 0.459 |
| *Puniceibacterium* | 4.34 | 11.79 | 14.28 | 11.68 | 4.81 | 6.76 | 0.39 | 0.551 |
| *Peteryoungia* | 11.27 | 28.02 | 33.60 | 23.21 | 3.80 | 6.50 | 1.12 | 0.221 |
| *Exiguobacterium* | 5.85 | 21.70 | 12.93 | 6.70 | 3.80 | 6.50 | 1.25 | 0.226 |
| *Haemophilus* | 17.56 | 7.35 | 9.74 | 8.88 | 3.80 | 6.50 | 0.85 | 0.233 |
| *Mesobacillus* | 7.59 | 71.76 | 44.01 | 38.32 | 4.56 | 6.50 | 1.32 | 0.331 |
| *Dictyostelium* | 9.11 | 12.99 | 10.41 | 11.84 | 8.61 | 6.50 | 0.27 | 0.396 |
| *Galdieria* | 10.62 | 7.18 | 5.04 | 12.31 | 8.61 | 6.50 | -0.26 | 0.554 |
| *Ascochyta* | 3.47 | 18.62 | 25.36 | 0.62 | 3.29 | 6.24 | 2.22 | 0.189 |
| *Thermoflavimicrobium* | 3.47 | 17.26 | 30.91 | 10.28 | 5.06 | 6.24 | 1.26 | 0.333 |
| *Sarocladium* | 0.87 | 35.88 | 0.67 | 10.28 | 0.76 | 6.24 | 1.11 | 0.628 |
| *Cutibacterium* | 7.37 | 9.57 | 9.57 | 12.15 | 10.38 | 6.24 | -0.12 | 0.722 |
| *Angulomicrobium* | 9.76 | 20.67 | 25.70 | 18.23 | 11.65 | 5.98 | 0.65 | 0.320 |
| *Monosporascus* | 6.94 | 19.99 | 11.25 | 16.98 | 4.30 | 5.98 | 0.49 | 0.547 |
| *Anaerobacillus* | 3.69 | 35.71 | 18.48 | 9.19 | 2.53 | 5.72 | 1.73 | 0.281 |
| *Endobacterium* | 8.02 | 15.04 | 27.04 | 16.51 | 5.82 | 5.72 | 0.84 | 0.338 |
| *Tersicoccus* | 5.42 | 7.86 | 17.81 | 13.09 | 5.57 | 5.72 | 0.35 | 0.652 |
| *Ornithinibacillus* | 6.29 | 131.90 | 74.75 | 13.24 | 2.28 | 5.46 | 3.34 | 0.219 |
| *Lentibacillus* | 4.55 | 124.55 | 21.33 | 6.23 | 4.81 | 5.46 | 3.19 | 0.356 |
| *Fontibacillus* | 5.85 | 35.54 | 12.26 | 13.09 | 5.57 | 5.46 | 1.15 | 0.391 |
| *Yersinia* | 17.56 | 3.25 | 9.07 | 8.88 | 6.08 | 5.46 | 0.55 | 0.531 |
| *Conyzicola* | 7.59 | 6.32 | 14.78 | 15.89 | 4.05 | 5.46 | 0.17 | 0.824 |
| *Galbitalea* | 8.24 | 7.52 | 15.96 | 7.95 | 5.32 | 5.20 | 0.78 | 0.238 |
| *Lihuaxuella* | 1.73 | 10.76 | 23.18 | 7.01 | 2.53 | 5.20 | 1.27 | 0.378 |
| *Tissierella* | 3.25 | 15.55 | 13.27 | 8.88 | 6.84 | 5.20 | 0.62 | 0.432 |
| *Rhodocytophaga* | 3.90 | 17.60 | 9.57 | 5.61 | 9.37 | 5.20 | 0.62 | 0.462 |
| *Fulvimonas* | 6.07 | 46.30 | 36.12 | 35.52 | 24.81 | 5.20 | 0.43 | 0.639 |
| *Schnuerera* | 3.90 | 53.99 | 34.77 | 8.26 | 2.28 | 4.94 | 2.58 | 0.219 |
| *Fibrella* | 3.04 | 4.61 | 34.94 | 4.83 | 5.82 | 4.94 | 1.45 | 0.477 |
| *Ureibacillus* | 3.25 | 99.78 | 27.55 | 7.48 | 4.81 | 4.68 | 2.94 | 0.321 |
| *Ectobacillus* | 20.81 | 31.95 | 46.70 | 47.98 | 6.08 | 4.68 | 0.76 | 0.461 |
| *Limnochorda* | 2.38 | 7.86 | 62.99 | 12.62 | 4.30 | 4.68 | 1.76 | 0.468 |
| *Psychrobacillus* | 3.69 | 28.36 | 37.12 | 6.54 | 2.03 | 4.42 | 2.41 | 0.200 |
| *Halobacillus* | 7.15 | 69.20 | 21.50 | 12.00 | 4.81 | 4.42 | 2.20 | 0.306 |
| *Puia* | 4.12 | 42.03 | 15.29 | 41.28 | 5.82 | 4.42 | 0.25 | 0.852 |
| *Sporolactobacillus* | 5.85 | 29.90 | 13.77 | 5.30 | 4.56 | 4.16 | 1.82 | 0.236 |
| *Planococcus* | 6.07 | 32.63 | 13.44 | 5.92 | 5.06 | 4.16 | 1.78 | 0.259 |
| *Shimazuella* | 3.69 | 12.64 | 31.92 | 73.07 | 3.04 | 4.16 | -0.74 | 0.698 |
| *Domibacillus* | 5.64 | 29.22 | 15.96 | 11.84 | 3.04 | 3.90 | 1.44 | 0.255 |
| *Tepidanaerobacter* | 0.43 | 12.30 | 58.12 | 5.76 | 2.03 | 3.90 | 2.60 | 0.379 |
| *Pseudogulbenkiania* | 45.75 | 2.22 | 3.53 | 1.71 | 3.80 | 3.90 | 2.45 | 0.430 |
| *Pullulanibacillus* | 12.36 | 25.12 | 15.79 | 9.81 | 6.84 | 3.64 | 1.39 | 0.084 |
| *Parageobacillus* | 3.47 | 20.16 | 17.30 | 4.99 | 1.27 | 3.64 | 2.05 | 0.178 |
| *Fredinandcohnia* | 6.94 | 40.83 | 82.14 | 17.92 | 3.04 | 3.64 | 2.40 | 0.245 |
| *Pueribacillus* | 3.25 | 595.60 | 32.92 | 7.48 | 3.04 | 3.64 | 5.48 | 0.397 |
| *Symbiobacterium* | 3.90 | 12.99 | 53.42 | 43.15 | 3.04 | 3.64 | 0.49 | 0.754 |
| *Fibrisoma* | 3.25 | 4.78 | 46.87 | 7.95 | 3.29 | 3.38 | 1.91 | 0.447 |
| *Polycladomyces* | 3.47 | 23.07 | 25.87 | 7.95 | 2.53 | 3.12 | 1.95 | 0.203 |
| *Pontibacillus* | 2.82 | 35.71 | 10.92 | 4.67 | 2.79 | 3.12 | 2.22 | 0.321 |
| *Pseudogracilibacillus* | 3.69 | 535.80 | 53.42 | 12.77 | 2.79 | 3.12 | 4.99 | 0.376 |
| *Caldicoprobacter* | 1.73 | 12.47 | 15.62 | 79.30 | 2.79 | 3.12 | -1.52 | 0.545 |
| *Compostibacillus* | 12.14 | 112.08 | 69.21 | 7.63 | 3.04 | 2.86 | 3.84 | 0.174 |
| *Candidatus* Nitrosacidococcus | 40.33 | 2.22 | 10.75 | 9.35 | 3.29 | 2.86 | 1.78 | 0.390 |
| *Diplodia* | 35.99 | 1.20 | 5.54 | 4.21 | 1.77 | 2.60 | 2.32 | 0.408 |
| *Telluribacter* | 1.30 | 18.45 | 15.79 | 77.89 | 2.03 | 2.34 | -1.21 | 0.603 |
| *Aquibacillus* | 3.25 | 36.39 | 12.93 | 6.39 | 1.77 | 1.82 | 2.40 | 0.285 |
| *Amphibacillus* | 2.17 | 34.51 | 9.74 | 4.36 | 2.79 | 1.56 | 2.41 | 0.327 |
| *Siminovitchia* | 1.73 | 34.51 | 11.25 | 6.54 | 2.03 | 1.56 | 2.23 | 0.329 |
| *Cerasibacillus* | 0.65 | 77.06 | 8.90 | 3.43 | 0.76 | 1.56 | 3.91 | 0.381 |
| *Sutcliffiella* | 1.73 | 21.87 | 20.16 | 5.76 | 1.01 | 1.30 | 2.44 | 0.201 |
| *Arsenicibacter* | 1.95 | 22.72 | 194.35 | 23.99 | 1.52 | 1.04 | 3.04 | 0.403 |
| *Baia* | 0.87 | 3.25 | 49.72 | 0.93 | 0.76 | 0.78 | 4.44 | 0.394 |
| *Vermamoeba* | 0.22 | 30.07 | 13.27 | 32.40 | 0.51 | 0.52 | 0.38 | 0.819 |
| *Tepidimicrobium* | 0.65 | 112.76 | 23.85 | 2.18 | 0.51 | 0.00 | 5.67 | 0.320 |

RPM, reads per million.
