## Supplementary material for "Horizontally acquired *IbACS* gene modulates rhizosphere microbiota and contributes to sweet potato growth": Table S3

**Table S3**. PCR primers used in this study.

| Primer name | Sequence | Note |
|---|---|---|
| acs_cc3_F2 | TCGAAGTAGTGATTGTTACAAGTTATCGAGCACAGGTTTTAGAGCTAGAA | Vector construction target 3 |
| acs_cc3_R2 | TTCTAGCTCTAAAACCTGTGCTCGATAACTTGTAACAATCACTACTTCGA | Vector construction target 3 |
| acs_cc4_F2 | TCGAAGTAGTGATTGACAGAGTTCCTCAGGAAAAAGTTTTAGAGCTAGAA | Vector construction target 1 |
| acs_cc4_R2 | TTCTAGCTCTAAAACTTTTTCCTGAGGAACTCTGTCAATCACTACTTCGA | Vector construction target 1 |
| acs_cc5_F | TCGAAGTAGTGATTGGTTCCTTCCATCCAGAAAGAGTTTTAGAGCTAGAA | Vector construction target 4 |
| acs_cc5_R | TTCTAGCTCTAAAACTCTTTCTGGATGGAAGGAACCAATCACTACTTCGA | Vector construction target 4 |
| acs_cc6_F | TCGAAGTAGTGATTGGATGCCACCGTCTTTCTGGAGTTTTAGAGCTAGAA | Vector construction target 2 |
| acs_cc6_R | TTCTAGCTCTAAAACTCCAGAAAGACGGTGGCATCCAATCACTACTTCGA | Vector construction target 2 |
| AtU6-26P_H | TAGGCGCGCCAAGCTTCGACTTGCCTTCCGCACAA | Vector construction |
| AtU6-26T_H | GCAGGCATGCAAGCTTATTGGTTTATCTCATCGGA | Vector construction |
| AtU6-26P_Asc | ATGTTACTAGGCGCGCCAAGCTTCGACTTGCCTTC | Vector construction |
| AtU6-26T_Mlu | TCGAAGCTTGGCGCGTCATGCAAGCTTATTGGTTT | Vector construction |
| Acs.F3 | GCAGCCCGGGGGATCCATGCTTTGCCAGCGAAAACCTAG | Verification of mutations in KO lines |
| Acs.R3 | CCGCTCTAGAACTAGTGAGAGTACGGAATCAATCC | Verification of mutations in KO lines |
